# RoSe-BaL: A neuroanatomically plausible model of routine action sequencing

**DOI:** 10.1101/2025.03.06.641869

**Authors:** Jen Lewis, Kevin Gurney

**Affiliations:** SCHARR, Division of Population Health, University of Sheffield, UK; Department of Psychology, University of Sheffield, UK

## Abstract

The performance of routine action sequences constitutes a significant proportion of human behaviour, and has received much attention in the cognitive psychology literature. However, a neuroanatomically plausible explanation of the cognitive processes underlying this routine sequential behaviour has hitherto remained elusive. This is despite wide acceptance that an established hierarchy of interconnected basal-ganglia thalamocortical (BGTC) loops appear to be heavily involved in organising and executing context-sensitive routine action.

Here, we build on existing computational models of action selection in the basal ganglia to develop the ‘Routine-Sequencing Basal-ganglia Loops’ (RoSe-BaL) model of multiple basal-ganglia thalamocortical loops occupying associative and motor territories of the brain. We demonstrate this model in the sequential routine tasks of tea– and coffee-making. This model incorporates a novel hypothesis of the nature of the organisation of cognitive information in associative territories of the BGTC hierarchy, which we term the ‘BG subset-selection hypothesis’. Erroneous behaviour made by the model under conditions of disruption shows similar trends to those observed in previous studies of action slips and action disorganisation syndrome. This finding provides support for the previously untested hypothesis that damaged temporal order knowledge and action schemas underlie many of the errors typically performed by humans. The neuroanatomical grounding of the model naturally reconciles two influential competing computational models of routine behaviour. Finally, we propose a novel interpretation of the widespread competitive queueing account, which we term ‘suitability queueing’. This is shown to be consistent with existing cognitive, neurophysiological and neuroanatomical findings.

## 1. Introduction

‘Routine’ action sequences are those that are familiar and well learned to the point of some automaticity. Often, they have a degree of flexibility in the detail of their execution and temporal ordering, though they are largely stereotypical, taking roughly the same form with each performance. Dynamic environmental factors often demand alterations in performance, such as when interruptions occur. The ability to vary the precise order and detail of a sequence of actions is therefore critical for successful routine action. Routine action requires the retrieval from long-term memory of the required steps for task completion (1) and their temporal order; imposing particular demands upon storage and retrieval processes, as well as ordering.

Given this combination of automaticity and flexibility, routine action seems to breach the divide between truly automatic (‘habit’), and goal-directed behaviour (2). Though routine action is often considered habitual, many of its features suggest otherwise. For example, such actions – like tooth-brushing or tea-making – are rarely evoked simply by the presence of a related stimulus, often considered one of the defining features of a habit (3). Furthermore, the performance of routine action appears to rely heavily on prefrontal regions of the brain (4–8), the involvement of which has historically not been considered to be important for habitual behaviour (9, 10). Moreover, a particular goal is usually present during routine behaviour (2, 11, 12). Indeed, the very need for flexibility and variation may prevents these routines from becoming completely habitual. Here, we investigate routine action using a neuroanatomically constrained model of action selection and sequencing. In doing so, we bring together features from the cognitive psychological and neurophysiology literatures to give a neuroscientifically grounded account of many experimentally observed features of routine action.

### 1.1. Cognitive perspectives on routine action

The part-habit, part-goal oriented nature of routine action makes the nature of its underlying cognitive representation particularly difficult to discern. What seems clear is that successful execution of complex action sequences is reliant on *context*: accounts such as chaining, that have attempted to explain sequential processing in the absence of its representation have proved problematic (13). Typically, psychological explanations have relied on the concept of action ‘schemas’ to understand the mediation of routine actions. Schemas are cognitive structures representing a set of features, including context, required for the direction and performance of a sequence, subsequence, or action. Schemas are proposed to exist at several levels of description, with higher level or ‘intention’ schemas activating lower level or ‘child’ schemas (14–17), with routine behaviour being evoked by the sequential activation of discrete action schemas. The proposed hierarchy of schemas is consistent with the popular notion that action itself is hierarchically organised (18–21). This makes intuitively sense; composite actions can be put together in various orders to accomplish short-term tasks and subgoals, which themselves can be used to work towards higher order goals. Thus, higher levels of an action hierarchy account for more temporally extended segments of action, and increasingly distal levels of structure. Routine action is then likely to require representation at multiple levels of a hierarchy (2).

#### Competitive queuing + contention scheduling

Support for the notion of discernible cognitive representations for action which may be interpreted as schemas came from a study in which monkeys were trained to draw geometric shapes, like triangles or rectangles, in response to a stimulus (22). Prefrontal recordings taken during drawing revealed a set of ‘ensembles’ of prefrontal neurons, each of which showed activity that correlated with the drawing of a particular segment of the shapes. Notably, these ensembles’ activities peaked during the drawing of their corresponding segment, but all showed some activation during a preparatory phase. This study has been cited in support of the ‘competitive queueing’ account of sequential behaviour (23, 24). In this account, representations of actions required for a sequence are activated in parallel at the beginning of that sequence. This activation is proportional to their temporal distance from the beginning of the sequence, resulting in an activation ‘gradient’ across representations. As each action is executed, its corresponding representation is inhibited, allowing the next to become selected. This account, while popular, does not naturally take dynamic contextual information into account, with the order of actions being strictly defined from the sequence outset. Moreover, the nature of the required sequencing mechanism would struggle with tasks requiring more than two hierarchical levels of action (25) and those with repeated actions (23).

In the similar ‘contention scheduling’ account (26), actions are executed when their corresponding schemas are activated beyond some threshold. However, rather than obeying a predetermined order, sequencing occurs through the satisfaction of pre– and post-conditions for the activation of schemas. This account allows for environmental or bottom-up influences on action, and affords a greater degree of flexibility than the competitive queueing account.

#### Cognitive modelling

The contention scheduling account was implemented in a computational model known as the ‘interactive activation network’ (16), which performed an abstracted coffee-making sequence. The model comprised a hierarchical network of abstract action schemas. Schemas were implemented as localist nodes, each with a dynamic activation value, and was selected once its activation exceeded a predetermined threshold. Selection of a schema at a high level of the hierarchy corresponded to goal selection (e.g., ‘prepare instant coffee’) and activated lower-level schemas, representing subgoal selection (e.g., ‘add coffee from jar’). The lowest level schemas represented discrete actions, the selection of which drove execution of the action by effectors. By varying the relative influences of top-down and bottom-up information, the model accounted for a variety of common patterns of human error while performing routine action. These included ‘action slips’ (15) and more severely disrupted performance in patients with action disorganisation syndrome (ADS) (5, 27). This provided support for a hypothesis that ADS resulted from an imbalance of bottom-up and top-down influence on action (28). This model provided strong initial support for the contention scheduling approach to understanding the cognitive dynamics underlying routine behaviour, and for taking a schema-oriented view to understanding performance error. Later adaptations of the model accounted for error patterns in multiple disorders of action by selectively disrupting various parameters (29).

A contrasting model of routine action developed by Botvinick and Plaut (25) took a different approach. The authors rejected the necessity for schemas, suggesting that the explicit hierarchical structure and localist representation scheme in the IAN lacked flexibility and generalisability. They proposed a three-layer simple recurrent network (SRN) model which utilised backpropagation to learn mappings between environmental inputs and action outputs. Recurrent connectivity maintained a dynamic record of its activation history – i.e., a record of the temporal task context – thus making the mappings context-dependent. This flexibility afforded by this context-dependency was emphasised as a critical distinction from explicitly hierarchical models. The SRN model was also able to replicate a range of data from observational studies on human error. Here however, this resulted from the noise in hidden layer nodes, consistent with an alternative hypothesis that ADS results from a general reduction in cognitive resources (5, 30). Consistent with observational studies, the model showed a general decline in performance with increasing noise, and replicated the ‘omission rate effect’, documented by several authors (5, 31). The model was able to perform a ‘quasi-hierarchical task’ with multiple variations, which, the authors claimed, demonstrated greater efficiency and flexibility in the underlying representations than is possible with a strict hierarchical structure. However, certain errors commonly reported in the preceding neuropsychological studies were extremely rare (32), suggesting that more specific deficits may be responsible for some errors in ADS.

Both these models provided important insights into the potential functional underpinnings of routine sequence production, accounting for many error patterns observed in behavioural studies, with implications for understanding the structure and dynamics of cognitive representations. However, neither is constrained from a biological perspective. While a recent model embeds the IAN model in a neuroanatomically inspired architecture (33), this model does not deal with sequential behaviour nor the types of errors encountered in ADS, thus the question remains whether either approach might reasonably be implemented neurally and retain the ability to reproduce ADS error profiles.

### 1.2. Neural substrates for action

In the neurobehavioural literature, routine action is naturally embraced within the wider context of the study of ‘action selection’ – the problem of choosing what to do next. In vertebrates, the basal ganglia (BG) are widely believed to be essential for the solution of this problem (34–36). The BG are arranged in topographically organised loops with thalamus and cortex (37). These ‘basal ganglia thalamo-cortical’ (BGTC) loops are present through multiple regions of the brain. This series of distinct circuits appears to not only select actions, but select information in several functional domains, including motor, cognitive and limbic (38). This group of subcortical nuclei display several structural and functional features suggesting their probable involvement in selection, including their wide-ranging connectivity with other neural structures, topographical organisation, and intrinsic functional architecture and internal dynamics (35). These allow a selective release of tonic inhibition on thalamocortical targets, which in turn allows the activation of effectors to execute the selected action, or corresponding equivalent in cognitive and limbic territories (35). The internal circuitry of the BG comprises multiple overlapping and recurrent pathways, and its resulting non-linearity enables it to perform more complex functions than simple gating, including contrast enhancement, resolution of competition, selective and general inhibition and signal normalisation (39–41).

#### Context for action

PFC is likely to play a prominent role in the cognitive control of action selection. The role of PFC and related functional areas in cognitive processing has been well documented (1, 2, 42–47). Activity in PFC is consistently reported to be sensitive to goals (47, 48), action plans (49, 50), context-specific action (51–53), sequence identity (22, 54), sequence progress tracking (55–57) and sequence category (58). Moreover, PFC neurons have been shown to be selective for, or modulated by, stimulus features which are relevant for guiding action, such as object identity, shape, colour and location, as well as more abstract information such as expected reward (45, 47). PFC is also recognised as an important substrate of working memory through sustained activation (42, 59, 60). Furthermore, sequential tasks are known to be affected by frontal lobe damage in humans (61). This and other evidence points to the importance of PFC in selecting and maintaining in working memory a representation of the task *context* (62, 63): information that is critical for guiding action through task sequences. Moreover, corresponding associative regions of BG have been implicated in planning for sequencing (64, 65), and like PFC, BG neurons show sensitivity to context and are involved in working memory (66, 67).

It is the structure and mediation of this cognitive information often linked with PFC that the IAN and SRN models focused on, and did not reach a consensus on the nature of its processing (32, 68–70). In this paper, we reframe this problem by constraining the possible solutions according to the known neuroanatomy and evidence from neurophysiology. We propose a novel hypothesis regarding the interactions of PFC and BG, and implement this in a series of neurobiologically grounded models demonstrating the ultimate compatibility of the previous, cognitive level approaches given in the IAN and SRN models. The model uses two BG loops – associative and motor – working at different levels of action description, and shows how these descriptions can be anatomically and conceptually integrated. We now outline some foundational computational and anatomical constraints for our programme of work.

### 1.3. Computational considerations

Any internal representation of information that is important for correctly selecting motor action must adhere to a set of basic criteria that should constrain any hypothesis of cognitive selection in neural circuits.

1. Uniqueness Representations of temporal task context must be unique. Temporal context includes (but may not be limited to) the information necessary to drive correct selection of the current action, and determine the next. This is consistent with the notion that PFC maintains a representation of contextual features (45, 47, 52), possibly in the form of some schema (2, 20, 71, 72). Uniqueness means that downstream influences of contextual representations are unambiguous, and must evolve to remain up to date with the ongoing task.
2. Sustainability Representations of context must be sustainable by the underlying substrate. Sustained activity of ensembles of PFC neurons encoding stimulus and task features is believed to underlie working memory (42, 45, 59). A plausible model of the associative BGTC loop should include this functionality by autonomously maintaining activation of selected representations of context.
3. Efficiency Information comprising context must be represented efficiently. The impracticality and implausibility of a ‘localist’ coding scheme for cognitive (but not motor) information (22, 25) tells us it is likely that complex, multidimensional contextual information is encoded efficiently at the neural level, with the re-use of individual coding elements across multiple representations (53).

### 1.4. Anatomical considerations

Selection of task representations and updating of working memory is believed to rely on prefrontal BGTC loops (63, 73, 74), but the functional anatomy supporting this is not well understood. In motor territories of BGTC circuitry, the relationship between cortex and BG is defined: motor cortices are arranged topographically (75, 76), and this organisation is largely preserved throughout BG (77). This results, in essence, in a localist coding scheme throughout motor territories, reflected in the channel-wise organisation of several BG models, including the ‘GPR’ model of selection in BG (39, 40). In contrast, although the precise structure, dynamics and encoding of the information within PFC remains unclear (53, 78–82), PFC representations of complex stimuli and task progress information are almost certainly at least partly distributed (47). The strong functional topography of the relationship between cortex and BG in the motor loop cannot then be maintained in the associative loop, since the competitive selection mechanisms therein appear to rely on a closed ‘micro-loop’ architecture requiring a localist code (35, 39).

### 1.5. The subset-selection hypothesis

Computational and anatomical considerations suggest a conflict between the implausibility of a localist code for complex representations, but the necessity of such a code for selection by BG. Thus, some information compression must be occurring in BG. This is consistent with the degree of converging projections from cortex (34) suggesting that compression or dimensionality reduction is likely (83). BG may therefore be performing selection in a *coarse-grained* manner at this level of the BGTC system. By this, we mean that each channel in BG may select a *subset* of the total representations available to PFC. Recurrent connectivity within cortex itself may then afford a complementary selection role, disambiguating the individual representations forming the selected subset. While this scheme necessarily results in a partial loss of information in BG, the architecture of the loop as a whole is such that representations in cortex – and subsequently the richer information therein – can be preserved.

We further propose a partially distributed or ‘feature-based’ coding scheme may be utilised in PFC for well learned information. This affords a more predictable system of overlap between representations, whereby those representations sharing a particular feature (e.g., a common goal) will share the same neural substrate to represent that feature. This structure may in fact be *more* plausible than a fully distributed scheme (84–86), and necessarily results in a systematic pattern of ‘overlap’ between similar representations, with clear conceptual benefits regarding the similarity of representations which encode similar contexts, such as generalisation (87) and graceful degradation (88). Indeed, partially distributed codes have been discovered in human temporal cortex (89) and previous computational work endorses ‘compositional’ schema representations (72).

We develop a model of the subset-selection hypothesis in several stages. First, construction of the associative loop with PFC (which is, in turn, broken into two steps for ease of exposition). Second, the relevant sensory and contextual (cognitive) inputs to the motor loop are described. Third, we give the complete architecture with both loops and report simulation results for normal behaviour. Finally, we examine two contrasting theories of the nature of the breakdown of contextual information underlying action slips and action disorganisation, and demonstrate the superior ability of one approach to account for observational and experimental data. In doing so, we reconcile the differences between the previous competing IAN and SRN models.

## 2. Modelling I: Associative loop

### 2.1. Basic architecture

We based our model on the Humphries & Gurney (41) model of the motor BGTC loop (hereafter known as the TC-GPR). The basic architecture of this model, incorporating the major pathways through the BG (34), is illustrated in figure 1a. Briefly, this model incorporates D1 and D2 expressing regions of striatum (SD1, SD2), subthalamic nucleus (STN), the external globus pallidus (GPe) and a region encapsulating substantia nigra pars retiulata and the internal globus pallidus (GPi/SNr). SD1 and SD2 function as the major input regions of the BG, receiving afferents from motor cortex. These give rise to selection and control pathways through BG, the activity of which are modulated by STN. These pathways coverge at the GPi/SNr, which functions as the major output nucleus of the BG. Projections from GPi/SNr modulate activity in thalamus, which is arranged in a recurrent loop with cortex. For more detail, see (39–41).

**Figure 1.**
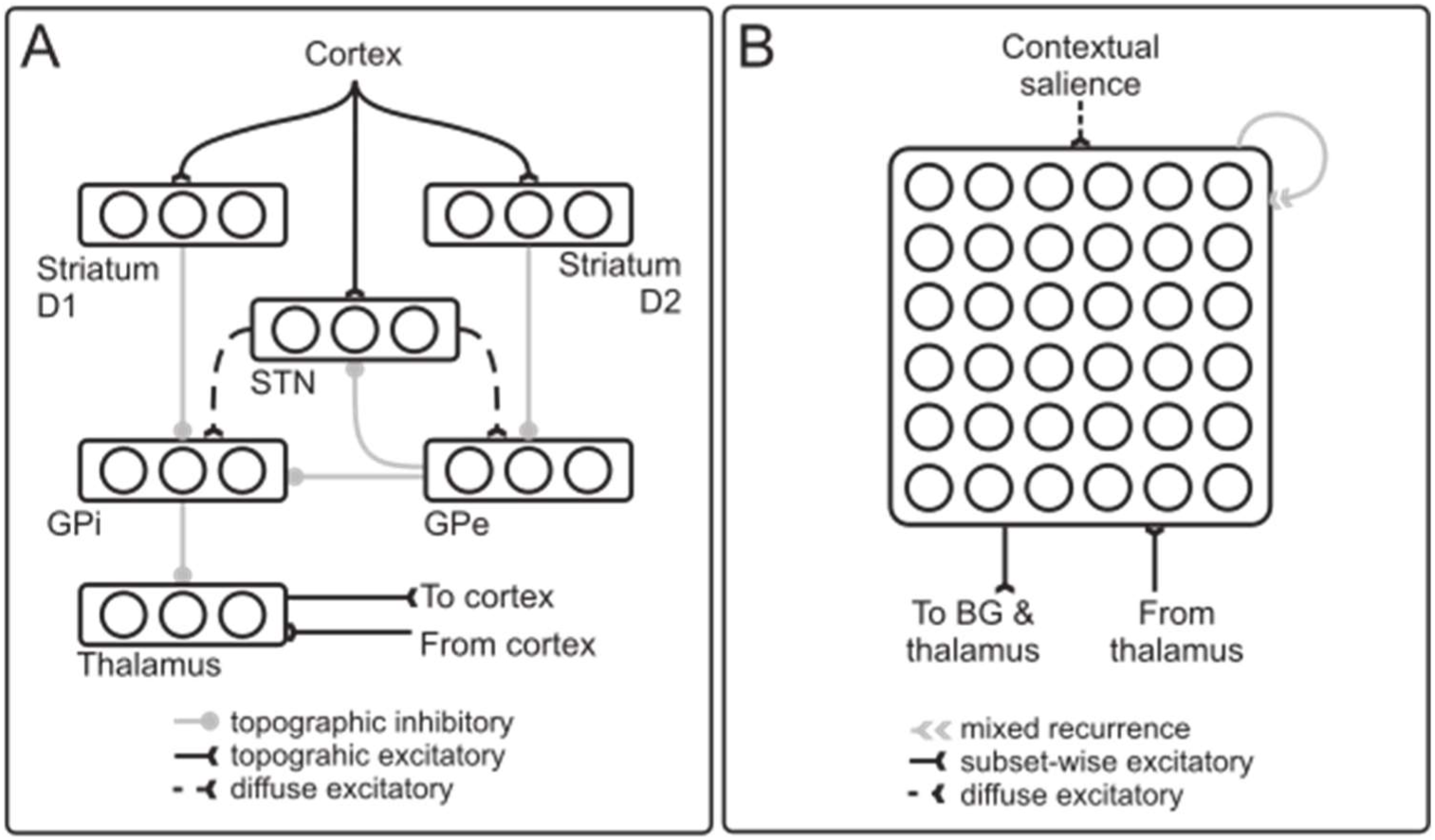
A) Schematic diagram of the TC-GPR architecture. In this paper we propose an expansion of cortical dimensions to allow distributed representations. B) New organisation of the present model. Multiple, high-dimensional representations across a 6×6 node cortex project to and receive afferents from each channel in basal ganglia.

The original model was arranged in ‘channels’, each of which represented a motor action. Externally supplied ‘saliences’ to cortex and striatum drove activation of the channels; the relative strengths of the saliences to each channel represented the relative urgency with which each action was being activated. ‘Selection’ was defined in the model by activity in a cortical channel reaching a pre-defined selection threshold. This model was shown to display several desirable properties for a selection mechanism (39–41).

To adapt this motor loop model to deal with more complex associative information, we replaced the cortical region of the TC-GPR with a 6×6-node recurrent network (figure 1b). All other modelled nuclei remained channel-wise, consistent with the original model. In contrast to previous GPR models we are concerned with the selection of *a pattern* of activity across several nodes in PFC, rather than the selection of a single element or channel. We implement representations of task context as conjunctions of ‘features’, where individual features are represented by particular nodes within PFC. Critically, each feature may be a component of multiple representations, resulting in overlap between representations sharing features.

We developed a set of six simple feature-based representations with which to analyse the functional capabilities of the proposed architecture. These are illustrated in (figure 2). Each consists of a conjunction of two features. Each feature is signalled by the activation of two adjacent nodes, and each representation shares one feature with two other representations. Each BG and thalamic channel is reciprocally connected with two of these representations. Thus, if a given BG channel is selected, *two* PFC representations are subsequently disinhibited. We refer to the set of all the representations disinhibited by a single channel as a *‘subset’*. Local recurrent connections within PFC itself then perform a supplementary selection role, and further select between the two disinhibited representations, resolving any remaining ambiguity and suppressing activation of the ‘losing’ representation. Connectivity for this subset-based selection scheme in the model is illustrated in figure 3. Apart from connections with and within PFC, all connectivity remains faithful to the TC-GPR model.

**Figure 2.**
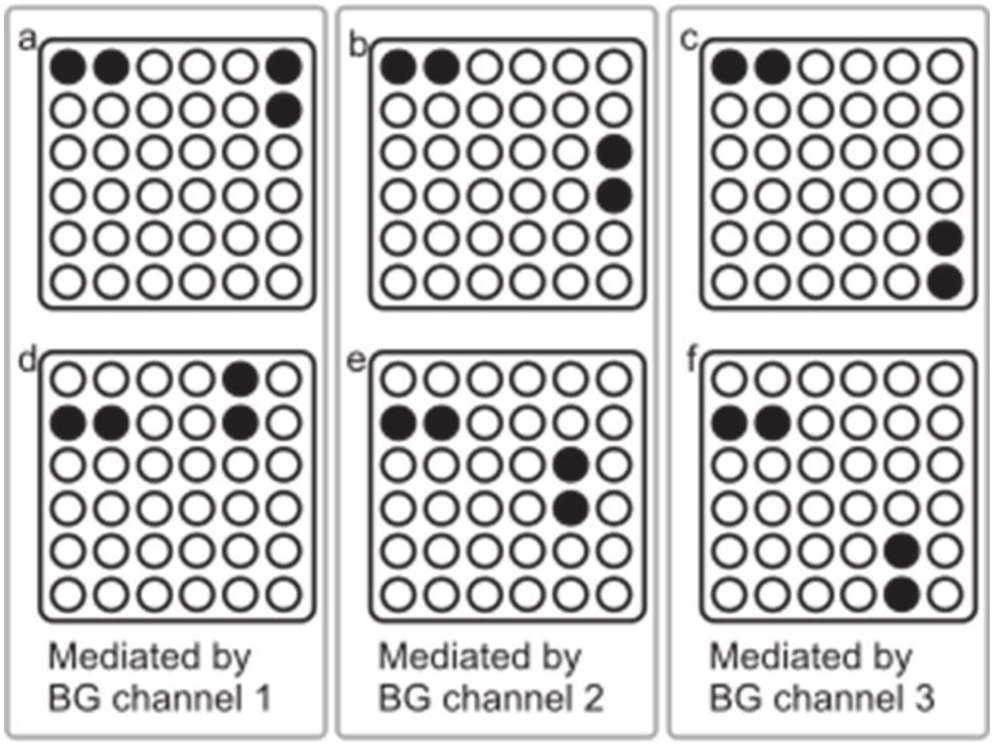
Feature-based representations encoded by the recurrent PFC in the present model. Each PFC representation comprises two features; each feature is represented by two adjacent nodes. Each BG channel supports multiple representations (see text for details). Filled nodes = active, and empty = inactive for that representation.

**Figure 3.**
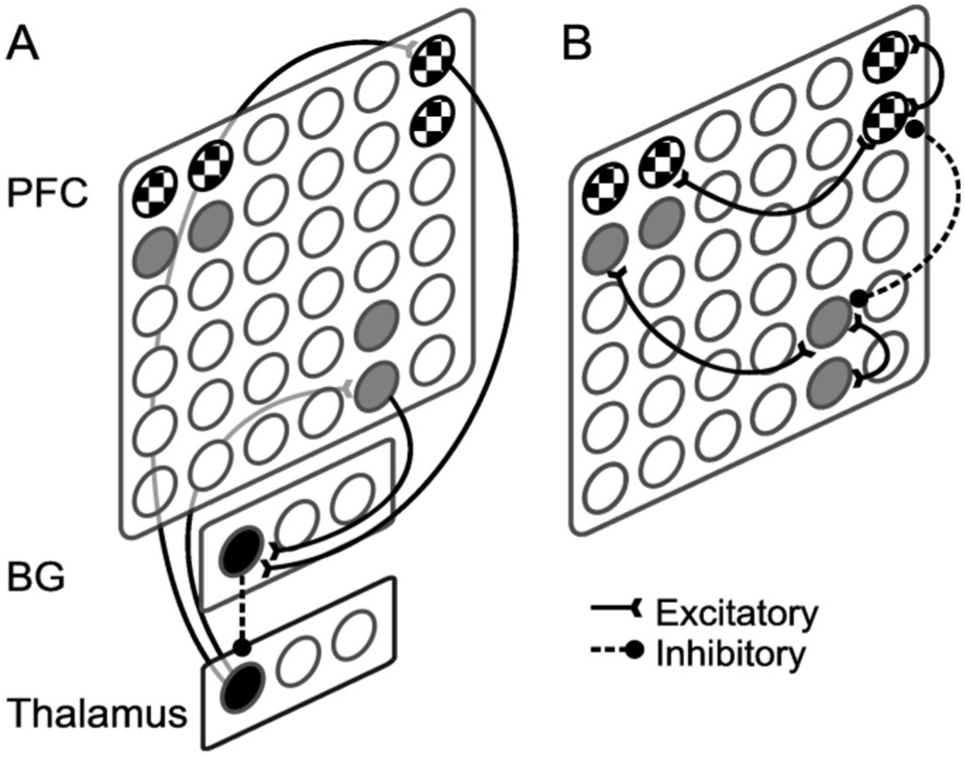
Schematic illustration of the connectivity underlying the subset-based selection scheme in the current architecture. A) Two distinct PFC representations (grey and check nodes, respectively) excite a common channel in BG (black node). The same PFC representations receive excitation from a corresponding common channel in thalamus. B) Within PFC, excitatory and inhibitory recurrent connections support the selection of a single representation from among the subset selected by the common BG. Illustrated connectivity is representative rather than exhaustive.

#### Formal description

Each node in the model represents a population of neurons and is modelled as a leaky integrator unit as described by,

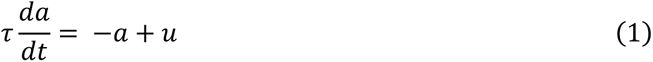

(note that we will loosely refer to such objects as ‘neurons’ even they stand for neural populations). Here, *a* is the neuron’s activation value, *u* is its input and *τ* is the representative membrane time constant. The input was calculated as weighted sums of inputs from a neuron’s afferent connections. The output *y* of each neuron is calculated by the following piecewise linear function of its activation:

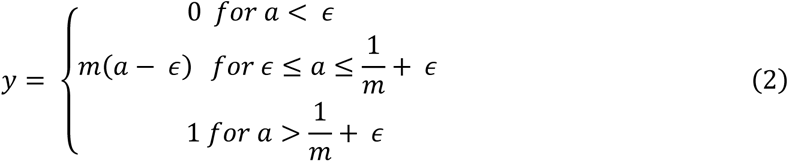

Where *ɛ* is the output threshold, and *m* the gradient of the output function.

Weights between intrinsic BG nuclei and thalamus were consistent with those in (41). Weights within PFC were individually hand tuned to support the selection of the representations illustrated in figure 2. The model’s input was characterised by saliences as in the original model; these were scalar additions to the activations of nodes comprising a given representation. ‘Selection’ was achieved when all nodes of a given representation had outputs *y* ≥ 0.9. All weights are given in Supplementary Material A.

### 2.2. Tests of selection

We tested the model under a series of different inputs, to test its ability to perform different types of selection and related functions:

#### i. Selection – different BG channels

Two PFC representations disinhibited by *different* BG channels were supplied with non-zero salience while others had a salience of zero. This mainly tested the selection ability of BG.

#### ii. Selection – same BG channel

Two PFC representations disinhibited by *the same* BG channels were supplied with salience. This predominantly tested the supplementary selection ability of PFC.

#### iii. Maintenance and switching

One PFC representation was supplied with salience until it was selected. External salience was removed to test the ability of the model to maintain selection (in ‘working memory’). Finally, salience to a competing representation was introduced. This tested the model’s ability to switch to a new representation in response to changing salience.

#### iv. Switching from a supported PFC representation

This was the same as the previous simulation, except that the salience to the first PFC representation was not removed. This tested the model’s ability to switch in the presence of competing input for an already-selected representation.

## Results

The model showed successful selection and switching behaviour in all simulations. Outputs from PFC and GPi/SNr are shown in figure 4. Test i results are displayed in column A. The top graph shows the strength of input saliences to representations 1 (dashed) and 2 (solid). The two components of the PFC representations show initial activity in both representations. With the aid of selective mechanisms in BG (bottom panel shows output nucleus activation), the competition between representations 1 and 2 in PFC is resolved; activity in representation 1 is enhanced via positive feedback in the thalamocortical loop, and activation in representation 2 is suppressed due to increased inhibition from BG output nuclei. Outputs for test ii-iv are shown in panels B-D and show appropriate selection and deselection of winning and competing representations according to the input saliences illustrated in the top row.

**Figure 4:**
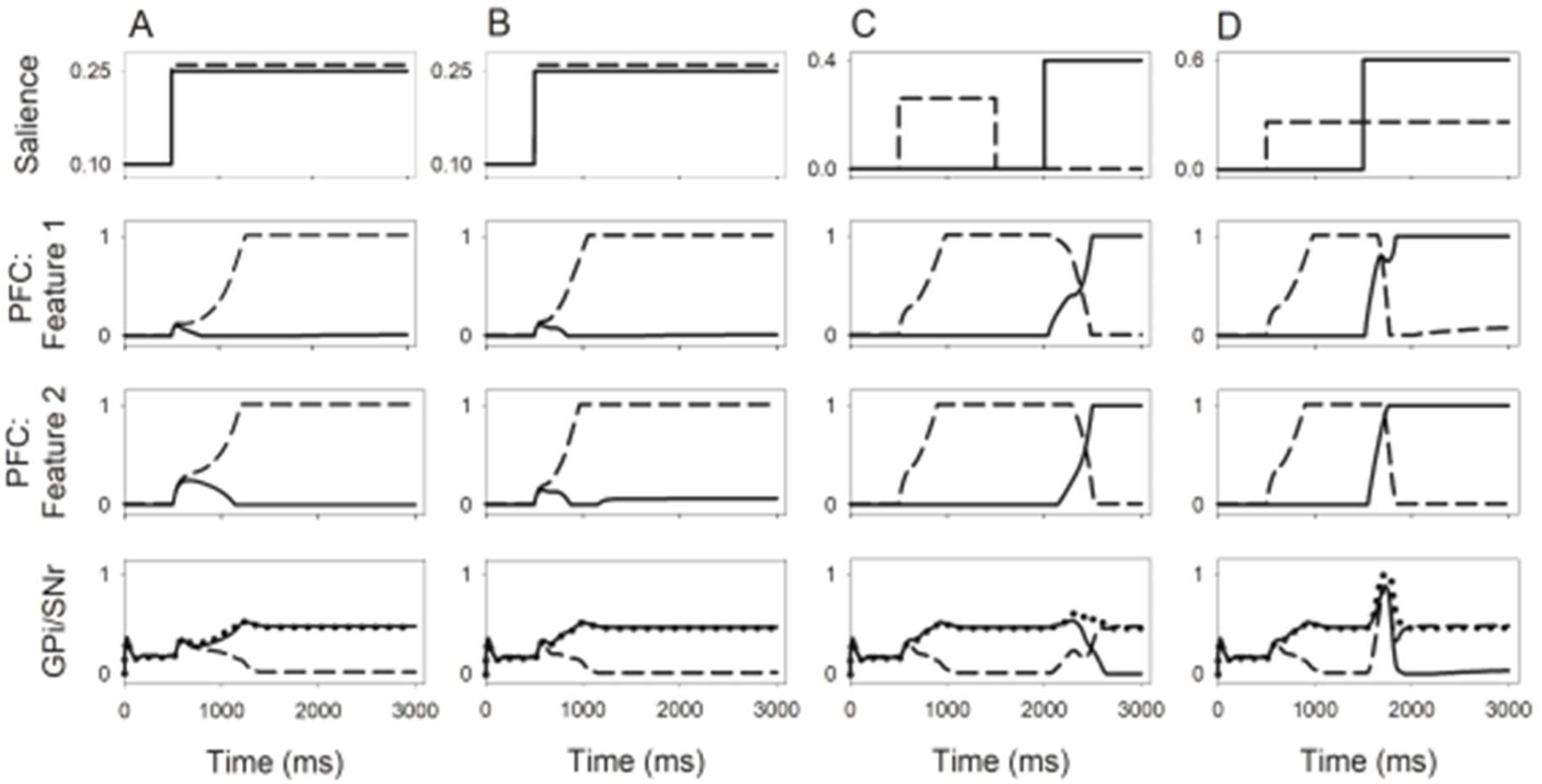
Output of key nuclei for simulations i-iv. A) Salience (top) was supplied to representations 1 (dashed) and 2 (solid) at t=500ms (see fig 2). Middle graphs show the average output for features of these PFC representations. Selection processes in PFC and BG (bottom) drive successful selection of the most strongly supported representation (dashed). Activation of the competing representation 2 (solid) remains below the selection threshold of 0.9. B) Salience was supplied to representations 1 (solid) and 4 (dashed). Here, both representations are mediated by a common BG channel. Selection of this channel is observed in BG output nuclei (bottom); PFC recurrence drives selection of the more strongly supported representation 1. Selection happens more quickly here than in simulation 1, because there is no competition to resolve within BG itself. C) Selection of representation 1 (dashed) is maintained after external salience driving selection is removed at t=1500ms. Salience is applied to representation 2 (solid) at t=2000ms. These representations are supported by distinct channels in BG. Timely deselection of representation 1 and selection of representation 2 are seen. D) Selection of representation 1 (dashed) is followed by the application of additional salience to representation 2 (solid) at t=1500ms without corresponding removal of salience to representation 1. Switching to representation 2 is observed, demonstrating the model’s ability to appropriately deselect a supported representation.

## Interpretation

These results demonstrate the ability of the proposed architectural scheme to mediate appropriate selection, maintenance and deselection of multiple cortical representations, where those representations outnumber the total channels in BG. The model is able to resolve competition between multiple representations, both within and between subsets, and is able to switch between representations upon the introduction of a sufficiently strong competing input salience.

The novel subset-selection hypothesis was proposed in response to a structural incompatibility between the organisation of information in PFC and BG. Our model resolves this conflict and allows the traditionally accepted topographical or ‘localist’ organisation BG to communicate in a clearly defined way with the multidimensional representational scheme in PFC. In doing so, the model is consistent with neurophysiological evidence and computational arguments that PFC is likely to utilise distributed representations, as well as overwhelming evidence that BG is arranged in distinct channels. Furthermore, it begins to address evidence of heavily converging projections from cortex to BG, where it suggests that wide regions of cortex are mediated by smaller areas in BG.

## Comparison with previous models

Though previous models of associative BGTC circuits have successfully showed selection and/or gating of cognitive information to working memory, the present model is only one to perform competitive selection, maintenance and *de*selection of distributed representations by BG. For instance, while most models of BG at associative levels of the BGTC hierarchy address the mediation of working memory by BG in some form, the precise role of BG in working memory is debated (90, 91), with several accounts emphasising the gating of information into working memory (74, 92–95), versus maintenance itself (96–100). Notably, models of working memory including distributed PFC representations tend not to rely on BG involvement for maintenance, instead focusing on cortico-centric mechanisms such as bi-stability in cortical cells (74, 101) and local recurrence (94, 102, 103). Conversely, models which utilise the thalamocortical loop to maintain information, as here, tend to rely on a fully localist architecture (96–99). To the best of our knowledge, our model is unique in proposing an architecture which utilises thalamocortical feedback for maintenance in concert with distributed representations.

Dominey and colleagues (104, 105) utilise distributed PFC representations of context for a saccade task requiring active maintenance of sequential information. Superficially, these models appear similar in structure to the present model. However, a minimal representation of BG and the presence of selection processes in superior colliculus suggests that BG is predominantly a locus for associative learning in these models, rather than performing crucial roles in selection or maintenance. Whether true self-maintenance is achieved is unclear, since several of their simulations involve tonic visual input, which may provide critical ongoing support to maintain the current state. Furthermore, implausibly long time constants were used in PFC, and it is difficult to determine whether the model would be capable of the same maintenance with more plausible key parameters.

Frank, O’Reilly and colleagues (74, 95, 101, 106, 107) have produced what is arguably the most substantial modelling contribution to understanding working memory functions of BG to date. Again, these appear similar to the present model. They, however, emphasise gating functions of BG, placing maintenance solely in cortex. Indeed, they actively discount recurrent thalamocortical loops for maintenance, suggesting that it is not possible to maintain multiple representations in cortex using this mechanism whilst adhering to the principle of corticothalamic convergence (101). Here we have demonstrated this is not the case. They also claim that thalamocortical interaction for maintenance is impractical because it requires the continued disinhibition of thalamus for the duration of maintenance. We argue that this continued disinhibition is necessary, due to the requirement of BG-based inhibition for *de*selection. This is a feature which few models of cortical-BG interactions explicitly address, despite neurophysiological evidence existing to suggest the ongoing involvement of BG during maintenance (90). Without inhibition-driven deselection, the mechanism for inactivating a selected representation remains an open question (92, 101), as does the general function of tonic inhibition from BG.

The gating role postulated for BG in these models may be further limited, where their framework implies no direct competition between channels in BG. In these models, PFC consists of a series of ‘stripes’, each of which represents a particular category of information. Stripes consist of a series of nodes, each node representing a particular instance of that category. Each node in BG ‘gates’ information into a stripe. Theoretically, all PFC stripes may be updated simultaneously, providing no role for the well-evidenced competitive selection function intrinsic to BG. Their architecture may struggle with simultaneous competing inputs to the same stripe: either BG updates the stripe, in which case both stimuli would be stored in working memory, or it does not, in which case neither would.

In contrast, the present study provides mechanisms for true competition-based selection in BG, and true maintenance of selected representations via BG. Thus, there is no potential for ambiguous updating, nor the need for constant external input. This unique combination of functionalities in a model of the BGTC loop architecture stems directly from our novel organisation.

## 3. Modelling II: Dual loop architecture

### 3.1. Defining sequential tasks

Having established the functionality of the associative PFC loop model in principle, we now examine its ability to mediate more complex representations for a particular set of tasks and embed it in a dual-loop BGTC architecture to integrate cognitive and motor selection functionality. For consistency with previous models and to explicate the task requirements that constrain modelling decisions, (16, 25, 32, 68) we defined simple tea– and coffee-making tasks. These are hierarchically organised into goals, subtasks and component actions. This is shown in figure 5. The tea task consisted of seven actions across three subtasks; these were fixed such that they had to be performed in a particular order. The coffee task consisted of ten actions in four subtasks; these were mostly fixed, although the order of two subtasks in the coffee task could be switched. This resulted in three valid sequences comprising 22 action representations. The tasks (and subtasks) shared several actions, but actions were represented distinctly for each context in which they appeared. For instance, each step in the ‘add sugar’ subtask was represented three times: once for the tea task, and twice for the coffee task, where sugar is added before or after milk. These unique context specific actions are numbered in Table 1.

**Figure 5:**
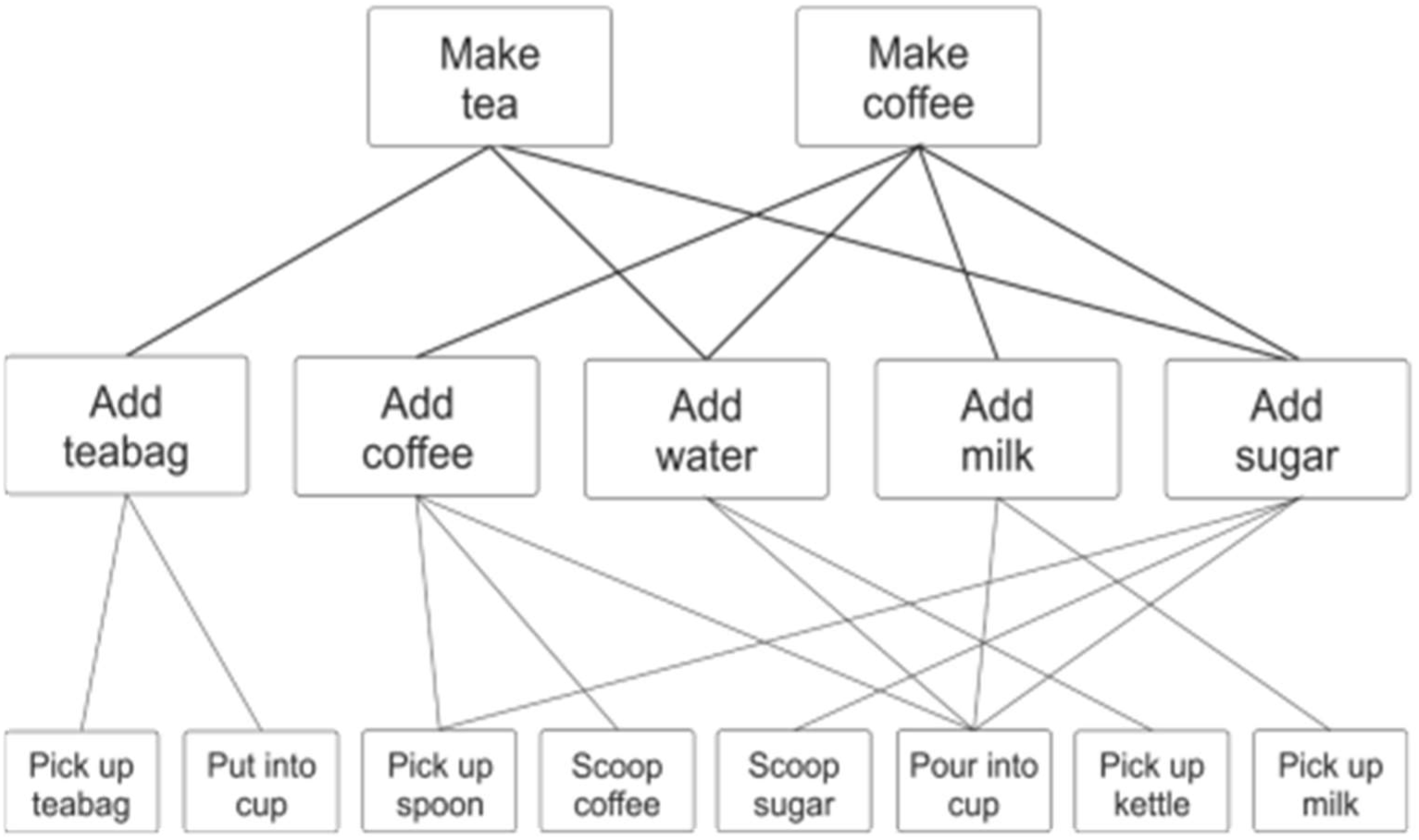
Hierarchical organisation of model tasks, illustrating the ‘re-use’ of actions and subtasks

**Table 1:** Tasks and their numbered actions in order.

| Make tea | Make coffee 1 | Make coffee 2 |
| --- | --- | --- |
| T1: Pick up teabag | C1: Pick up spoon | C1: Pick up spoon |
| T2: Put into cup | C2: Scoop coffee | C2: Scoop coffee |
| T3: Pick up kettle | C3: Pour into cup | C3: Pour into cup |
| T4: Pour into cup | C4: Pick up kettle | C4: Pick up kettle |
| T5: Pick up spoon | C5: Pour into cup | C5: Pour into cup |
| T6: Scoop sugar | C6a: Pick up spoon | C6b: Pick up milk |
| T7: Pour into cup | C7a: Scoop sugar | C7b: Pour into cup |
|  | C8a: Pour into cup | C8b: Pick up spoon |
|  | C9a: Pick up milk | C9b: Scoop sugar |
|  | C10a: Pour into cup | C10b: Pour into cup |

In addition to shared subtasks and actions, this task set includes actions that are repeated within a single task. Furthermore, we added the constraint that one subtask (‘add sugar’) does not result in a ‘visible’ environmental change. These features tax the model by requiring it to maintain an internal record of the goal it is working towards, and the actions it has performed to successfully complete a sequence. Reusing a single action within the same sequence, and utilising the same actions in different tasks have caused difficulties in previous modelling work (e.g., (108); see also (25, 105) for discussion).

We omitted orienting (‘look at’) and release (‘put down’) actions from the task set. This improves manageability of a large model and reduces complexity, allowing us to focus on the ‘crux’ and facilitatory actions within each subtask. Here, ‘crux’, is used in the sense of (28) to refer to the individual action which achieves the goal of the subtask. Here, there is no qualitative difference between orienting or release actions and the manipulative actions we include, thus we do not sacrifice theoretical value by excluding these actions. To account for release actions, it is assumed that whenever a ‘pick up’ action is selected, it is preceded by a ‘put down’ of any currently held object. Orienting actions are accounted for by use of a ‘deictic’ scheme, detailed in **Error! Reference source not found.** below.

### 3.2. Expanding the associative loop

Each of the 22 context specific actions detailed in Table 1 required unique PFC representation for successful performance of all task variants. The PFC-loop model was augmented to deal with the greater number of representations required and the higher dimensionality of each representation. In the new instantiation, PFC comprised a 7×7 network. Weights therein were hand-tuned to support the set of 22 representations. In our feature-based representational scheme, each feature was encoded consistently across representations by two nodes. Features active for a given stage of the task included the relevant goal, subgoal, action and object(s) required to complete that stage. Thus, each representation comprised the set of relevant features for that stage. Example representations are illustrated in figure 6. Where subtasks could be performed in a different order, nodes tracking the order of performance were included. These were termed ‘rank order’ nodes, and neurons with this apparent functionality have been documented in neurophysiological studies (57, 109, 110).

**Figure 6:**
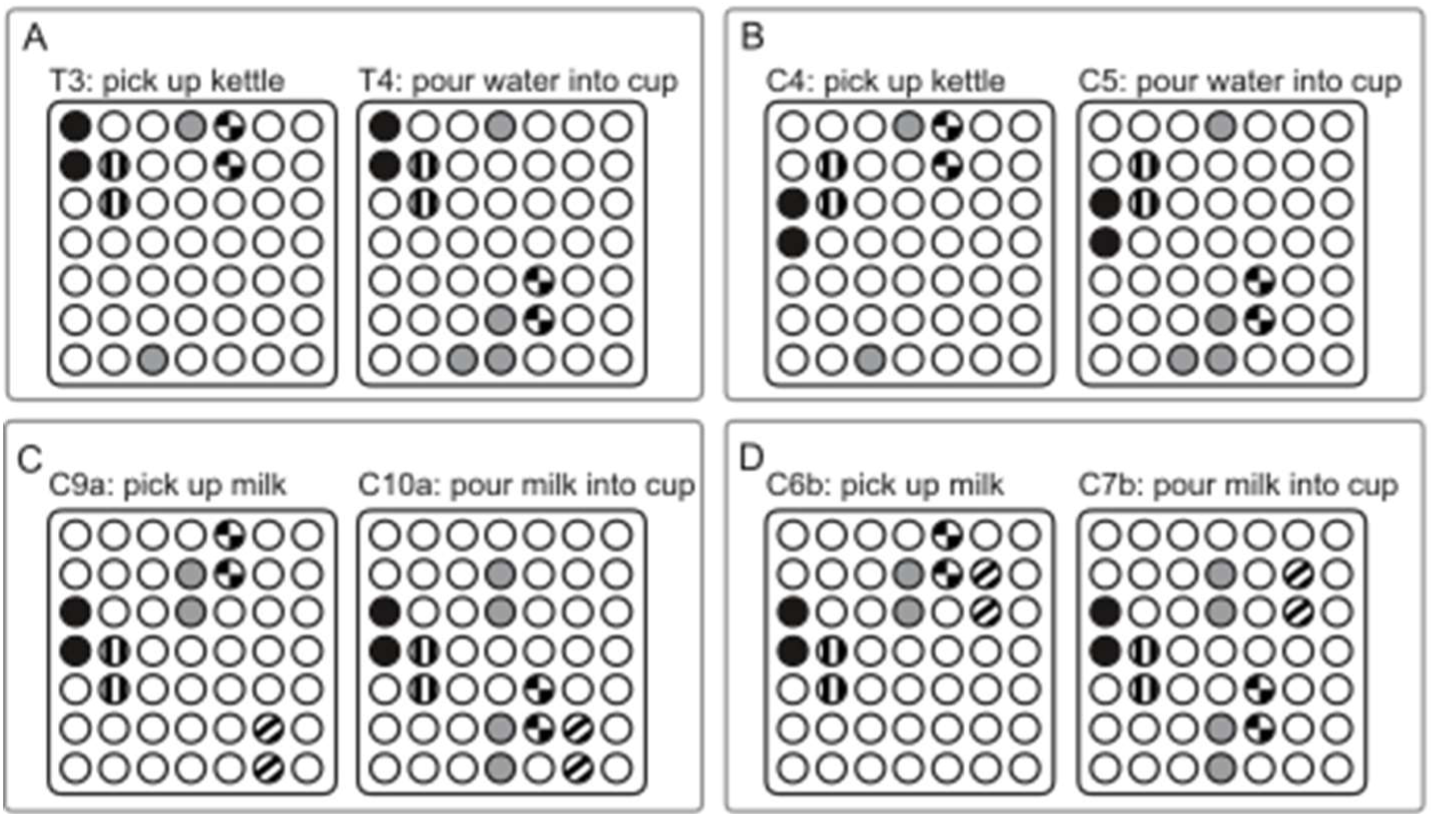
Example representations used by the model to represent task stages. Colours show the different level of information represented by pairs of nodes. Black = goal/task; vertical stripe = subtask; grey = object(s); check = action; diagonal stripe = rank order. Each representation contains sufficient information in the activated ‘features’ to distinctly represent that stage of the task. Example: panel A shows the representations for the two stages of the ‘pour water’ subtask for the tea task. Common black and vertical striped nodes representing the ‘make tea’ goal and ‘pour water’ subtask are active for both stages. Grey object nodes representing ‘kettle’ are active for both stages. Grey object nodes representing ‘cup’ are active for the second stage. Different checked action nodes are active for the two stages, ‘pick up’ in the first stage, and ‘pour’ for the second. Panel B shows the same subtask as panel A but for the coffee task. Note all the same nodes are active except for the black goal nodes. Panels C and D show representations for the milk subtask in the coffee task, performed before and after the sugar subtask, respectively. Diagonal striped rank order nodes distinguish these versions of the subtask, allowing the model to track internally whether the milk or sugar subtask was performed first, thus facilitating an internal ‘memory’ of the task history.

Like the earlier instantiation, this PFC-loop is organised such that each BG channel disinhibits a subset of the total representations. Here, each channel disinhibits representations which share a common subtask. For example, four representations correspond to the two actions comprising the add water subtask: those in the coffee task context, and those in the tea task context (actions T3-4 and C3-4 in Table 1). These are heavily overlapping (see figure 6). Each of these four representations is disinhibited by a single BG channel, and recurrent inhibition within PFC resolves remaining ambiguity. Likewise for the remaining four subtasks, resulting in a 5-channel BG in the updated PFC-loop model; one channel for each of the 5 subtasks (figure 5).

### 3.3. Incorporating the motor loop

#### Modelling sensorimotor information

The motor centres of the brain must continuously integrate dynamic sensory (‘bottom up’) and cognitive (‘top-down’) information to perform flexible, goal-oriented action selection. Existing single loop GPR models of BG have incorporated an abstract global salience signal reflecting the overall ‘urgency’ for an action. This signal encompassed many different types of functional information. Here, we begin to deconstruct this salience signal by examining some of its possible associative and sensory components.

Since the motor loop is concerned with physical actions, an important component of sensory information to this loop is necessarily action-specific, regarding suitable actions given the immediate environment; we refer to these as ‘candidate actions’. Candidate actions are probably largely object-oriented. This relates closely to the notion of ‘action affordances’ (111), or features of the immediate environment relating to physical attributes of perceived objects and the actions they promote. Affordances are likely derived from information from visual perception and object recognition (112, 113). Several parietal regions appear to be loci of action-related information derived from sensory sources (114). In particular, the anterior intraparietal area (AIP) is implicated in the specification of object-related grasping movements (115), and is likely a critical locus of visuomotor transformations for object manipulations (116). Projections from AIP impinge on motor territories of the BGTC hierarchy, such as premotor cortex (117) and putamen (118). AIP receives afferents from inferotemporal cortex (IT) (117), which is widely accepted as the primary locus of object recognition and the terminus of the ventral, or ‘what’, visual pathway. It is also the target of afferents from the dorsal visual stream (117), believed to provide information for movement specification based on visual, rather than semantic, properties of objects. These pathways suggest two ‘routes to action’ (112, 119), in which AIP is involved in the integration of semantic and visual information for the specification of grasp. It is likely, then, that an important component of the salience signal for a given action in BG is derived from this AIP, as has been implemented in existing models (114, 120).

#### Cognitive influences on motor selection

Cognitive information selected by prefrontal-BGTC loops must be communicated to motor regions to successfully modulate action selection. Under normal circumstances, sensory influences likely pose a more direct constraint on action selection than contextual information. This reflects the notion that sensory information ultimately defines which actions are appropriate or possible in the current environment (72). Contextual influences conversely support and modulate action selection by *biasing* the selection of a single action from a set of candidate actions (121). Such a modulatory effect may be implemented in the motor BGTC loop via a direct influence on striatum: since the influence of the BG on action selection is not directly excitatory but disinhibitory, additional excitation to motor striatum will not activate new action representations but will modulate the activation of actions which already have some degree of cortical excitation.

While BGTC loops have historically been regarded as heavily segregated (34, 37, 77), this organisation is now known to be more nuanced, with a range of evidence existing for a significant degree of interaction between loops. Loops are connected at the cortico-cortical (122–124), thalamocortical (125, 126), and striatonigral and striatopallidal levels (127, 128). Most notably, evidence for overlapping corticostriatal projections in primates has been found for multiple motor and premotor territories (129, 130), and also from apparently distinct territories including associative and premotor (131), and limbic and associative (132). A similar pattern of corticostriatal convergence has also been evidenced in humans (133). This corticostriatal overlap implies a direct influence from cognitive cortices on the pre-motor BGTC loop, and is particularly relevant for understanding the cognitive biasing of motor action.

### 3.4. Integrating the loops – the RoSe-BaL model

The evidence described above suggests that the ‘salience’ signal of previous motor-GPR models includes bottom-up affordance information and top-down information based on task context. Affordance specification is also subject to top-down influences in the form of goal-directed visual search for task-relevant objects (134). Bringing this together with the associative loop model including PFC, and the original GPR loop with motor BG gives the dual-loop architecture for the complete RoSe-BaL model (figure 7). This functional architecture includes two pathways between loops based on the pathways described above to translate cognitive information into motor action.

**Figure 7:**
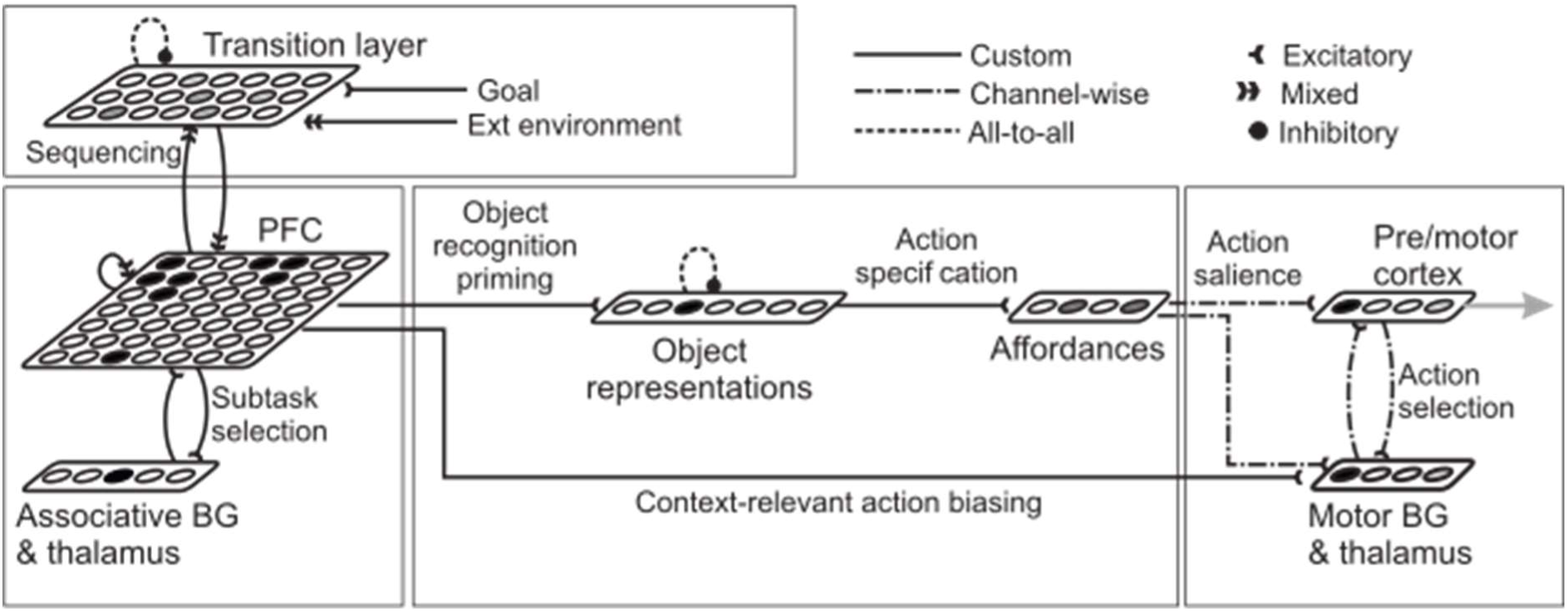
Full RoSe-BaL model architecture. Model consists of four functional regions: action selection region in the motor BG loop (right), schema selection in the associative loop (bottom left), task sequencing in the transition layer (top left) and the translation of cognitive information to action-related parameters (centre). For ease of display, a simple row of nodes subsumes the BG architecture in both associative and motor loops.

One of these pathways exploits cognitive information to ‘prime’ recognition and attention to task relevant objects (‘object representations’ in figure 7). The activation of a node in this region indicates fixation of an object represented by that node. This in turn activates affordances for actions on those objects. Affordances specify candidate actions for selection and provide salience for those actions to pre/motor cortex and motor striatum. This pathway is based on that from IT through AIP to motor BGTC loops and mimics a projection from PFC to IT embodying a top-down influence on object recognition. Though highly simplified, this pathway is supported by neuroanatomical evidence (135), and is used here as a substitute for a targeted visual search process, driving fixation of the target object. This pathway also accounts for the effects of ‘orienting’ via a deictic scheme (136). Thus, any action selected by the motor loop is deemed to be directed towards the fixated object. Similar schemes have been implemented in previous models of routine action (16, 25).

The second of these pathways originates in action-related context nodes in PFC, and projects to the corresponding action in motor striatum. This provides a goal-directed biasing influence on action selection. Its influence in striatum, rather than cortex, means it can only help to disinhibit an action representation already supported by sensory influences; this prevents object-directed actions being triggered by context alone.

#### Sequencing

Stability and sequencing are both required in the PFC loop, whereby a sequence of PFC representations must be evoked to drive serial action, but each one must be individually stable for the time taken for downstream action to be selected. This trade-off is rarely encountered in connectionist models, since most use either fixed point attractors to study retrieval of single memories in isolation, or dynamic attractors to study sequential processing. However, this trade-off has been addressed in the computational literature, and where stability of representations is required, an external mechanism to drive sequencing is shown to be necessary (137). As such, we implemented a sequencing mechanism external to the main body of the BGTC loop architecture, in order to provide the necessary external bias to drive dynamics between individually stable PFC representations and evoke sequencing. In this solution (‘transition layer’ in figure 7), each PFC representation has a corresponding transition node which, when active, provides excitation to all PFC nodes comprising that representation, thus functioning as the primary source of salience to the PFC loop. Afferents to the transition layer from PFC convey information regarding the current task context. This influence from PFC ‘primes’ – or activates to a sub-threshold level – all transition nodes which correspond to potential subsequent PFC representations. Several transition nodes may receive this subthreshold activation at a given time, particularly where multiple actions may feasibly follow the current one. When an action is selected by the motor loop, a corresponding change in a simplified model world occurs. This world is represented with a series of binary nodes indicating the task-relevant status of objects (whether an object was ‘on-table’, ‘in-hand’, etc). Alterations in the world are propagated to the transition layer as a phasic burst of excitation, again to those nodes which correspond to compatible subsequent actions.

This combination of influences from PFC and the model world preferentially excites a single transition node, which then provides salience to the subsequent PFC representation which is most compatible with both internal and external influences. If this salience is of sufficient strength and duration, a new representation will be selected and any previously selected representation ‘deselected’ through inhibitory and competitive selection mechanisms, thus driving sequential action. The combination of internal and external influences allows the transition layer to integrate information regarding the environment and information about the task history implicit within PFC. This avoids problems associated with ‘chaining’ (13).

#### Formal description

Like the previous model, all nuclei were modelled with a series of leaky integrator neurons according to equations 1–2. Weights associated with projections are detailed in Supplementary Material B.

### 3.5. Full task simulations

#### i. Basic sequencing

We tested the model’s ability to perform tea and coffee tasks. Input to the model consisted of activation of transition node ‘T1’ or ‘C1’, which corresponded to the first stage of the tea or coffee task, respectively. We ran 200 trials each for tea and coffee tasks. Simulations were run for 30s using a timestep *dt*=1ms, with a time constant τ=40ms for all nuclei.

The RoSe-BaL model could consistently perform all three sequences without error, and milk-first and sugar-first versions of the coffee task were spontaneously performed with roughly equal frequency. This demonstrates the ability of the architecture we have proposed to integrate multiple sources of information to mediate goal-directed action sequences. It is notable that component actions may be recruited in a flexible order, independently of the action just performed, independently of the goal, and multiple times in one sequence. The model is also able to maintain a record of the performance of previous actions which do not result in a visible change to the environment (e.g., adding sugar).

##### Analysis

Outputs from the object representations region and pre/motor cortex for a coffee trial are illustrated in figure 8. Sequential selection is seen as the consecutive traversal of the selection threshold by the outputs of each action channel in pre/motor cortex. Action selection is driven in part by activation of object representations, via actions specification along the PFC-objects-affordances pathway. This causative influence is reflected in an offset in the dynamics of the two regions, where activation of object representations precedes that of action representations in cortex.

**Figure 8:**
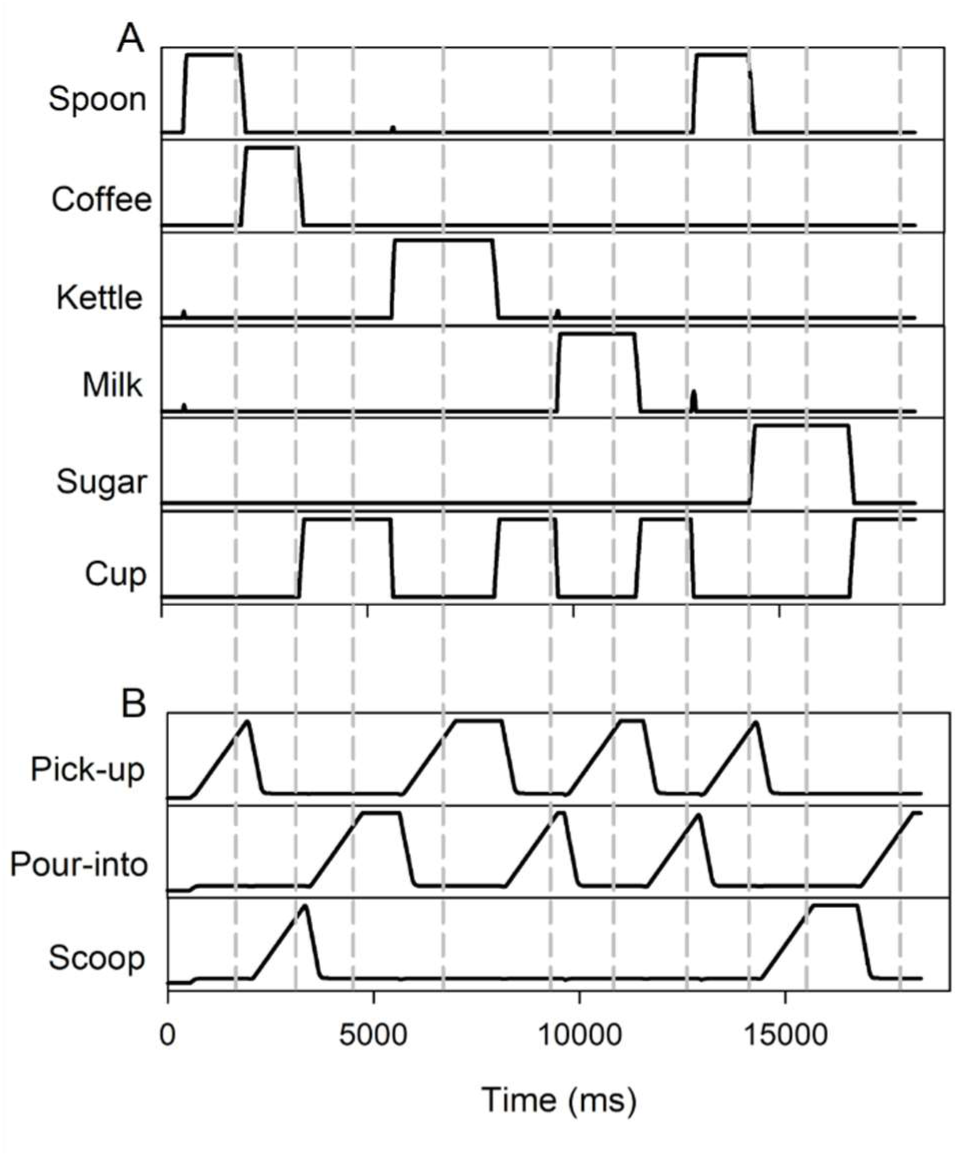
Time course of activation of key nuclei for a typical coffee-making trial. A) Output of object nodes in the object representations region. B) Output of action channels in pre/motor cortex. Action selection times are indicated by the grey dashed lines. Activation of object nodes indicates ‘fixation’ of that object. Activation of action channels indicates selection of that action; fixation of objects occurs prior to action selection. The sequential activation of the targeted actions is visible: ‘pick-up-spoon’ (t∼1800ms), ‘scoop-coffee’ (t∼3000ms), ‘pour-into cup’ (t∼4500ms), etc.

The time course of activation of PFC nodes encoding the goal, subtasks and actions for the coffee task is illustrated in figure 9a. This drives action selection in the motor loop, via the PFC-objects-affordances pathway and the PFC-BG pathway. Selection and deselection of feature nodes in PFC is caused by transition node activation triggering a switch in the selected PFC representation. The time course of transition node activation is shown in figure 9b; comparison with 9a demonstrates maintenance of a selected representation in the absence of transition node activity.

**Figure 9:**
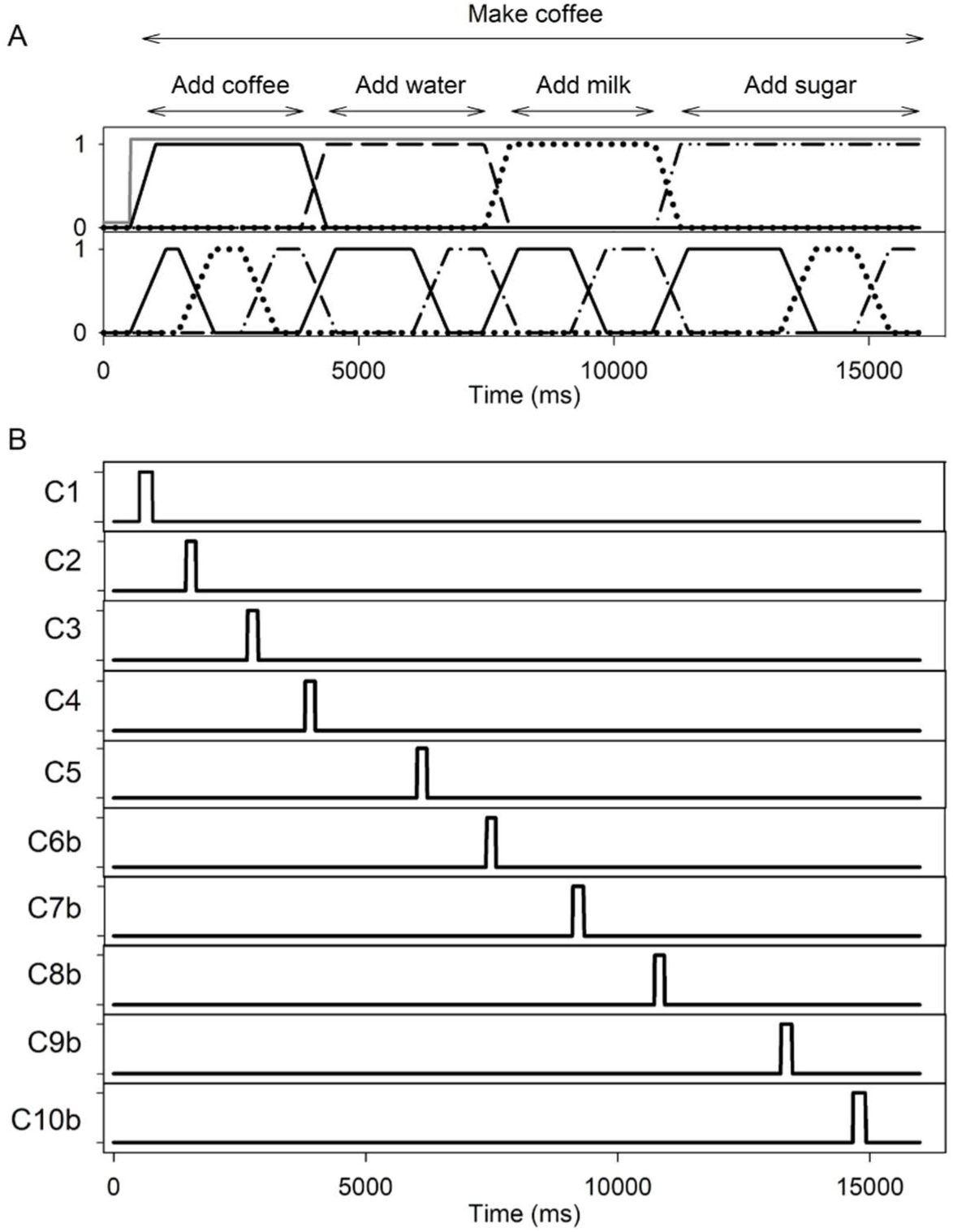
A) Time course of activity of PFC nodes encoding goal (top, grey), subtasks (top, black), and actions (bottom). For visibility, moving averages are shown for subtasks and actions using window sizes of 500 and 700ms, respectively (in reality, outputs were close to binary in nature). B) Time course of transition node output, for transition nodes C1-C10b, which provided excitation to the corresponding PFC representations C1-C10b. Note transience of transition node output, and maintenance of PFC node activation during transition node quiescence.

The different time courses of PFC nodes encoding information at different levels of the task hierarchy demonstrate the sensitivity of our representations to aspects of the task which vary at different timescales. The features of these representations and their activation time courses are functionally equivalent to schemas at different levels of the IAN schema network (16). The dynamics of PFC nodes (figure 9a) is comparable with the schema activation profile from the IAN model of coffee making within the contention scheduling system (figure 5 from Cooper & Shallice, 2000 (16)).

The primary functional difference between the RoSe-BaL and the IAN model is the top-down flow of activation in the IAN model from higher to lower-level schemas. In the RoSe-BaL model all features, akin to low level schemas, are mutually supportive to create stable ‘high level’ schema-like representations in PFC. The PFC nodes encoding actions may be considered cognitive representations of action schemas; our channels in the motor loop, conversely, are motor representations of the same schemas. Thus, activation of motor representations of action by cognitive ones is analogous to the flow of activation from higher to lower-level schemas in the IAN model.

The overall dynamics of our PFC network are functionally equivalent to those of Botvinick and Plaut’s (25) SRN hidden layer, which itself encodes temporal task context. We visualise this equivalence using multidimensional scaling (MDS) analysis, the same approach they adopt for the analysis of their hidden layer. MDS extracts from a multidimensional dataset (here, the 49-dimensional PFC representation at each timepoint) two abstract dimensions which preserve most of the information encoded by the data at each point. The subsequent two-dimensional dataset may then be easily visualised. We performed an MDS analysis of the output of our PFC nodes over the course of a tea trial, and a milk-first coffee trial. Results of this analysis are displayed in (figure 10). These graphs show the trajectory of PFC activation in the extracted two-dimensional state space in each case.

**Figure 10:**
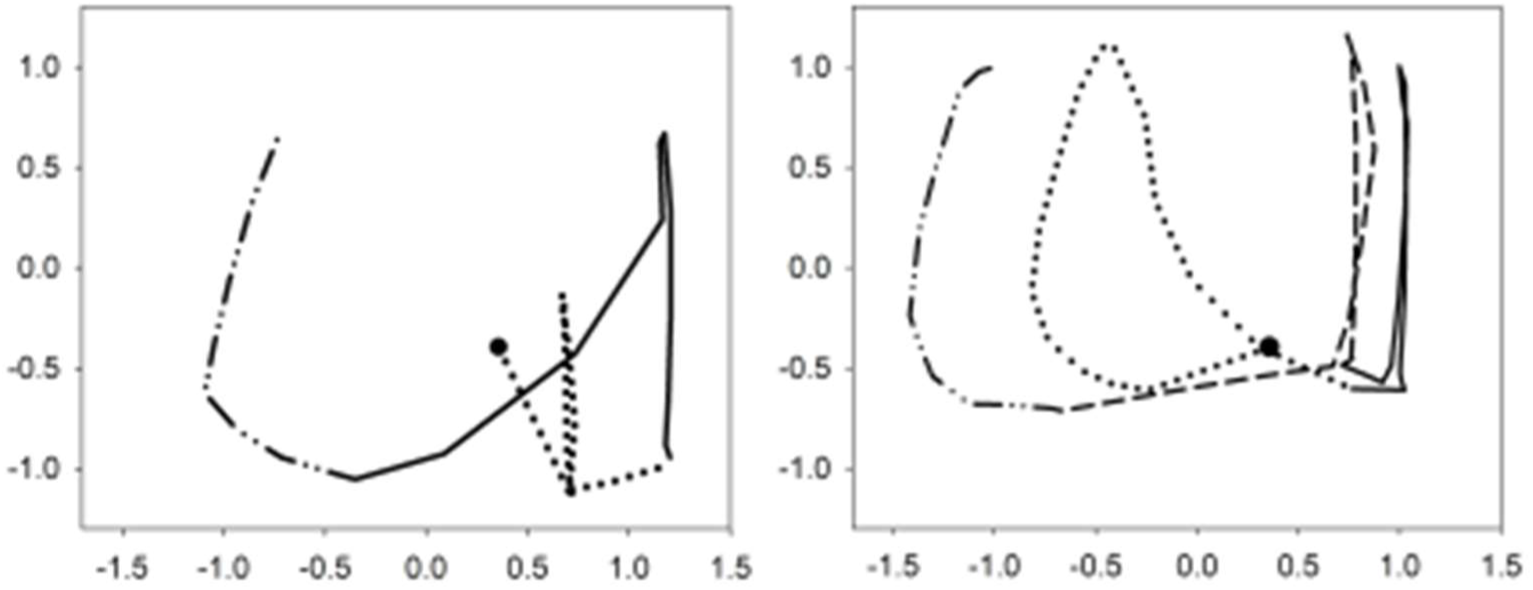
Trajectory through 2D ‘state space’ taken by the model for tea (left) and coffee (right) trials. Tasks begin at the circle, traversing the first subtask (dotted lines; teabag/coffee subtask), followed by the water subtask (solid line). The coffee task then performs the milk subtask (dashed line). Finally, both tasks perform the sugar subtask (dashed-dot line). This emphasises the similar trajectories taken by the model for different instances of the same subtask (e.g., sugar subtask). Axes represent abstract dimensions extracted by MDS analysis.

Since the PFC representations in the RoSe-BaL model were hand-coded, we can make inferences about the semantic underpinning of the similarities and differences in the trajectories from the two tasks. For instance, note the similar pronounced ‘uptick’ components of the trajectories from around –1 to 1 on the y-axes, during the milk and coffee subtasks (red; solid & dashed) and water and sugar subtasks (both red and blue, dotted and dot-dashed). This uptick feature is not shared by the tea subtask (blue, solid), suggesting it is driven by the ‘pour’ action, which is shared by all other subtasks. Note also the similarity between trajectories of the coffee and sugar subtasks; driven by the identical sequence of actions (pick up, scoop, pour) and object (spoon) not shared by other subtasks.

Due to the emergent nature of their representations, Botvinick & Plaut (25) could not make a similar interpretation of their equivalent analysis. However, our results demonstrate a functional equivalence of the representations encoded by the activity of our PFC nodes and by the hidden layer of the SRN adopted by Botvinick & Plaut, in terms of the encapsulation of key similarities and differences between variations of a task. It is important to note that this was observed despite the more explicit degree of localism in our representations, which was of a similar nature to that employed by Cooper & Shallice (16).

##### Significance for understanding neural data

A result of implementing feature-based representations is the natural overlap of representations that share features. Consequently, representation selection will effectively result in the partial activation of all representations that share features with the activated one. The more features a representation shares with the selected one, the higher its own partial activation will be. Figure 11a illustrates representations T1-T7 (the seven stages of the tea task). 11b shows the *average* output of *all* the nodes comprising each representation during each stage of a tea task (the total activity over all nodes for a particular representation, divided by the total nodes composing it). Naturally, each representation’s activation peaks during the stage of the task that it encodes. Notice that, from task initiation (stage 1), all representations are partially active (because all representations share the same active goal). Moderate activation is often observed adjacent to the point of peak activity for a given representation: representations generally show a gradual increase and then decrease in activation during the preceding and subsequence stages of the task. This is particularly evident for representations T1, 3, 6 and 7.

**Figure 11:**
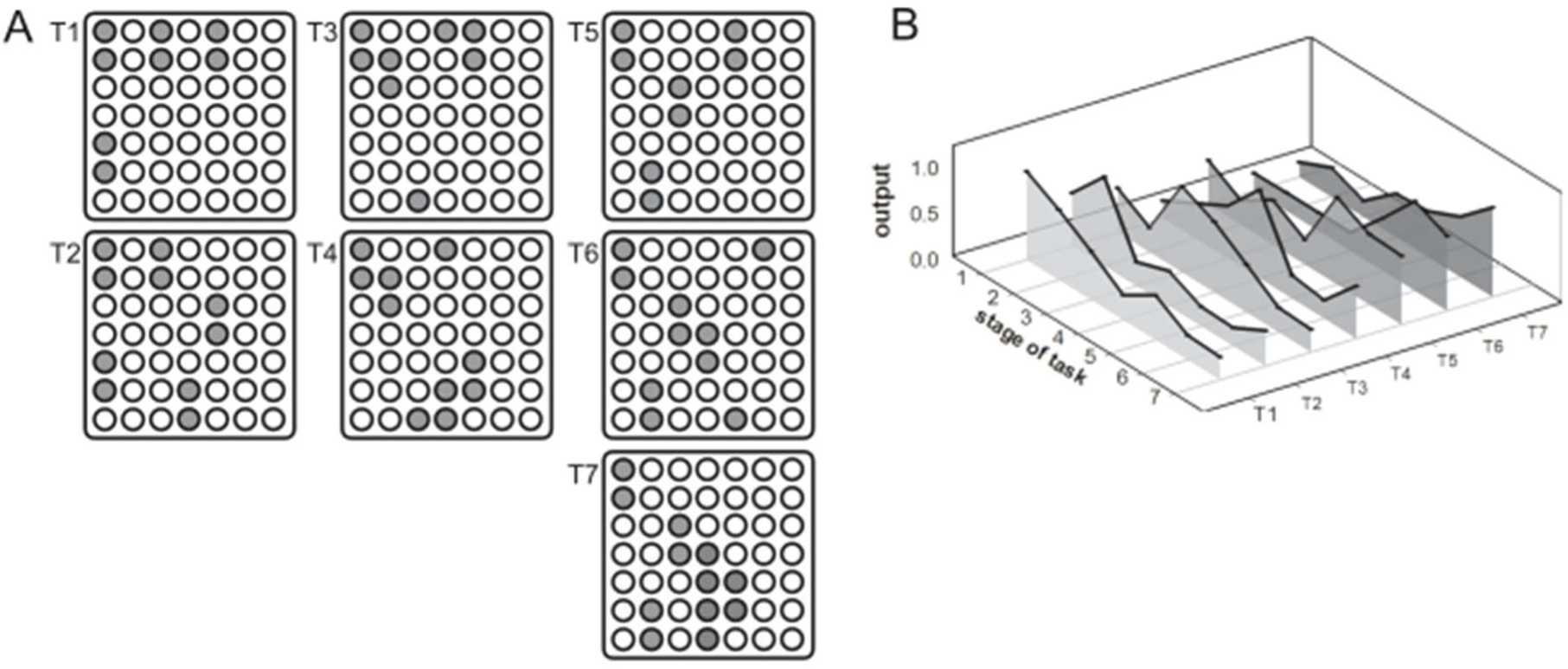
A) PFC representations for the tea task. Grey nodes denote PFC representations T1-T7, encoding each stage of the tea task (see table 1). B) Average output for these PFC representations for each stage of a successfully performed tea-making task. Note that all representations show a degree of output during the entire task, due to overlapping features in the representations.

This profile resembles activation of neural ‘ensembles’ in macaque PFC during a sequential drawing task (figure 2 from Averbeck et al., 2002 (22)). Averbeck and colleagues interpreted the early activation of each ensemble as ‘preparatory’ activation of each representation required for the task. Our study leads to an alternative possible explanation for this finding: that it may be a consequence of partial activation of representations, resulting from overlap with the currently selected representation. Here, the apparent gradual onset and offset of each representation over multiple task stages is a result of greater overlap between representations that encode adjacent stages of the task. This lends support to our interpretation of PFC representations as being feature-based. The preparatory-activation interpretation indicates that multiple possible mechanisms may give rise to these trends. It would be of interest for future work to attempt to distinguish these possibilities.

#### ii. Waiter Scenario

A distinction between the IAN and SRN models was the type of representation used to encode contextual information. The SRN model emphasised the power of distributed representations to encode key similarities between different versions of a task. The authors claimed this posed specific difficulties for the IAN model, since such variations must be represented by independent schemas. Botvinick & Plaut (25) argued that the schema network would be unable to properly or usefully capture the contextual representations for the performance of ‘quasi-hierarchical’ tasks, giving the example of a waiter making several coffees with different amounts of sugar. They presented a simulation based around this task: successful performance required the coffee task to be completed with zero, one, and two sugars. We replicated this simulation to examine the RoSe-BaL model’s ability to perform different variations of a task. We designed three alternative sets of representations for our tea task: each set included a distinct intermediate-level goal node indicating the number of sugars to be added, in addition to the overall goal node. For the two-sugars version of the task, additional rank order nodes were included to indicate the sugar adding task history. Additional context nodes were included to help signify the end of the task for the zero– and one-sugar versions of the task. These nodes separated heavily overlapping representations and helped resolve ambiguity in transition nodes (in a larger network, more nodes encoding the basic features would increase the distance between representations and likely render such additions unnecessary). The RoSe-BaL model was consistently able to complete all three versions of the task without error, according to the specific instruction given.

##### Analysis

Botvinick & Plaut (25) argued that the waiter task showed the insufficiency of schema-based accounts, since the distinct ‘goal nodes’ required by a schema network fail to capture contextual similarities between the tasks. However, our semi-localist, schema-like implementation successfully captures this similarity. MDS plots in (figure 12) illustrate this, showing that the model traverses similar trajectories through state space in each version of each subtask. In particular, trajectories for the add-teabag subtask are almost identical in each version of the task. Slight differences result from the activation of different intermediate goal nodes. In the ‘no sugar’ version of the task, a shorter trajectory is seen for the water subtask, reflecting the absence of a switch into the sugar subtask. Similarly, the trajectory of the sugar subtask is longer in the ‘two sugars’ version of the task, and ‘doubles back’ on itself, reflecting the repetition of the same actions for the second ‘add sugar’. The important contextual similarity is captured here, despite the localised representation of goals. Indeed, this functionality is present *because* of the semi-localist representation used here, and exists due to the overlapping nature of the representations.

**Figure 12:**
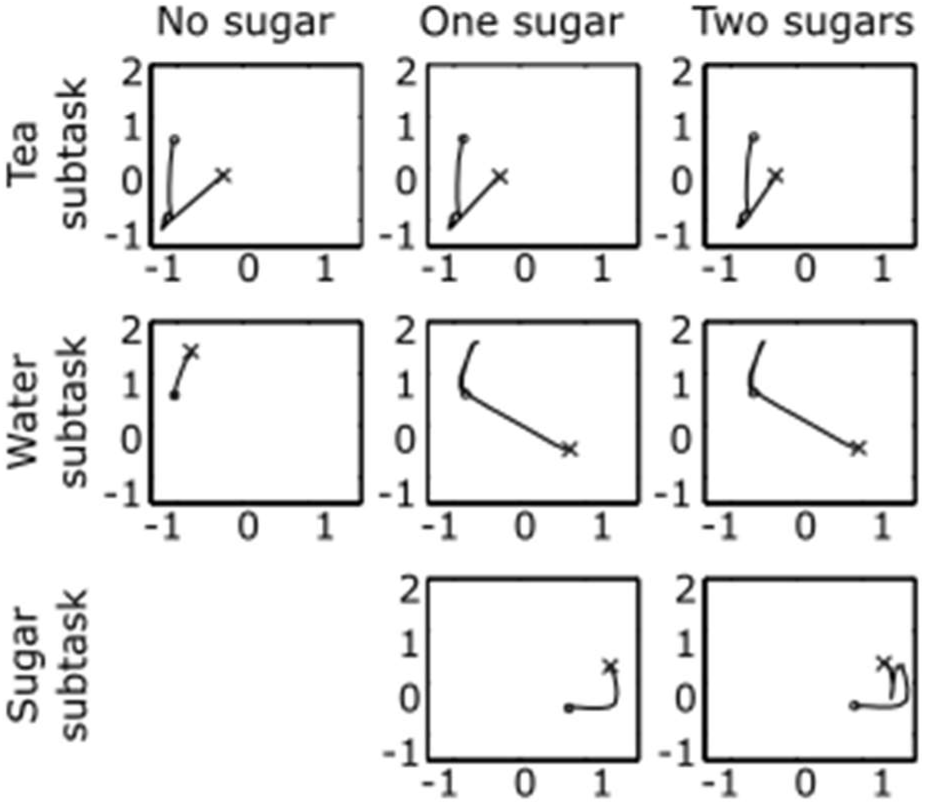
MDS plots of the trajectory of PFC through each stage of each subtask in the current model. Similar trajectories are observed for each version of each subtask. Left-hand columns show activity for the no-sugar versions of the task, middle columns for one-sugar, and right columns for two-sugars. Circles and crosses indicate the beginning and end of each subtask, respectively. Compare with Figure 8 from Botvinick & Plaut, 2004 (25))

Comparing Figure 12 with equivalent MDS plots produced by Botvinick & Plaut further illustrates the functional equivalence of the dynamics of the two models. The differences in the trajectories in Botvinick & Plaut’s SRN model are likely to reflect the same information as that encoded by our distinct goal and context nodes. It is possible that the SRN model also learned to represent the distinct ‘goals’ separately on each version of the task. This suggests that the explicit hierarchies of schema-based models and the implicit hierarchies embodied in the distributed representations of SRNs are functionally equivalent, with both able to capture the similarity of related tasks.

### 3.6. Discussion

The novel ‘subset-selection’ scheme within the RoSe-BaL architecture can mediate the successful performance of two highly similar but distinct tasks, with complex demands on sequencing and selection.

#### Neurophysiology of PFC ‘features’

The functionality of PFC in the RoSe-BaL model is consistent with previous discussion about the role of PFC in maintaining representations of goals and strategies (45). The orthogonal representation of features in PFC is consistent with evidence for prefrontal cells selective for distinct stimuli (47–49). Notably, neurons encoding ‘rank order’ of stimuli have been documented (57, 109, 110). Such neurons have been the focus of existing sequence learning models (138). These neurons were critical for flexible sequencing in our model and may have been documented frequently due to their vital importance in mediating sequencing behaviour. A prediction arises from our model that such neurons would only be observed in studies where the temporal order of stimuli or actions is variable. Where the order of stimuli or actions is fixed, no explicit sensitivity to temporal order is required, as sequence information is implicit in the stimulus or action itself.

#### Transition layer & sequencing

Sequencing mechanisms have caused difficulty for previous models, which often rely on mechanisms such as inflexible lateral inhibition (108), or recurrent dynamics (25, 104) which cause problems for sustained activation. Our solution to this, the transition layer, may be considered to encode the feature of ‘sequence knowledge’. This represented a flexible, context-sensitive approach to sequencing which allowed sustained activation in PFC. Its separation from PFC itself was important for the stability of representations; indeed, a computational imperative exists for a mechanism external to PFC for controlled sequencing of stable representations (137). Our model predicts that such a separation might be found in the brain. Distinct PFC regions and layers have been shown to be selectively involved in maintenance or selection (139, 140). Here, the transition layer may perform a more selective function, whereas the associative loop itself is more critical for maintenance. This specialisation in PFC may occur as a result of the computational incompatibility of internal sequencing and stability (137).

#### Sensory + cognitive convergence

Where cognitive influences reflect intention to act, sensory influences in the form of affordances reflect action salience guided by object knowledge. The convergence of these is necessary: an affordance may not be sufficiently strong to trigger selection independently, requiring additional top-down support. Additionally, multiple affordances will require contextual disambiguation if they are activated with similar strength. This convergence is consistent with evidence suggesting that damage to the ventral visual stream can result in the inability to use object knowledge to guide action, suggesting damage to affordances (141). Similarly, certain PFC lesions result in ‘utilisation behaviour’ where action is guided by external stimuli rather than intention, suggesting a loss of top-down influence (142). Deficits like these would be expected from the RoSe-BaL model, which makes explicit the functional relationship of these two pathways and are explored in section 4.

#### Corticostriatal sparseness

Our corticostriatal projection between loops is sparse in that it arises only from those particular nodes encoding action. This is consistent with evidence that inter-territory corticostriatal projections are less widespread than those within loops (131). This may be because only certain categories of information need to be propagated through different functional territories for goal-directed action; in this case, only those critical for motor selection itself. This sparseness would be unlikely if prefrontal representations were fully distributed, as the entire representation would be important for encoding action. This observation thus provides further support for our feature based or semi-localist organisation of PFC.

#### Reconciling existing models

Having used neuroanatomical constraints on model development, the RoSe-BaL model has naturally reconciled aspects of the two existing competing models (16, 25), within an anatomically constrained framework. RoSe-BaL is thus a ‘hybrid’ model displaying some functional equivalence to both: the necessary structure of the representations needed for the task gave rise to activation profiles similar to those presented by both sets of authors. Both sets of authors do acknowledge that their differences may be reconcilable, with Cooper & Shallice (32) stating,

> ‘[the networks] may be reconciled through the mapping of nodes within the interactive activation network to discrete point attractors … one can be optimistic about the development of a model which functions at one level according to the principles of Botvinick and Plaut, and at another according to our … principles’ (p906).

This is precisely what the RoSe-BaL model achieves. This reconciliation, however, has emerged naturally from the implementation of a neurally constrained architecture. It follows that the inscrutable nature of the emergent representations in the SRN model may obscure the likely fact that the same key information is represented in both models. In particular, the representations of ‘goals’, which provoked much debate between the two sets of authors, are functionally equivalent to those of overall ‘schemas’, where both may be regarded as features or combinations of features within the overall representation of context. While this possibility was accepted in the debate, it is demonstrated concretely in our model. We have therefore shown that it is possible to capture contextual similarity using schemas; this emerges from overlapping representations despite the localist representation of features. As Cooper and colleagues (32, 72) point out, the IAN model effectively incorporates overlapping representations where higher level schemas make use of the same subschemas for different tasks. Notably, evidence suggests that neural representations are unlikely to be fully distributed (84–86), making semi-localist representations both plausible and functionally appealing.

## 4. Modelling III: Error commission

Much research exists on the patterns and occurrence of human error commission during sequential performance, both in healthy volunteers (143, 144) and patient populations (5, 27, 30). Much of this has focused on errors made during routine tasks described as ‘activities of daily living’ (ADLs); of which well learned sequences such as tea– and coffee-making are common examples. Models of sequential processing have been examined under a degree of disruption in order to examine their ability to account for patterns in these error data: this was a focus of the IAN (16) and SRN (25) models. Here, we show how the new dual-loop model can account for a range of error commission data.

### 4.1. Experimental findings

Diary studies examining action slips (143, 144) indicated that healthy participants commonly made errors during routine tasks. These tended to be perseverations, omissions or intrusions of well-learned sub-sequences, and generally occurred at natural branch points in performance. Object substitutions were common, where an action was performed with the wrong object. In brain injury patients, distinct patterns of errors are seen. Sufferers of action disorganisation syndrome (ADS) show disjointed sequential behaviours, often displaying ‘single action errors’, contrasting with the tendency in healthy people to make errors at the level of full sub-sequences. Single action errors may manifest as omissions of single actions, or the performance of independent actions: those performed outside the boundaries of clear sub-sequences. Independent actions include ‘toying’ behaviour, where objects may be aimlessly picked up and put down (29). Perseverations and intrusions of single actions and sub-sequences, and object substitutions, are also observed in ADS sufferers.

Various explanations have been offered to account for these findings, many of which revolve around the disruption or faulty activation of schemas (15, 27, 28). Two main hypotheses have been examined in computational modelling work. The first, that ADS results from an imbalance of top-down and bottom-up influences on the activation of action schemas (28) was tested with the IAN model (16). The second, that it results from a general deficit in cognitive resources (5), was examined by using the SRN model (25, 29). These models reproduced a number of features typical of human error commission lending support to the hypotheses, though both models failed to replicate certain error types, and neither focused on neuroanatomical or physiological explanations for their findings.

Evidence from patients suggests that the latter ‘resources deficit’ account is insufficient to account for errors in ADS (145). An alternative hypothesis which has received less attention suggests the disruption of temporal order knowledge, and/or the breakdown of stored schemas encoding component actions is a main cause of errors in ADS (27, 31, 145, 146). In experimental work, patients showed both impaired action and order knowledge (147), but the authors suggested a dissociation between these processes was possible (146, 147).

The architecture of our model lends itself well to testing these hypotheses, given the functional separation of action schemas in PFC, and temporal order knowledge, encoded in the projections between the transition layer and PFC. In the following simulations, we test the breakdown of these processes, to examine whether their disruption may be responsible for action slips or the degraded performance seen in ADS.

### 4.2. Classifying errors

Errors have proven difficult to classify in previous work (5). Observational and modelling studies have used slightly different systems to describe and classify errors. Here, we attempt to define a system which classifies errors in a way that allows comparison with as much of the existing work as possible.

1. Omissions. An omission occurs whenever an item is not added to the cup. This may the omission of a full subtask. Alternatively, if for example, the ‘pick-up kettle’ action is performed, but the ‘pour-into cup’ action is not, this is counted as an omission. (In this case, the ‘pick-up kettle’ action would be further classified as an independent action.)
2. Sequence errors. Previous work has defined errors inconsistently. Some accounts include ‘anticipation omission’ errors (5, 27); other accounts do not (16). Perseverations are included by some (5) but not others (27). Some include atypically early performance of a subtask (16, 27), others do not (5). Due to this wide variation, we examine our results in terms of different definitions where appropriate.
3. Additions. These are also inconsistently defined, sometimes including errors that could be considered object substitutions (25), or anomalous actions (5). We only include intrusion errors in this category, where an action or subtask belonging to a different task is performed.
4. Semantic errors. These consist primarily of object substitutions: actions that are appropriate in context but performed with the wrong object. We also include object omissions, such as stirring coffee without a spoon. Although some previous work (27) has classed object omissions separately, we see this as a distinct class of substitution and thus a semantic error.
5. Quality/spatial errors. These consist of actions involving inappropriate quantities or positioning. Since our model currently includes no representation of space or volume, we do not examine these errors in our analysis. While our model can produce certain errors previously classed as quality errors (pouring cream four times in a row; (25), we consider this a perseveration error.
6. Subtask-based vs single action errors. Subtask-based errors consist of omissions, perseverations, reversals or intrusions of entire subtasks. Single action errors consist of omissions of single actions, object substitutions, or the performance of independent actions outside of clear subtasks. Independent actions may be further categorised as perseverations, intrusions, reversals, or anticipation-omissions.

#### Detection and quantification

In the following simulations, where sequences contained only subtask-based errors, these were extracted by an automated process. To examine the prevalence of single action errors, we examined a random sample of 100 trials at each noise level by eye (25). This approach was taken due to the complexity of model behaviour at high noise; automated classification of the resulting errors resulted in some error misclassification.

### 4.3. Error simulation i: Disrupting sequence knowledge

First we sought to test the hypothesis that disruption of temporal order information alone could account for errors in routine behaviour (27). We added noise to the transition nodes, increasing the probability of erroneous sequencing via spurious activation of incorrect transition nodes. Noise was drawn randomly from a normal distribution with mean μ=0. We tested the model at 9 levels of noise. These levels corresponded to the standard deviation of the noise distribution, which took values of σ=0.01, 0.02, 0.05, 0.1, 0.2, 0.3, 0.5, 0.75, 1. 200 trials were run for each of the tea and coffee tasks at each level of noise, giving 3600 trials in total. Noise was added to the activation *a* of each transition node at each simulation timestep.

#### Results

Figure 13 (top) shows the number of correctly performed trials for each level of noise. The model is fairly robust, with substantial numbers of errors only occurring at σ ≥ 0.2, and increasing numbers of errors appearing gradually with increasing noise. Examples of typical erroneous sequences seen and our error classifications at various levels of noise are given in Table 1.

**Figure 13:**
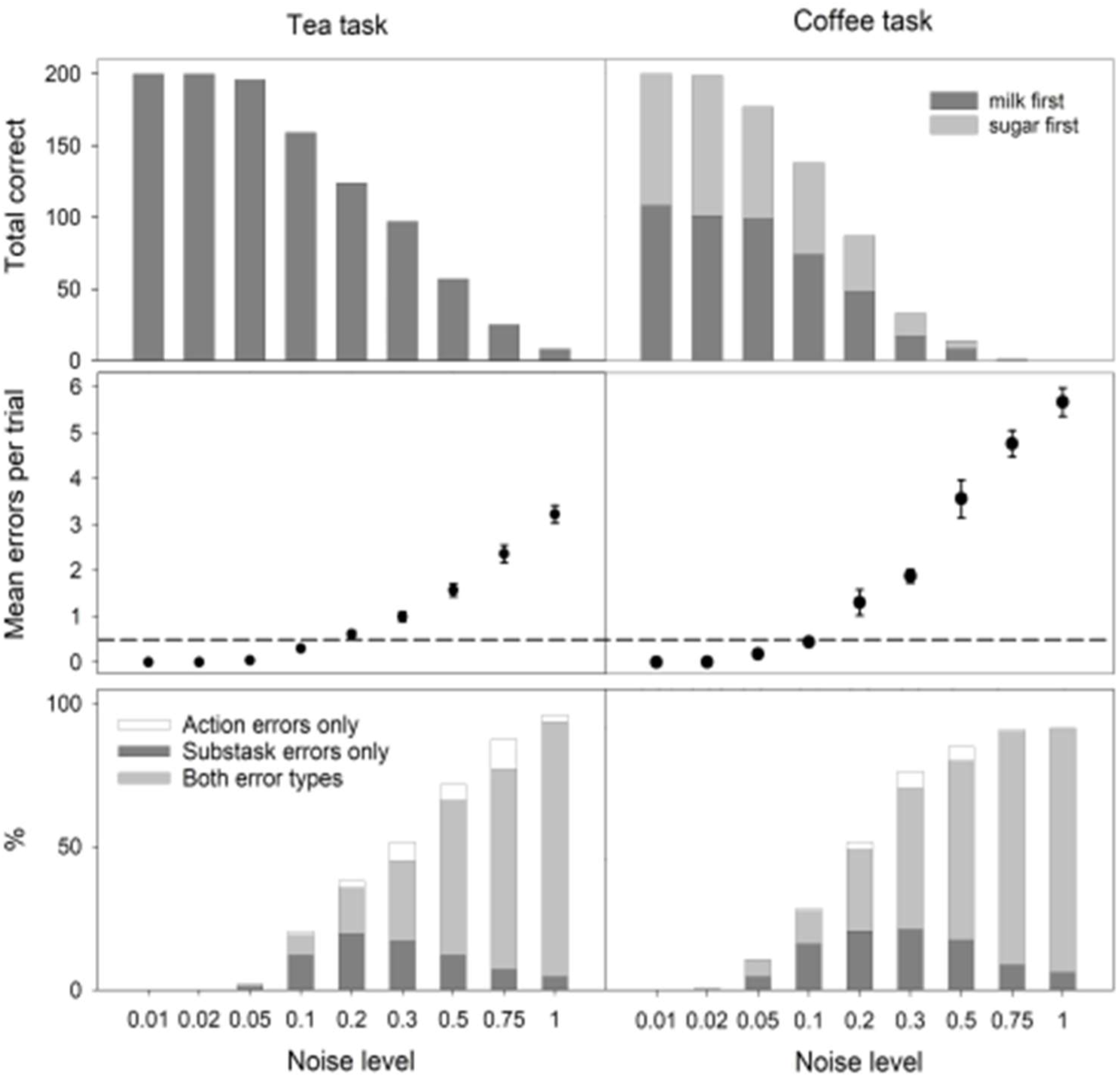
(Top) Overall model performance shown as the number of correctly performed trials at each level of noise for both tasks, out of 200 trials run at each noise level. (Middle) Mean errors per trial and sample standard error at each level of noise. Dashed line shows threshold for classification as impaired performance. (Bottom) Percentage of trials displaying subtask-based errors only, subtask based and single action errors, and single action errors only.

**Table 1.** Correct coffee-making sequence (σ= 0) compared with erroneous sequences at low, medium and high noise. σ = 0.2: includes the abandonment of the sugar subtask (underline), giving an omission error and an independent action (spoon picked up but not used). σ = 0.5: the sugar subtask is executed successfully but includes four perseverations (bold). The coffee subtask is not completed (underline), giving an omission error and an independent action. The milk subtask is omitted, giving a second omission error. σ = 1: Both coffee and sugar subtasks were completed in a disorganised manner, with the former including perseverations (bold), and the latter achieved through an object substitution (italics), using the kettle to scoop the sugar. Again, the milk subtask is omitted.

| $\sigma=0$ | $\sigma=0.2$ | $\sigma=0.5$ | $\sigma=1$ |
| --- | --- | --- | --- |
| Pick up spoon | Pick up spoon | <u>Pick up spoon</u> | Pick up spoon |
| Scoop coffee | Scoop coffee | <u>Scoop coffee</u> | Scoop coffee |
| Pour In cup | Pour In cup | Pick up kettle | **Pick up spoon** |
| Pick up kettle | Pick up kettle | Pour in cup | **Scoop coffee** |
| Pour in cup | Pour in cup | Pick up spoon | **Pick up spoon** |
| Pick up spoon | <u>Pick up spoon</u> | **Pick up spoon** | Pour In cup |
| Scoop sugar | Pick up milk | **Pick up spoon** | Pick up kettle |
| Pour in cup | Pour in cup | Scoop sugar | Pour in cup |
| Pick up milk |  | **Scoop sugar** | *Scoop sugar* |
| Pour in cup |  | **Scoop sugar** | *Pour in cup* |
|  |  | Pour in cup |  |

#### Normal action slips

For consistency with previous work (25), we divided the results into categories of normal and impaired performance based on the mean number of errors per trial; performance was impaired if more than 0.5 errors per trial were observed on average (figure 13, middle), based on a sample of 100 trials at each level of noise. Noise at σ ≥ 0.2 produced impaired behaviour for both tasks.

Reason’s diary studies (143, 144) indicate that subtask-based errors, or ‘action slips’ at sequence branch points, are dominant in normal error-making, whereas single action errors are more likely in patient populations (28). Thus, where performance was classified as healthy, we expected to see predominantly subtask-based errors. Indeed, this was observed. Figure 13 (bottom) shows the distribution of subtask-based and single action errors across noise levels. Trials containing only subtask-based errors comprise around half of all erroneous trials up to σ = 0.2. With more noise, this fraction decreases as behaviour becomes more disjointed. Patterns of errors were stereotyped in healthy performance, with repetitions and omissions of the sugar subtask and additions of the milk subtask being very common. Repetitions of whole subtasks in healthy performance tended to be recurrent, where a subtask was repeated after an intervening subtask (148).

#### Mechanisms underlying action slips

“Contextual confusion” was the primary mechanism in the model for most subtask-based errors. This is similar to that described by Botvinick & Plaut (25). Here, noisy transition node activity accounted for ‘confusion’ as follows. Each of the 22 legitimate task stages (Table 1) had a corresponding transition node providing excitation to PFC. In a successful trial we see consecutive activation of the task-relevant transition nodes as the model progresses through the task (figure 11b). Transition node activity during a trial in which an intrusion error occurred (milk added to tea) is illustrated in figure 14. This shows the expected pattern of activity for the first 5 stages of the task. However, where we should see activation of transition node T6 (triggering a ‘scoop-sugar’ action), instead we see activation of transition node C7a from the coffee task, which further activates representation C7a in PFC. Note that transition nodes T6 and C7a activate distinct PFC representations which *both* drive a ‘scoop-sugar’ action. This means, at the behavioural level, no error is yet apparent. However, T6 and C7a have different overall representations of *context*. Thus, there is a delayed effect on outward behaviour: the sugar subtask is completed as normal, but then the sequence is continued *as if it were the coffee task*. The milk subtask is subsequently performed, and an addition error occurs. This is analogous to ‘forgetting’ the goal of tea-making. As observed by Botvinick & Plaut (25), the *cognitive* error occurs some time before the outward *behavioural* error. Notably, the distraction of normal participants during the performance of routine subtasks is more likely to result in error than distraction between subtasks (149), in line with these results.

**Figure 14:**
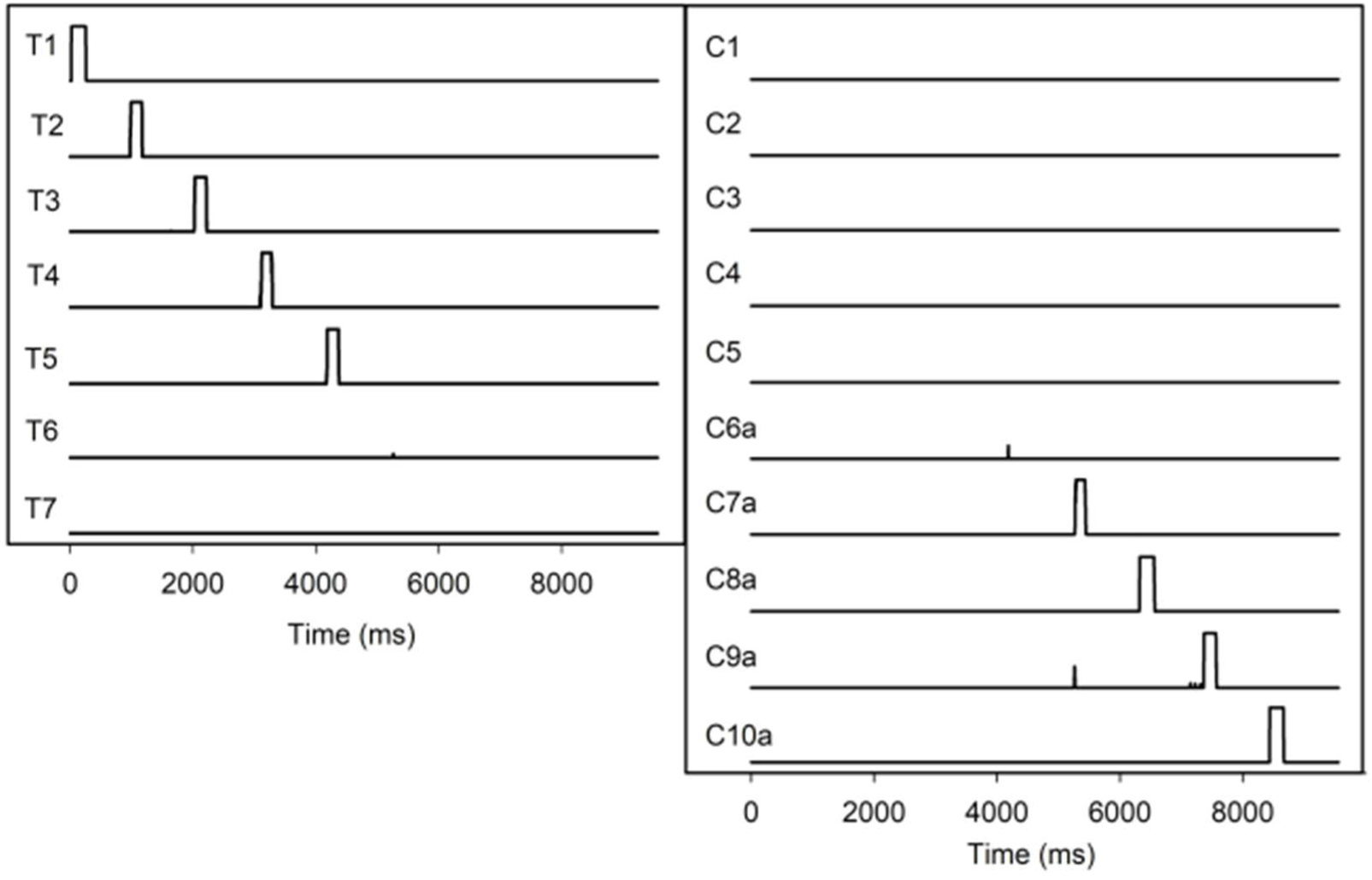
Activity of transition nodes T1-T7 (left) and C1-C10b (right) during a tea-making trial in which milk was added. Noise causes erroneous activation of transition node C7a rather than T6 (∼5500ms), triggering a contextual representation from the coffee-making task. This causes a contextual ‘switch’ to the coffee task, and the model proceeds as if performing the coffee task. This phenomenon is a direct result of the feature-based nature of our PFC representations. An active representation preferentially excites a single transition node. However, the feature-based organisation of PFC means that varying levels of subthreshold activation occurs across several transition nodes. In the present case, representation T5 preferentially activates transition node T6. Similarly, representation C6a preferentially activates transition node C7a. However, significant overlap between representations T5 and C6a means that representation T5 partially activates transition node C7a. Since the transition layer approximates a WTA network, a small amount of noise added to this region is sufficient to drive the erroneous activation of transition nodes C7a, rather than the correct T6, resulting in the observed behavioural error.

#### Pathological errors

At levels of noise resulting in impaired performance, we saw frequent performance of disjointed, independent actions. This is an important component of ADS. Schwartz and colleagues (28, 150) described two ADS patients, HH and JK, who performed up to 31% and 16% independent actions that were not clearly contributing to the assigned task. We saw similar fragmentation, with independent actions accounting for 15-33% at the two highest levels of noise. In patients, the proportion of independent actions was variable across tasks, and for patient HH, was reduced with practice and recovery, suggesting an increasing vulnerability to fragmentation with increasing impairment. Correspondingly, we saw a general increase in independent actions as noise increased (figure 15). The error-making trends at impaired levels in the model are also reflective of the experimental data: figure 16 shows the overall trends in error rates across our two highest levels of noise for the four main error types, and their similarity to trends observed in patient data from Humphreys & Forde (27) and Schwartz et al. (5).

**Figure 15:**
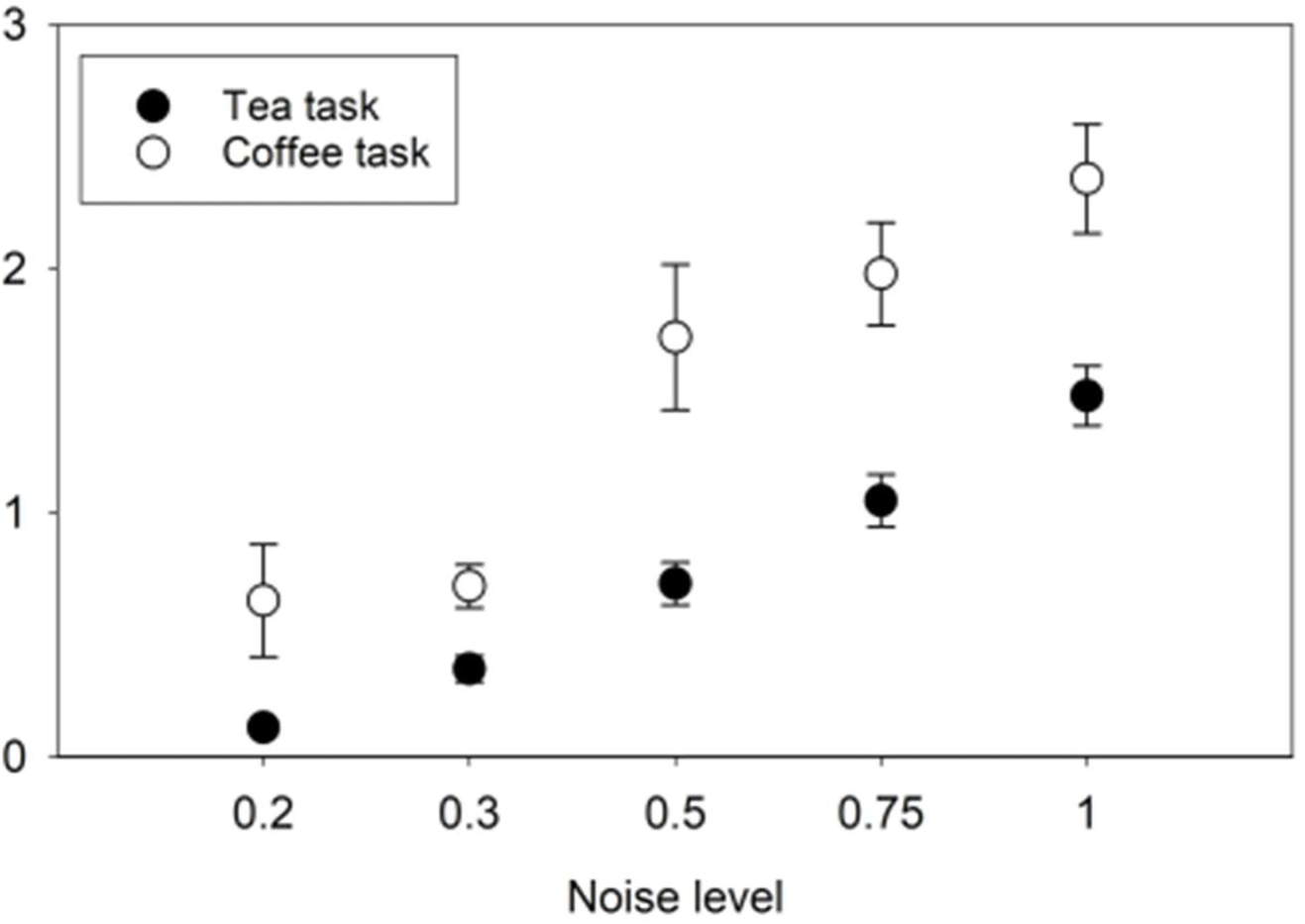
Mean number of independent actions per trial, based on a sample of 100 trials at each noise level. The increasing number of disjointed actions with increasing noise reflects the trend documented by Schwartz et al. (1991), suggesting increasingly fragmented behaviour with increasing impairment. Error bars show sample SE.

**Figure 16:**
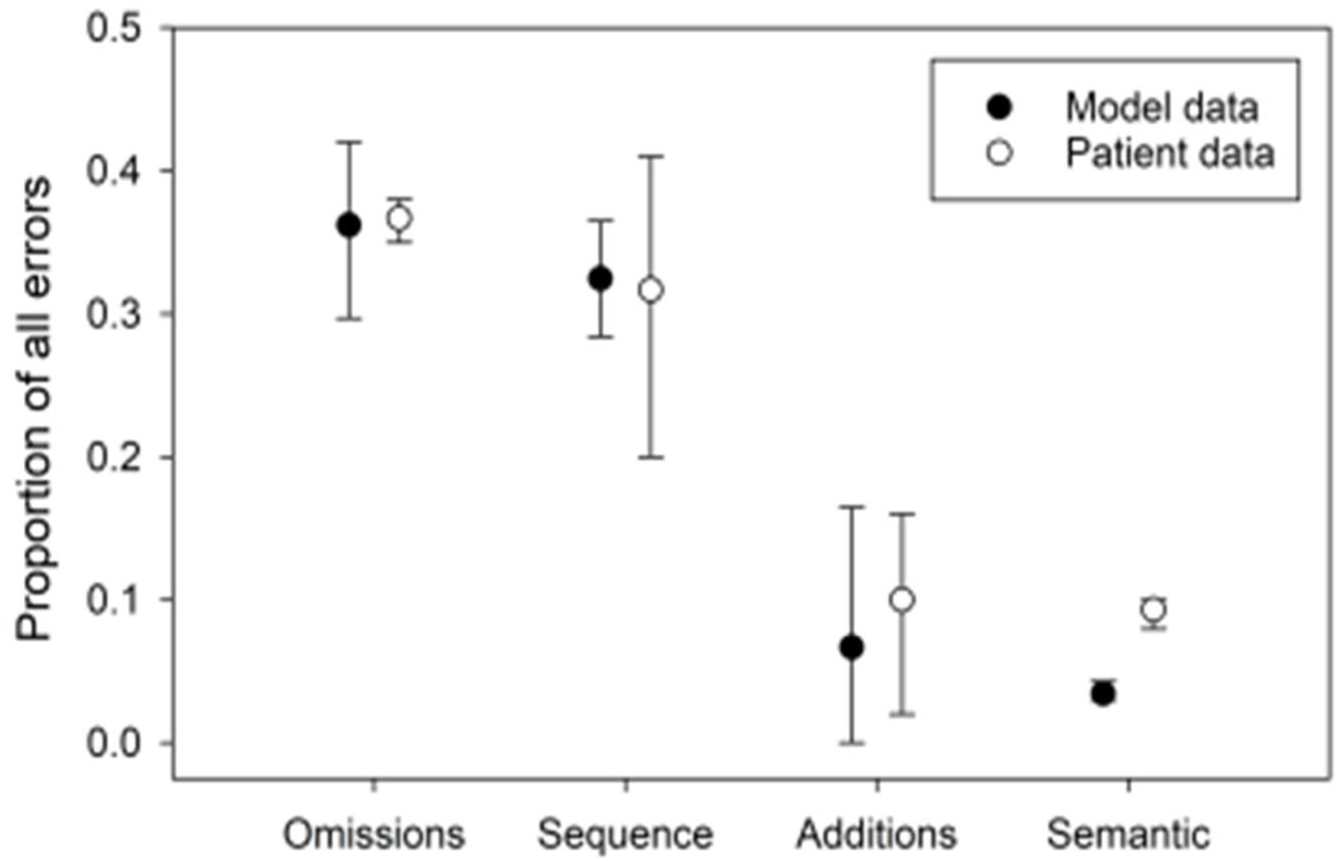
Summary of error types as a proportion of all errors. Black circles: grand mean of average error proportions for tea and coffee trials over the two highest noise levels. White circles: average of reported data from patients FK, HG, and CHI patients reported by Humphreys & Forde (1998), and Schwartz et al. (1998). Error bars show the range of the averaged values.

#### Omissions

Omissions generally account for around 30-40% of errors in ADS, across labs, patients and tasks (5, 27, 30), and are generally the most common error type in ADS. Omission rates at high noise in our simulations showed a strong resemblance with rates observed in ADS patients (figure 16). Omissions also outnumbered any other error type at all noise levels that resulted in impaired performance. Dividing the number of omissions made by the number of subtasks gave the omissions *per opportunity* for an omission (Figure 17a). This suggests the coffee task was more prone to omissions, even accounting for its extra subtask. This was probably due to the flexible order of the milk and sugar subtasks, which generated more opportunities for confusion about the model’s action history, resulting in the model ‘forgetting’ having performed (or not) the sugar or milk subtasks.

**Figure 17:**
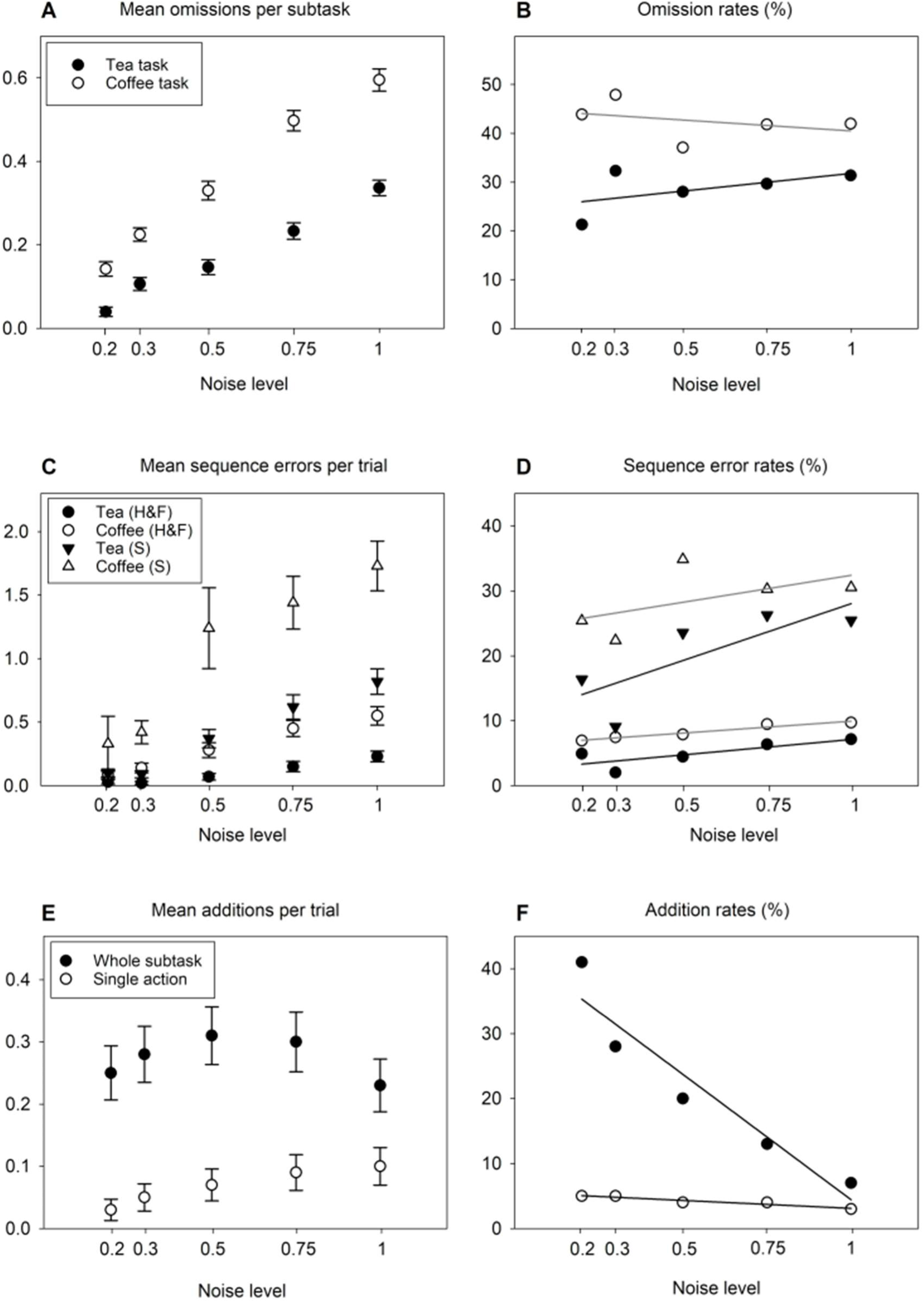
A) Average omissions per subtask in each task. B) Omission rates calculated as the proportion of all errors. C) Mean number of sequence errors per trial, using definitions of ‘sequence error’ used by Humphreys and Forde (circles) vs Schwartz (triangles). D) Sequence errors as a proportion of all errors. E) Average number of subtask based and single action additions per trial, for the tea task only (no additions were observed in the coffee task). F) Additions as a proportion of all errors. Error bars show sample SE; regression lines indicate trends for increasing/decreasing error rates.

While the total omissions increased with noise for both tasks, we found only weak (tea task) or no evidence (coffee task) for an increasing omission *rate* (figure 17b. This was unexpected given previous reports of a strong ‘omission rate effect’, whereby omissions tend to account for a higher proportion of all errors with increasing disorder severity (5, 31). This may be due to the task structure employed here, and un-intuitively, the relative opportunities for *intrusion* errors masking any omission rate effect in the coffee task as follows. Many erroneous tea trials at low noise involved intrusions of the milk subtask. Intrusions are less common in the tea task at high noise (figure 17f). As a result, omissions appear more dominant at higher noise in the tea task, since intrusions are rarely observed. However, since there is no equivalent opportunity for intrusion from the tea to the coffee task, intrusions are *not* observed at low noise in coffee trials, resulting in a possible over-representation of omissions in coffee trials at low noise and a consequent masking of any omission rate effect. Thus, including a suitable opportunity for intrusions might reveal an omission rate effect in the coffee task.

#### Sequence errors

Regardless of the definition used, the number of sequence errors per trial increased with noise (figure 17c). However, the definition significantly influences the sequence error *rates* observed (figure 17d). The definition of sequence errors given by Humphreys & Forde (27) includes anticipation-omissions and reversals but not perseverations. According to this definition, our model made sequence errors at an average rate of 6.7% and 9.6% for the tea and coffee tasks for the two highest levels of noise.

This is similar to the rates reported using the same definition, of 16% and 10%. When we categorise sequence errors according to Schwartz et al. (5), now including perseverations, we see rates of 25.9% and 30.4% for the tea and coffee tasks, compared with rates around 20% reported by Schwartz et al. using the same error definition. Notably, if perseverations are included in the sequence error rates reported by Humphreys & Forde (27), this increases to 34% and 41%. Clearly, perseverations tend to dominate sequence errors. This illustrates the importance of using consistent definitions to classify error types, and highlights potential problems with generalisations about sequence errors against which model data have been compared (25).

#### Additions

At high levels of noise, addition rates were similar to those reported in the previous literature. For the tea task additions accounted for around 13.4% of errors on average; Humphreys & Forde (27) reported addition rates of 16% for patient FK, though patient HG made fewer, at around 2%. Schwartz and colleagues reported 12% additions in their patient population (5). Subtask-based intrusions comprised a larger proportion of addition errors at low noise (figure 17f). There were no additions in the coffee task, because there were almost no opportunities for intrusion errors to this task. The only possible intrusion (the teabag subtask) is unlikely due to the lack of an appropriate context during the coffee task, again emphasising context confusion as the origin of most errors. Consistent with this, Forde et al. (146) found that intrusion errors were triggered by overall task context, but not sensory, ‘bottom-up’ interference.

#### Mechanisms underlying pathological errors

Like action slips or subtask-based errors, several single action errors are driven by a confusion of context. However, more noise is generally required for contextual confusion to drive single action errors, because fewer features are shared between the confused representations. This accounts for the more fragmented nature of errors at high noise; actions in different subtasks tend to have underlying representations which differ more than those within the same subtask, thus requiring more noise for confusion to take place. Furthermore, full subtask-based errors manifest less often with increasing noise because any misplaced subtask will be increasingly subject to errors within itself.

#### Disinhibition of transition nodes

Figure 18a shows transition node activity during a coffee trial in which the model perseverated the pour-water action. The outward behaviour suggests a faulty transition, causing the selected representation to be maintained too long, driving repetition. However, transition node activity reveals a different mechanism. Around t=5500ms, transition node C6b is (correctly) activated, which should activate the corresponding PFC representation triggering a pick-up-milk action.

**Figure 18:**
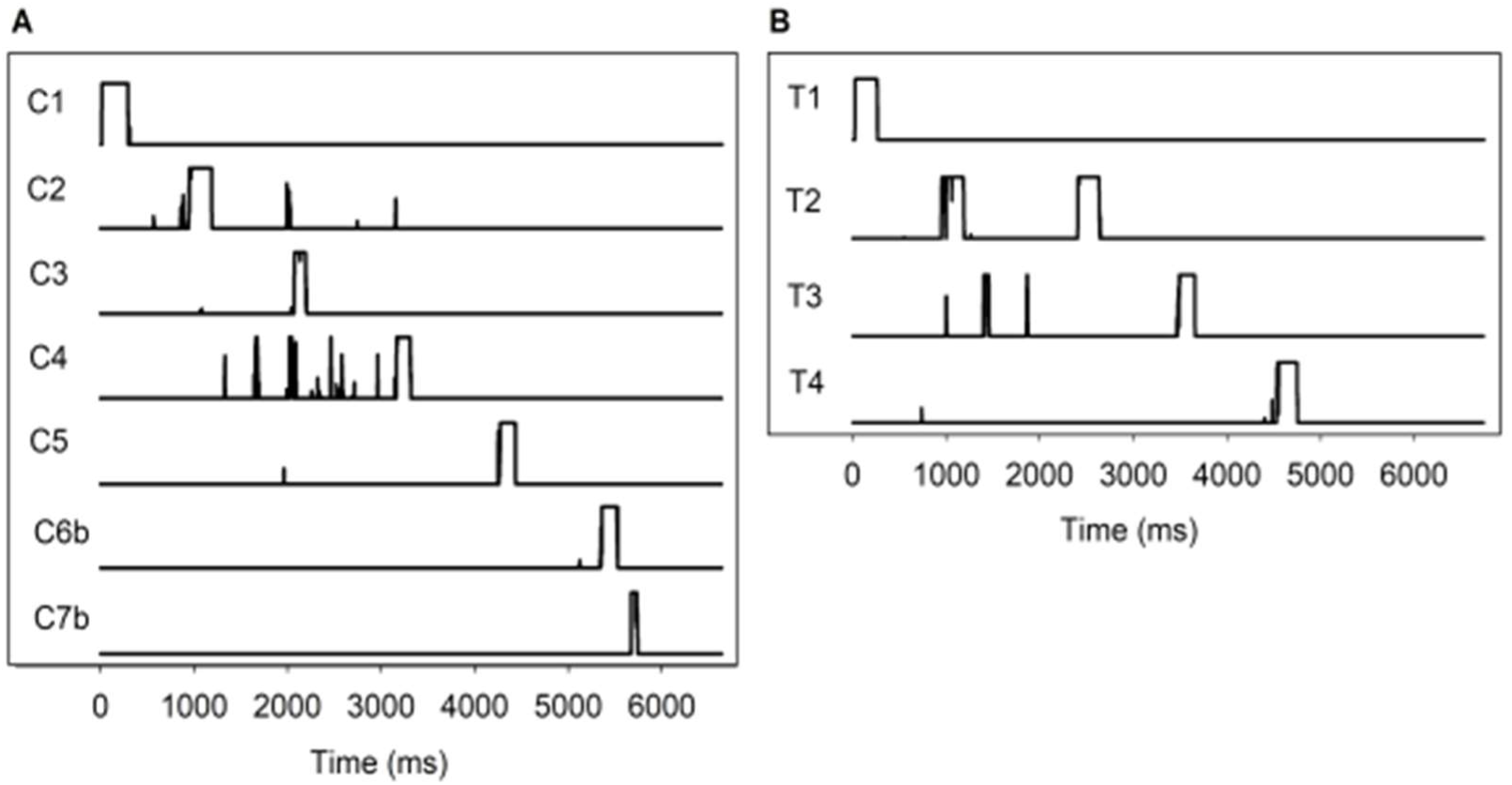
A) Transition node activity during a coffee-making task in which the pour-into-cup action was performed twice. Early activation of transition node C7b causes rapid switching in PFC, resulting in a cognitive-level omission of the action corresponding to C6b, and performance of the action corresponding to C7b. This manifests behaviourally as a continuous perseveration, since action C7b (pour into cup) is the same as action C5. B) Transition node activity during a tea-making task in which the pick-up-teabag action is performed twice. This ‘toying’ behaviour results from inefficient activation of representation T2, which itself is caused by interference from multiple transition nodes (1000-2000ms). This leads to a ‘decay’ back to the previously selected PFC representation T1, and a perseveration of its corresponding action.

However, noise-induced activation of transition node C7b at around t=5800ms causes a transition to the PFC representation C7b *before* the pick-up-milk action has been executed. The prematurely selected representation C7b triggers a pour-into-cup action, which is duly executed by the motor loop. Since action C6b – pick-up-milk – was not performed, the result appears behaviourally to be a perseveration of pour-water. This mechanism also resulted in several behavioural anticipation-omissions and object substitutions. Thus, the outward behaviour of the model is not necessarily representative of the underlying cognitive dynamics. To our knowledge, no previous modelling work has shown similar insights regarding this apparent discrepancy between cognition and behaviour.

#### Interference

Another mechanism for error is illustrated in Figure 18b. In this erroneous tea trial, the pick-up-teabag action was perseverated. This was caused by noisy activation of transition node T2 (put-(teabag) into-cup), and simultaneous activation of node T3 (pick-up-kettle). This caused interference in the signals sent to PFC, in turn causing representation T2 to be insufficiently activated to compete with the currently selected representation T1. Representation T2 is unable to reach the required stability, and the pattern of PFC activation decays back to the previously selected T1. It is interesting to note, that both this interference effect and the premature switching discussed above are both significant causes of immediate perseverations, which are indistinguishable at a behavioural level.

#### Summary

The present model can account for a variety of data on behavioural errors using only a single locus of disruption: noise in transition nodes. Analysis of the model dynamics underlying particular errors resulting from this noise has revealed a variety of mechanisms that are not clear from the observed error itself. This supports Humphreys, Forde and colleagues’ (27, 146) hypothesis that disordered sequence knowledge is responsible for many of the effects of ADS. Intriguingly, our data at high noise bore a particularly close resemblance to the error profiles of patient FK (27, 145), for whom the explanation of disrupted temporal order knowledge was originally suggested.

There were some important differences between our data and those from observational studies. We saw low rates of semantic errors, (not illustrated) and high numbers of additions at low noise. Due to the specific implementation of the model, we are unable to account for quality or spatial errors. However, there is potential in the model to explore the effects of disruption in additional loci, which may allow replication of more data. For instance, incorporating semantic properties of objects could feasibly increase semantic errors, particularly if distractor objects were included.

### 4.4. Error simulation ii: Disruption of schemas

We now examine Humphreys & Forde’s alternative hypothesis that disruption of action schemas themselves underlies both action slips and error-making in ADS (27, 146, 147). Schemas, as learned models of typical scenarios, are likely dependent on long-term storage mechanisms. Their chronic disruption is therefore likely to result from structural change, rather than the temporary, noise-based disruption modelled in the previous simulation. Therefore, to model schema disruption for this simulation we lesioned a randomly selected proportion of intrinsic PFC connections. We ran seven simulations of 200 trials each for both tea and coffee tasks, lesioning 1, 2, 5, 8, 10, 12 and 14% of intrinsic PFC connections, respectively.

#### Results

Like the previous simulation, this simulation resulted in a fairly gradual increase in the number of erroneous trials with increasing disruption for both tasks, as shown by the number of correct trials at each level of disruption illustrated in figure 19 (top).

**Figure 19:**
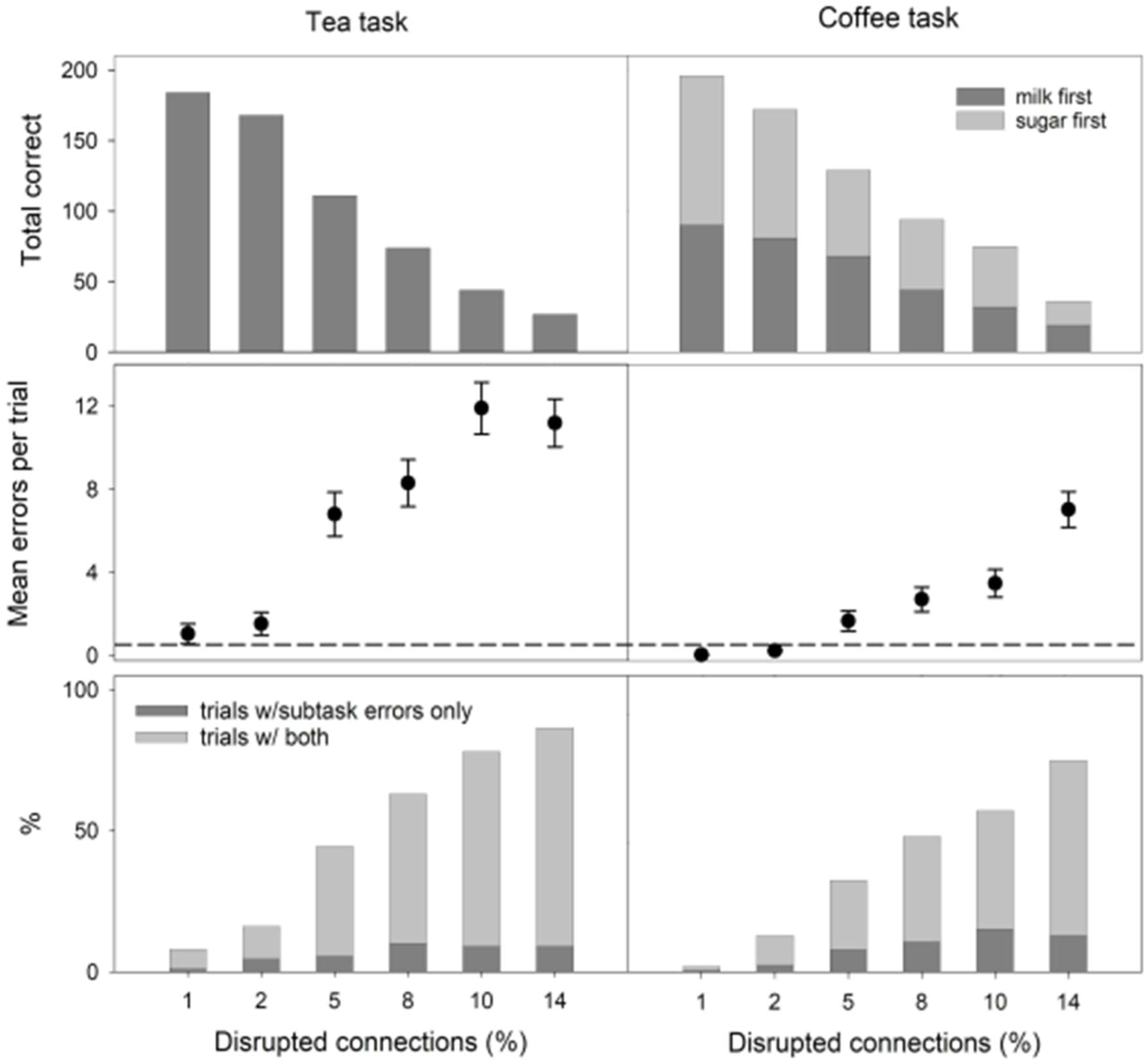
(Top) overall model performance shown as the percentage of correctly performed trials at each level of disruption for both tasks, out of 200 trials run at each noise level. (Middle) Mean errors per trial and sample SE at each level of noise. Dashed line shows threshold for classification as impaired performance. (Bottom) Percentage of trials displaying subtask based errors only, and subtask based and single action errors.

#### Normal + pathological error making

According to the hypothesis that action slips result from schema degradation, low levels of disruption in this simulation should result in ‘healthy’ performance, with few overall errors consisting mainly of subtask based errors (143, 144). Interestingly, at low disruption, although most trials were performed correctly, the average number of errors per trial was high, indicating a small number of highly error prone trials. The tea task averaged 1.04 errors per trial at the lowest disruption level (based on a sample of 100 trials), compared with zero in the previous simulation (figure 13). The present simulation thus produced ‘pathological’ error rates even at low disruption according to the 0.5 errors-per-trial threshold, particularly for the tea task (figure 19, middle).

Furthermore, the model predominantly produced single action errors; we saw few subtask-based errors at any disruption level (figure 19, bottom). While the coffee task was slightly more robust to errors in general, in this task again, most errors were single-action errors. This contrasts with the previous section dealing with disruption of temporal order knowledge in which, at low noise, trials containing only subtask-based errors comprised up to 58% of erroneous trials, better reflecting the observational literature. In the current simulation, the maximum proportion of erroneous trials comprising subtask-only errors was just 28% (figure 20). Thus, where the previous simulation accounted well for healthy error-making, these results suggest the disruption of action schemas does not account for normal slips of action.

**Figure 20:**
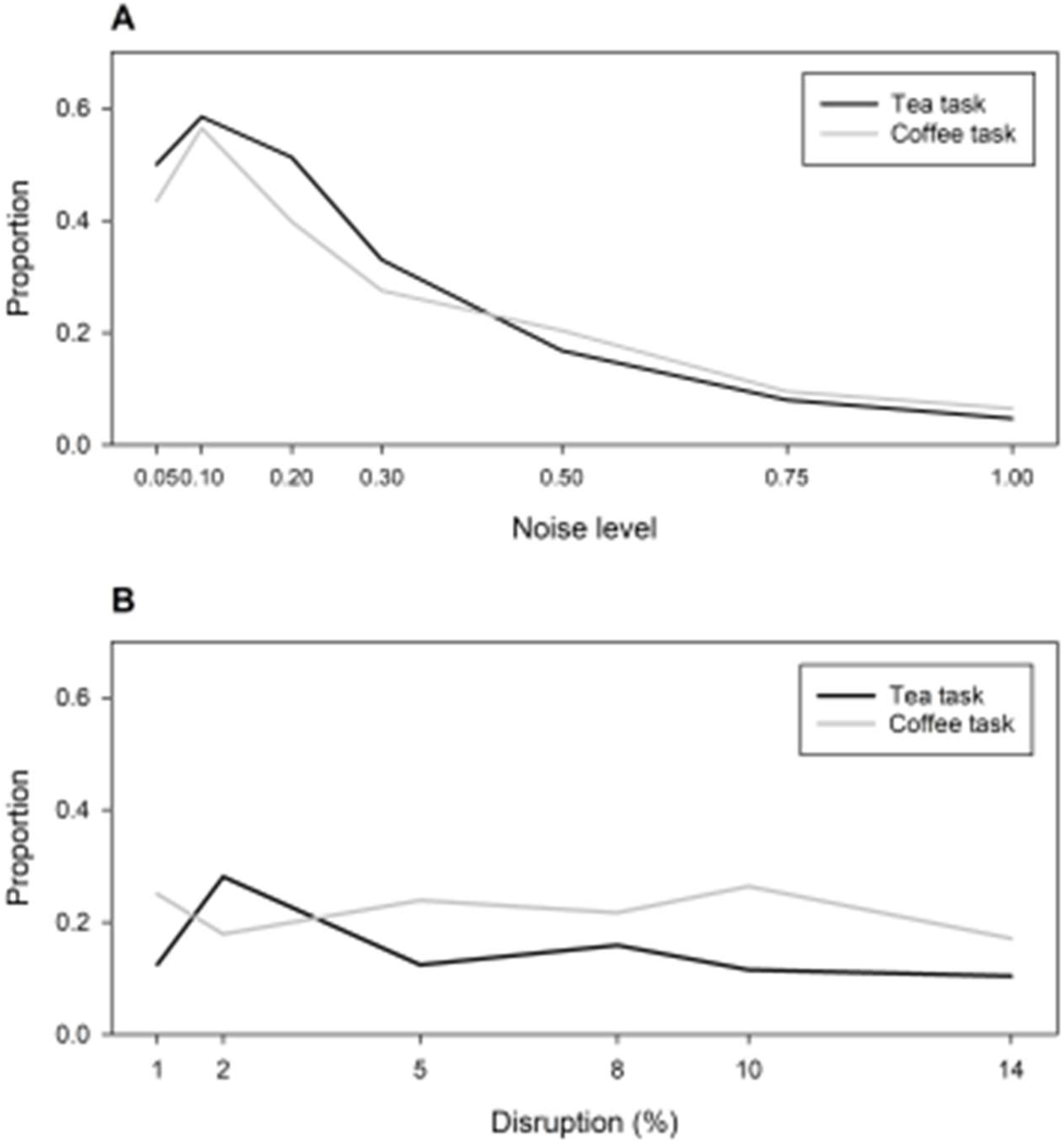
Proportion of erroneous trials displaying only subtask-based errors in simulations i (top) and ii (bottom). Disrupted order knowledge resulted in a higher proportion of subtask-only error trials at lower disruption, similar to patterns observed in human behavioural studies. In contrast, disrupted schemas resulted in a low proportion of subtask-only error trials across all levels of disruption.

#### Error types and rates

In the tea task, a tendency to perform many continuous perseverations of single actions accounted for most of the total errors committed and was chiefly responsible for the high numbers of errors at low levels of disruption. A sample of 100 trials at disruption levels of 2, 8 and 14% indicated that perseverations accounted for around 70-80% of all errors for the tea task. Omissions, in contrast, accounted for around 10%. Where omissions did occur, this was frequently a result of the model getting ‘stuck’ in continuous perseverations, becoming unable to complete a subtask. Perseverations and omissions together accounted for around 90% of all errors. Other sequence errors accounted for a very small proportion of all errors, averaging around 1-3%.

The coffee task did not exhibit as many perseverations, with these accounting for up to 58% of all errors, compared to 30% omissions. These rates were consistent across ‘impaired’ conditions, not varying significantly with increasing disruption. Sequence errors again accounted for a small proportion of all errors. The finding that there were significantly more perseverations than other error types across trials and tasks is not consistent with observational data on error patterns in ADS, suggesting that, like healthy error-making, schema disruption does not account for general pathological patterns of error making in ADS.

#### A perseverative ADS subtype?

While schema disruption does not appear to account for the general pathological patterns of error making in ADS, closer examination of the characterisation of these errors suggests it may account for a subclass therein. Forde and Humphreys (145) pointed to a dissociation between recurrent and continuous perseverations in two patients, FK and HG, and suggested distinct underlying deficits. This coupled with the dominance of continuous perseverations in our current simulation may point towards some disruption of schemas in patients who show a strong tendency to perseverate. This tendency might be understood in terms of attractor dynamics in the ‘state space’ of the recurrent neural network that comprises the model’s associative loop (62, 81). PFC representations can be considered stable attractors in the associative loop system’s state space. Transition nodes can be thought of as providing the ‘energy’ required to move the system from one representation – or attractor – to another. When connections in the associative loop are intact, we assume each attractor has a similar ‘depth’, so that the energy supplied by the transition layer is sufficient to move the system out of any given attractor. Suppose that, by lesioning PFC connectivity, the relative depth of each of these attractors is disrupted; some are made ‘deeper’, others ‘shallower’. The greater the proportion of connections lesioned, the more exaggerated this effect may be. After lesioning, once the system makes a transition into a ‘deep’ attractor, transition node activity may no longer be sufficient to drive changes to the expressed PFC representation, resulting in continued perseverations of the same action.

#### Summary

The simulation of schema disruption produced a strikingly different profile of errors to that observed for the simulation examining disrupted sequence knowledge. Schema disruption cannot account for the subtask-oriented errors typically performed by healthy controls under distraction, and is therefore unlikely to be the primary result of distraction in healthy controls. Nor is it likely that schema disruption is a central mechanism in ADS, as it does not account for the relative rates of errors typically observed in human observational studies. However, it is possible that schema disruption is a component of a wider pattern of damage in ADS. This notion is supported by the existence of patients such as HG, in whom a strong tendency to perseverate may point to the involvement of schema disruption; indeed, we would predict that ADS with such a tendency would show evidence of action schema breakdown on tests such as those utilised by Forde et al. (146).

### 4.5. Discussion

We have demonstrated that disruption of temporal order knowledge could preferentially account for a wide variety of trends observed human error data in both healthy controls and ADS patients, while disruption of action schemas themselves had less explanatory power. In particular, degradation of temporal order knowledge account for the dominance of subtask based errors in healthy controls, the dominance of omission errors in patients, and both omission and sequence errors in patient data. However, schema disruption may account for a particular nuance in the patient data, possibly accounting for patients with a propensity to commit continuous perseverations (145).

In error simulation i, noise applied to the transition layer ‘contextual confusion’, resulting from the overlapping nature of feature-based representations of context and environment. This organisation has the result that higher levels of noise increase the probability of activating increasingly dissimilar representations to the intended one. At low noise, the model tended to select representations corresponding to the correct *action*, but in an incorrect context. This manifested in behavioural errors at a later stage of the task, a phenomenon which has been observed previously in modelling (25) and experimental (149) work. At higher noise this resulted in selection of more dissimilar representations to the intended one, causing single action errors and disjointed behaviour. We also found that noise caused spurious and/or untimely activity in transition nodes, causing interference or premature switching of PFC representations, often resulting in anticipation-omissions, object substitutions and continuous perseverations.

In error simulation ii, damage to intrinsic PFC connectivity – and thus to the schemas such connectivity supported – appeared to critically destabilise a set of representations, whereas other representations became ‘super’ stable. At low disruption, this manifested differently in the two tasks. This is possibly due to a greater robustness of the coffee task, resulting from the flexibility of the milk and sugar subtasks. Having two versions of each representation for these subtasks in the coffee task may have permitted a greater degree of disruption before large numbers of errors occurred, effectively allowing a ‘backup’ version of the task to be performed if one was damaged. At higher disruption, the damage was sufficiently severe that performance resembled that for the tea task, where few representations remain ‘intact’ and the model is unable to perform a task to completion.

That different error profiles resulted from different types of disruption suggests that action schemas and knowledge of their temporal order may be represented distinctly in the brain. This is consistent with evidence that different components of PFC are selectively involved in the selection and maintenance of working memory representations (139, 140). Selective damage to such areas would be expected to result in the distinct error profiles we observed in our simulations.

#### Cognitive vs behavioural identification of errors

This study illustrated that different types of disruption at the cognitive level may have the same or similar behavioural manifestations, and that the behavioural manifestation of an error may be qualitatively different to the cognitive dynamics inducing it. Premature switching of PFC representations – an ‘omission’ at the cognitive level – frequently caused a perseveration error at the behavioural level. This suggests that omissions – particularly of single actions – may not be a problem of selection, but a problem of *maintenance*, with the further implication that ADS may not simply be a disorder of selection, but also one of timing. Furthermore, errors occurred which would not intuitively be attributed to the disruption that underpinned them. The disruption of temporal order knowledge was responsible for a number of errors, such as ‘toying’ behaviour and object substitutions, which have been said to be caused by imbalanced cognitive and sensory influences on behaviour (16, 28).

It is important for future behavioural studies to examine tasks which allow the distinction of ‘cognitive’ error types where possible. In the current tasks, for example, correct performance involved the repeated pouring of items into the cup. In a behavioural study, use of such tasks could lead to a situation where many cognitive level errors result in an apparent perseveration. Tasks involving more distinct component actions might provide more opportunity to understand the cognitive processes underlying error types.

#### ‘Suitability queueing’

Humphreys and colleagues (27, 145, 146) interpreted sequence knowledge in terms of a competitive queueing account, in which disruption was envisaged as a degradation of the activation gradient and rebound inhibition across component action schemas. This suggests all actions in a sequence are activated from the outset, and that each representation is automatically selected as the prior action is inhibited. Our model provides a closely related, but more interactive view of sequential selection in which pre-activation is not limited to the next temporal step, but to *multiple candidate next-actions,* similarly to the affordance competition hypothesis (121, 151). This pre-activation results from the overlapping projections from PFC to the transition layer, and is indicative not of an action’s rank order in the sequence as in competitive queueing accounts, but of its similarity with the intended subsequent action. This then becomes a ‘suitability-based’ queueing account, rather than the more traditional order-based account. Importantly, this allows for influences on the cognitive representations of actions to be dynamic; each representation receives pre-activation in an ‘on-the-fly’ manner according to the evolving temporal context, rather than a static gradient of activation from the task outset. Sequential performance may then be more flexible, taking into account changing information from the environment which may alter the planned sequence, rather than requiring a predetermined order. Since each representation is individually stable, no external gradient of activation need be maintained upon later actions in the sequence, and inhibition on prior actions need not be explicitly maintained to prevent reactivation. This avoids the inherent problems of maintaining inhibition on previously selected actions over the course of a sequence when the same action is required multiple times in a single task.

Findings showing pre-preparedness of action representations (22) have been cited as evidence for competitive queuing. However, these findings are also consistent with our model, and our suitability-queuing account. Furthermore, our suitability-queueing account better explains behavioural data: as Botvinick & Plaut (25) discuss and we have shown, subtask-based error dominance at low noise results from confusion between similar contexts due to similarity in their representations. A traditional competitive queueing account, however, would imply that low noise would result in single action errors where confusion occurs between two adjacent actions in a sequence, which the human behavioural data do not support.

#### Variability across tasks

The appearance of more errors in the coffee task in error simulation i, and for the tea task in error simulation ii, suggests that different tasks are robust to different types of disruption. Additionally, rates of particular error types differed for each task. This is consistent with previous research (145), and indicates that errors are a function of task features. Error profiles may reflect the specific cognitive demands and flexibility of the tasks. Longer tasks are probably more vulnerable to omissions: there are more actions to omit, and the probability of a mistake increases with sequence length. Greater flexibility in the order of component subtasks may result in more recurrent perseverations and subtask omissions, due to a greater chance of ‘losing track’. Subtasks without visible results (e.g., adding sugar) should experience more omissions and repetitions, and similar tasks are likely to suffer more intrusions from one another. Future experimental work should focus on relating task demands to the errors observed to better understand the underlying cognitive processes.

#### Comparison with existing models

Previous models simulating different types of disruption have accounted for a range of errors and error rates, and it is likely that ADS patients suffer from each of these problems to a greater or lesser degree. Botvinick & Plaut (25) applied noise to the hidden layer of the SRN model during testing, which is comparable with the ‘generalised cognitive deficit’ hypothesis (28). Notably, their overall rates of omissions were markedly higher than those observed in the behavioural data. Omissions may be caused by disruption to several different processes, suggesting that their high omission rate might be a result of damage to multiple embedded processes in the SRN hidden layer. Given the abstract emergent representations in their model however, it is not possible to examine the effects of damage to individual processes in their model. Conversely, the IAN model (16) accounts less well for the patterns seen in the human errors literature than the SRN model, but gives a more precise account of the disruption leading to the observed errors. This is because the IAN model incorporates a separation of function into distinct regions. This allows a deeper analysis of the nature of the errors that are observed with regard to specific processes. This is made particularly clear in a follow up study which suggested that disruption of separate processes differentially accounted for patterns of errors in distinct disorders of action (12).

Our model points to a reconciliation of the IAN and SRN models. The partially distributed nature of the representations of schemas we have included allows the production of many of the human-like error profiles produced by the SRN model (25), and an explanation of errors at the mechanistic level in terms of the underlying representations. This is despite the hierarchical structure of the representations themselves. By separating function into distinct regions of the model, we also retain the explanatory power of the IAN model (16). Additionally, given the neuroanatomical basis of our model, we can interpret our results in terms of the underlying neural substrate. The real power of our model then lies not just in the amount of empirical data that we have replicated, but in its ability to provide insight into the underlying mechanistic and neural processes governing sequential performance; a combination which has not previously been achieved. This ‘hybrid’ nature of our model allows us to confront several problems discussed by each set of authors. Cooper & Shallice (32) suggest that errors in the SRN model occur only when the error exists as a correct action elsewhere in the training corpus. We have shown, however, that in a model utilising a recurrent network, omissions of both single steps and entire subtasks occur even when the preceding and following actions do not appear together in any valid version of the task. Conversely, Botvinick & Plaut (68) point out that patterns in the behavioural literature were not replicated by the IAN model. However, by implementing schemas as overlapping representations and disrupting a process which may be central to ADS, we have shown that this lack of replication is not due to the fundamental organisation of the model in terms of schemas and goals within a hierarchical framework. Rather, this is more likely to be due to the locus of disruption in the original study.

The macro-architecture of our model is similar to some existing models which have also utilised cross-loop projections in a similar fashion (78, 95, 107). Such models have successfully been applied to various learning and memory tasks, highlighting the diverse functionality afforded by this architecture. Our main contribution here, then, is the novel organisation of the associative loop, which, in combination with the cross-loop projections, has led directly to our main findings.

## 5. Conclusion

By bringing together insights from cognitive psychology, neurobiology and computational modelling to develop a novel, neuroanatomically grounded model of routine sequential behaviour, we have provided a reconciliation of two prominent models of cognition and action (16, 25), and an account of behavioural data from healthy volunteers (143, 144) and disorders of action (5, 27), which remains consistent with neurophysiological data from sequencing tasks (22). This unification relies on an innovative functional architecture in associative areas embodying our subset-selection hypothesis, which allows competitive selection, maintenance and deselection of distributed representations by BG, flexible goal-directed action execution, and hierarchical functional organisation of task representations. Further, our model provides evidence in favour of an existing but previously untested hypothesis of the mechanisms underlying sequence breakdown in ADS patients (27), and offers mechanistic explanations that are grounded in neuroanatomy, thereby providing a platform for testable predictions linking brain function and task performance (discussed in sections 3.6 & 4.4). We have also suggested a new interpretation of the competitive queuing hypothesis – termed the ‘suitability queueing account’, which better accounts for behavioural data pertaining to actions of error.

On the basis of neuroanatomical (125, 130–133), neurophysiological (47, 110) and neuropsychological (5, 27) evidence, and the well-supported hypothesis that BG are specialised for selection (34, 35), we suggested the hierarchical BGTC system is a likely substrate for the organisation of goal-directed action sequences. Our model of this system, developed from the ‘GPR’ model of action selection in BG (39–41), introduced a novel, ‘subset-based’ functional architecture of the selection of contextual information in associative BGTC loops. Our associative loop model represents the first neuroanatomically constrained model of associative BG which implements competitive selection, timely deselection, active maintenance of distributed representations in PFC, and convergence in corticostriatal projections; features which have previously been considered incompatible (101). We have shown how selection of ‘schemas’ for tasks and subtasks may be used to modulate selection of action affordances to produce coherent sequential behaviour.

Context sensitive action selection in the model is driven by integration of sensory and contextual influences in motor BGTC territories (117, 118, 131). Sequence generation relies on a novel transitioning mechanism consistent with evidence that stable representations require external influence to generate transitions (137); this may represent a component of PFC distinct from that responsible for maintenance (139, 140). The complete model performed sequential tasks requiring action repetition within and between tasks and ‘memory’ of task history, and the ‘quasi-hierarchical’ waiter scenario, which, according to Botvinick & Plaut (25), should be difficult for this model given its semi-localist representation. Analyses showed similar dynamics to both the SRN (25) and the IAN (16) models, as well as resembling existing neural data (22), thus pointing towards a reconciliation of these models and a neurally focused account of their processing. Testing the model under disruption to examine its ability to account for action errors lent support to the hypothesis that many documented error profiles can result from damaged temporal order knowledge. More nuanced patterns of disruption may result in more specific deficits; in particular, the tendency to commit continuous perseverations (145) may result from changes in the stability of schemas.

### Potential model developments

Beyond the achievements of the present research, there are several areas in which the current model can be expanded to address further questions. Including learning processes may allow examination of the mechanisms required to develop connectivity necessary to mediate routine sequencing. More complex task environments, spatial parameters, semantic object knowledge and the inclusion of release and orienting actions would increase the opportunity for particular error types. This could serve to further validate our conclusions regarding the role of temporal order knowledge in ADS. Inclusion of error monitoring would facilitate the exploration of the role of error detection and remedy in overall performance. In extensions of earlier work Cooper and colleagues found that disruption of distinct processes accounted preferentially for error data from patients suffering from ADS, ideational apraxia and utilisation syndrome (29), and more recently, Parkinson’s disease (33). Future developments to the current model should aim to account for such data. Indeed, the GPR model upon which the present study is based has been shown to display problems with selection, reminiscent of Parkinsonian akinesia, as a result of reduced levels of simulated tonic dopamine in striatum (40), lending the present model well to modelling the effects of Parkinson’s disease on routine action in a multi-loop architecture.

## Data availability

Data and the code used to generate it are available upon request to the corresponding author.

