## Supplementary Material A for "RoSe-BaL: A neuroanatomically plausible model of routine action sequencing"

### Full list of parameters: Associative loop model

**A1 Table: Neuron output thresholds/gradients**

| Nucleus | Threshold ϵ | Output gradient *m* |
|---|---|---|
| PFC | 0 | 1 |
| Thalamus | 0 | 1 |
| Caudate | 0.2 | 1 |
| STN | -0.25 | 1 |
| GPe | -0.2 | 1 |
| GPi | -0.2 | 1 |

**A2 Table: Simulation parameters**

| Parameter | Value | Units |
|---|---|---|
| T | 3 | Seconds |
| $\tau_m$ | 0.04 | Seconds |
| dt | 0.001 | Seconds |
| θ | 0.9 | *NA* |

**A3 Table: Associative loop scalar synaptic weight values and type**

| Source | Target | Type | Weight |
|---|---|---|---|
| PFC | Caudate | Subset → channel | 0.25 |
| PFC | Thalamus | Subset → channel | 0.25 |
| PFC | STN | Subset → channel | 0.4 |
| Caudate (D1) | GPi | Channel-wise | 1 |
| Caudate (D2) | GPe | Channel-wise | 1 |
| STN | GPe | Diffuse | 0.8 |
| STN | GPi | Diffuse | 0.8 |
| GPe | STN | Channel-wise | 1 |
| GPe | GPi | Channel-wise | 0.3 |
| GPi | Thalamus | Channel-wise | 1 |
| Dopaminergic modulation | Caudate | *NA* | 0.2 |

**A4 Table: Thalamus → PFC weight matrix**
Rows = Source thalamus channels
Columns = Target PFC nodes

| | 1 | 2 | 3 | 4 | 5 | 6 | 7 | 8 | 9 | 10 | 11 | 12 | 13 | 14 | 15 | 16 | 17 | 18 |
|---|---|---|---|---|---|---|---|---|---|---|---|---|---|---|---|---|---|---|
| 1 | 0.8 | 0.8 | 0 | 0 | 0 | 0 | 0.8 | 0.8 | 0 | 0 | 0 | 0 | 0 | 0 | 0 | 0 | 0 | 0 |
| 2 | 0.8 | 0.8 | 0 | 0 | 0 | 0 | 0.8 | 0.8 | 0 | 0 | 0 | 0 | 0 | 0 | 0 | 0 | 0 | 0 |
| 3 | 0.8 | 0.8 | 0 | 0 | 0 | 0 | 0.8 | 0.8 | 0 | 0 | 0 | 0 | 0 | 0 | 0 | 0 | 0 | 0 |

| | 19 | 20 | 21 | 22 | 23 | 24 | 25 | 26 | 27 | 28 | 29 | 30 | 31 | 32 | 33 | 34 | 35 | 36 |
|---|---|---|---|---|---|---|---|---|---|---|---|---|---|---|---|---|---|---|
| 1 | 0 | 0 | 0 | 0 | 0 | 0 | 0 | 0 | 0 | 0 | 0.8 | 0.8 | 0.8 | 0.8 | 0 | 0 | 0 | 0 |
| 2 | 0 | 0 | 0 | 0 | 0 | 0 | 0.8 | 0.8 | 0 | 0 | 0 | 0 | 0 | 0 | 0.8 | 0.8 | 0 | 0 |
| 3 | 0 | 0 | 0 | 0 | 0 | 0 | 0 | 0 | 0.8 | 0.8 | 0 | 0 | 0 | 0 | 0 | 0 | 0.8 | 0.8 |

**A5 Table: Recurrent PFC weight matrix**

Rows = Source PFC nodes; Columns = Target PFC nodes

Continued on next page.

| | 1 | 2 | 3 | 4 | 5 | 6 | 7 | 8 | 9 | 10 | 11 | 12 | 13 | 14 | 15 | 16 | 17 | 18 | 19 | 20 | 21 |
|---|---|---|---|---|---|---|---|---|---|---|---|---|---|---|---|---|---|---|---|---|---|
| 1 | 0 | -0.3 | 0 | 0 | 0 | 0 | 0.15 | -0.3 | 0 | 0 | 0 | 0 | 0 | 0 | 0 | 0 | 0 | 0 | 0 | 0 | 0 |
| 2 | -0.3 | 0 | 0 | 0 | 0 | 0 | -0.3 | 0.15 | 0 | 0 | 0 | 0 | 0 | 0 | 0 | 0 | 0 | 0 | 0 | 0 | 0 |
| 3 | 0 | 0 | 0 | 0 | 0 | 0 | 0 | 0 | 0 | 0 | 0 | 0 | 0 | 0 | 0 | 0 | 0 | 0 | 0 | 0 | 0 |
| 4 | 0 | 0 | 0 | 0 | 0 | 0 | 0 | 0 | 0 | 0 | 0 | 0 | 0 | 0 | 0 | 0 | 0 | 0 | 0 | 0 | 0 |
| 5 | 0 | 0 | 0 | 0 | 0 | 0 | 0 | 0 | 0 | 0 | 0 | 0 | 0 | 0 | 0 | 0 | 0 | 0 | 0 | 0 | 0 |
| 6 | 0 | 0 | 0 | 0 | 0 | 0 | 0 | 0 | 0 | 0 | 0 | 0 | 0 | 0 | 0 | 0 | 0 | 0 | 0 | 0 | 0 |
| 7 | 0.15 | -0.3 | 0 | 0 | 0 | 0 | 0 | -0.3 | 0 | 0 | 0 | 0 | 0 | 0 | 0 | 0 | 0 | 0 | 0 | 0 | 0 |
| 8 | -0.3 | 0.15 | 0 | 0 | 0 | 0 | -0.3 | 0 | 0 | 0 | 0 | 0 | 0 | 0 | 0 | 0 | 0 | 0 | 0 | 0 | 0 |
| 9 | 0 | 0 | 0 | 0 | 0 | 0 | 0 | 0 | 0 | 0 | 0 | 0 | 0 | 0 | 0 | 0 | 0 | 0 | 0 | 0 | 0 |
| 10 | 0 | 0 | 0 | 0 | 0 | 0 | 0 | 0 | 0 | 0 | 0 | 0 | 0 | 0 | 0 | 0 | 0 | 0 | 0 | 0 | 0 |
| 11 | 0 | 0 | 0 | 0 | 0 | 0 | 0 | 0 | 0 | 0 | 0 | 0 | 0 | 0 | 0 | 0 | 0 | 0 | 0 | 0 | 0 |
| 12 | 0 | 0 | 0 | 0 | 0 | 0 | 0 | 0 | 0 | 0 | 0 | 0 | 0 | 0 | 0 | 0 | 0 | 0 | 0 | 0 | 0 |
| 13 | 0 | 0 | 0 | 0 | 0 | 0 | 0 | 0 | 0 | 0 | 0 | 0 | 0 | 0 | 0 | 0 | 0 | 0 | 0 | 0 | 0 |
| 14 | 0 | 0 | 0 | 0 | 0 | 0 | 0 | 0 | 0 | 0 | 0 | 0 | 0 | 0 | 0 | 0 | 0 | 0 | 0 | 0 | 0 |
| 15 | 0 | 0 | 0 | 0 | 0 | 0 | 0 | 0 | 0 | 0 | 0 | 0 | 0 | 0 | 0 | 0 | 0 | 0 | 0 | 0 | 0 |
| 16 | 0 | 0 | 0 | 0 | 0 | 0 | 0 | 0 | 0 | 0 | 0 | 0 | 0 | 0 | 0 | 0 | 0 | 0 | 0 | 0 | 0 |
| 17 | 0 | 0 | 0 | 0 | 0 | 0 | 0 | 0 | 0 | 0 | 0 | 0 | 0 | 0 | 0 | 0 | 0 | 0 | 0 | 0 | 0 |
| 18 | 0 | 0 | 0 | 0 | 0 | 0 | 0 | 0 | 0 | 0 | 0 | 0 | 0 | 0 | 0 | 0 | 0 | 0 | 0 | 0 | 0 |
| 19 | 0 | 0 | 0 | 0 | 0 | 0 | 0 | 0 | 0 | 0 | 0 | 0 | 0 | 0 | 0 | 0 | 0 | 0 | 0 | 0 | 0 |
| 20 | 0 | 0 | 0 | 0 | 0 | 0 | 0 | 0 | 0 | 0 | 0 | 0 | 0 | 0 | 0 | 0 | 0 | 0 | 0 | 0 | 0 |
| 21 | 0 | 0 | 0 | 0 | 0 | 0 | 0 | 0 | 0 | 0 | 0 | 0 | 0 | 0 | 0 | 0 | 0 | 0 | 0 | 0 | 0 |
| 22 | 0 | 0 | 0 | 0 | 0 | 0 | 0 | 0 | 0 | 0 | 0 | 0 | 0 | 0 | 0 | 0 | 0 | 0 | 0 | 0 | 0 |
| 23 | 0 | 0 | 0 | 0 | 0 | 0 | 0 | 0 | 0 | 0 | 0 | 0 | 0 | 0 | 0 | 0 | 0 | 0 | 0 | 0 | 0 |
| 24 | 0 | 0 | 0 | 0 | 0 | 0 | 0 | 0 | 0 | 0 | 0 | 0 | 0 | 0 | 0 | 0 | 0 | 0 | 0 | 0 | 0 |
| 25 | -0.25 | 0 | 0 | 0 | 0 | 0 | -0.25 | 0 | 0 | 0 | 0 | 0 | 0 | 0 | 0 | 0 | 0 | 0 | 0 | 0 | 0 |
| 26 | -0.25 | 0 | 0 | 0 | 0 | 0 | -0.25 | 0 | 0 | 0 | 0 | 0 | 0 | 0 | 0 | 0 | 0 | 0 | 0 | 0 | 0 |
| 27 | -0.25 | 0 | 0 | 0 | 0 | 0 | -0.25 | 0 | 0 | 0 | 0 | 0 | 0 | 0 | 0 | 0 | 0 | 0 | 0 | 0 | 0 |
| 28 | -0.25 | 0 | 0 | 0 | 0 | 0 | -0.25 | 0 | 0 | 0 | 0 | 0 | 0 | 0 | 0 | 0 | 0 | 0 | 0 | 0 | 0 |
| 29 | -0.25 | 0 | 0 | 0 | 0 | 0 | -0.25 | 0 | 0 | 0 | 0 | 0 | 0 | 0 | 0 | 0 | 0 | 0 | 0 | 0 | 0 |
| 30 | -0.25 | 0 | 0 | 0 | 0 | 0 | -0.25 | 0 | 0 | 0 | 0 | 0 | 0 | 0 | 0 | 0 | 0 | 0 | 0 | 0 | 0 |
| 31 | 0 | -0.25 | 0 | 0 | 0 | 0 | 0 | -0.25 | 0 | 0 | 0 | 0 | 0 | 0 | 0 | 0 | 0 | 0 | 0 | 0 | 0 |
| 32 | 0 | -0.25 | 0 | 0 | 0 | 0 | 0 | -0.25 | 0 | 0 | 0 | 0 | 0 | 0 | 0 | 0 | 0 | 0 | 0 | 0 | 0 |
| 33 | 0 | -0.25 | 0 | 0 | 0 | 0 | 0 | -0.25 | 0 | 0 | 0 | 0 | 0 | 0 | 0 | 0 | 0 | 0 | 0 | 0 | 0 |
| 34 | 0 | -0.25 | 0 | 0 | 0 | 0 | 0 | -0.25 | 0 | 0 | 0 | 0 | 0 | 0 | 0 | 0 | 0 | 0 | 0 | 0 | 0 |
| 35 | 0 | -0.25 | 0 | 0 | 0 | 0 | 0 | -0.25 | 0 | 0 | 0 | 0 | 0 | 0 | 0 | 0 | 0 | 0 | 0 | 0 | 0 |
| 36 | 0 | -0.25 | 0 | 0 | 0 | 0 | 0 | -0.25 | 0 | 0 | 0 | 0 | 0 | 0 | 0 | 0 | 0 | 0 | 0 | 0 | 0 |

**A5 Table: Recurrent PFC weight matrix**

Continued from previous page.

| | 22 | 23 | 24 | 25 | 26 | 27 | 28 | 29 | 30 | 31 | 32 | 33 | 34 | 35 | 36 |
|---|---|---|---|---|---|---|---|---|---|---|---|---|---|---|---|
| 1 | 0 | 0 | 0 | -0.25 | -0.25 | -0.25 | -0.25 | -0.25 | -0.25 | 0.15 | 0.15 | 0.15 | 0.15 | 0.15 | 0.15 |
| 2 | 0 | 0 | 0 | 0.15 | 0.15 | 0.15 | 0.15 | 0.15 | 0.15 | -0.25 | -0.25 | -0.25 | -0.25 | -0.25 | -0.25 |
| 3 | 0 | 0 | 0 | 0 | 0 | 0 | 0 | 0 | 0 | 0 | 0 | 0 | 0 | 0 | 0 |
| 4 | 0 | 0 | 0 | 0 | 0 | 0 | 0 | 0 | 0 | 0 | 0 | 0 | 0 | 0 | 0 |
| 5 | 0 | 0 | 0 | 0 | 0 | 0 | 0 | 0 | 0 | 0 | 0 | 0 | 0 | 0 | 0 |
| 6 | 0 | 0 | 0 | 0 | 0 | 0 | 0 | 0 | 0 | 0 | 0 | 0 | 0 | 0 | 0 |
| 7 | 0 | 0 | 0 | -0.25 | -0.25 | -0.25 | -0.25 | -0.25 | -0.25 | 0.15 | 0.15 | 0.15 | 0.15 | 0.15 | 0.15 |
| 8 | 0 | 0 | 0 | 0.15 | 0.15 | 0.15 | 0.15 | 0.15 | 0.15 | -0.25 | -0.25 | -0.25 | -0.25 | -0.25 | -0.25 |
| 9 | 0 | 0 | 0 | 0 | 0 | 0 | 0 | 0 | 0 | 0 | 0 | 0 | 0 | 0 | 0 |
| 10 | 0 | 0 | 0 | 0 | 0 | 0 | 0 | 0 | 0 | 0 | 0 | 0 | 0 | 0 | 0 |
| 11 | 0 | 0 | 0 | 0 | 0 | 0 | 0 | 0 | 0 | 0 | 0 | 0 | 0 | 0 | 0 |
| 12 | 0 | 0 | 0 | 0 | 0 | 0 | 0 | 0 | 0 | 0 | 0 | 0 | 0 | 0 | 0 |
| 13 | 0 | 0 | 0 | 0 | 0 | 0 | 0 | 0 | 0 | 0 | 0 | 0 | 0 | 0 | 0 |
| 14 | 0 | 0 | 0 | 0 | 0 | 0 | 0 | 0 | 0 | 0 | 0 | 0 | 0 | 0 | 0 |
| 15 | 0 | 0 | 0 | 0 | 0 | 0 | 0 | 0 | 0 | 0 | 0 | 0 | 0 | 0 | 0 |
| 16 | 0 | 0 | 0 | 0 | 0 | 0 | 0 | 0 | 0 | 0 | 0 | 0 | 0 | 0 | 0 |
| 17 | 0 | 0 | 0 | 0 | 0 | 0 | 0 | 0 | 0 | 0 | 0 | 0 | 0 | 0 | 0 |
| 18 | 0 | 0 | 0 | 0 | 0 | 0 | 0 | 0 | 0 | 0 | 0 | 0 | 0 | 0 | 0 |
| 19 | 0 | 0 | 0 | 0 | 0 | 0 | 0 | 0 | 0 | 0 | 0 | 0 | 0 | 0 | 0 |
| 20 | 0 | 0 | 0 | 0 | 0 | 0 | 0 | 0 | 0 | 0 | 0 | 0 | 0 | 0 | 0 |
| 21 | 0 | 0 | 0 | 0 | 0 | 0 | 0 | 0 | 0 | 0 | 0 | 0 | 0 | 0 | 0 |
| 22 | 0 | 0 | 0 | 0 | 0 | 0 | 0 | 0 | 0 | 0 | 0 | 0 | 0 | 0 | 0 |
| 23 | 0 | 0 | 0 | 0 | 0 | 0 | 0 | 0 | 0 | 0 | 0 | 0 | 0 | 0 | 0 |
| 24 | 0 | 0 | 0 | 0 | 0 | 0 | 0 | 0 | 0 | 0 | 0 | 0 | 0 | 0 | 0 |
| 25 | 0 | 0 | 0 | 0 | 0.15 | -0.25 | -0.25 | -0.25 | -0.25 | 0 | 0 | -0.25 | -0.25 | 0 | 0 |
| 26 | 0 | 0 | 0 | 0.15 | 0 | -0.25 | -0.25 | -0.25 | -0.25 | 0 | 0 | -0.25 | -0.25 | 0 | 0 |
| 27 | 0 | 0 | 0 | -0.25 | -0.25 | 0 | 0.15 | -0.25 | -0.25 | 0 | 0 | 0 | 0 | -0.25 | -0.25 |
| 28 | 0 | 0 | 0 | -0.25 | -0.25 | 0.15 | 0 | -0.25 | -0.25 | 0 | 0 | 0 | 0 | -0.25 | -0.25 |
| 29 | 0 | 0 | 0 | -0.25 | -0.25 | -0.25 | -0.25 | 0 | 0.15 | -0.25 | -0.25 | 0 | 0 | 0 | 0 |
| 30 | 0 | 0 | 0 | -0.25 | -0.25 | -0.25 | -0.25 | 0.15 | 0 | -0.25 | -0.25 | 0 | 0 | 0 | 0 |
| 31 | 0 | 0 | 0 | 0 | 0 | 0 | 0 | -0.25 | -0.25 | 0 | 0.15 | -0.25 | -0.25 | -0.25 | -0.25 |
| 32 | 0 | 0 | 0 | 0 | 0 | 0 | 0 | -0.25 | -0.25 | 0.15 | 0 | -0.25 | -0.25 | -0.25 | -0.25 |
| 33 | 0 | 0 | 0 | -0.25 | -0.25 | 0 | 0 | 0 | 0 | -0.25 | -0.25 | 0 | 0.15 | -0.25 | -0.25 |
| 34 | 0 | 0 | 0 | -0.25 | -0.25 | 0 | 0 | 0 | 0 | -0.25 | -0.25 | 0.15 | 0 | -0.25 | -0.25 |
| 35 | 0 | 0 | 0 | 0 | 0 | -0.25 | -0.25 | 0 | 0 | -0.25 | -0.25 | -0.25 | -0.25 | 0 | 0.15 |
| 36 | 0 | 0 | 0 | 0 | 0 | -0.25 | -0.25 | 0 | 0 | -0.25 | -0.25 | -0.25 | -0.25 | 0.15 | 0 |
