## Supplementary Material B for "RoSe-BaL: A neuroanatomically plausible model of routine action sequencing"

### Full list of parameters: Complete RoSe-BaL model

#### B1: General model parameters

##### B1.1 Table: Neuron output thresholds/gradients

| Nucleus | Threshold ϵ | Output gradient *m* |
|---|---|---|
| PFC | 0 | 1 |
| Associative thalamus | 0 | 1 |
| Caudate | 0.2 | 1 |
| Associative STN | -0.25 | 1 |
| Associative GPe | -0.2 | 1 |
| Associative GPi | -0.2 | 1 |
| Pre/motor cortex | 0 | 1 |
| Motor thalamus | 0 | 1 |
| Putamen | 0.2 | 1 |
| Motor STN | -0.25 | 1 |
| Motor GPe | -0.2 | 1 |
| Motor GPi | -0.2 | 1 |
| Object representations | 0 | 1 |
| Action affordances | 0 | 1 |
| Transition nodes | 0.5 | 10 |

##### B1.2 Table Simulation parameters

| Parameter | Value | Units |
|---|---|---|
| T | ≤ 30 | Seconds |
| $\tau_m$ | 0.04 | Seconds |
| dt | 0.001 | Seconds |
| θ | 0.9 | *NA* |

#### B2: Model weights: basic sequencing

This section details the model weights for the basic sequencing simulation in section 3.5.

##### B2.1 Table: Associative loop scalar synaptic weight values and type

| Source | Target | Type | Weight |
|---|---|---|---|
| PFC | Caudate | Subset → channel | 0.3 |
| PFC | Associative Thalamus | Subset → channel | 0.15 |
| PFC | Associative STN | Subset → channel | 0.4 |
| Caudate (D1) | Associative GPi | Channel-wise | 1 |
| Caudate (D2) | Associative GPe | Channel-wise | 1 |
| Associative STN | Associative GPe | Diffuse | 0.8 |
| Associative STN | Associative GPi | Diffuse | 0.8 |
| Associative GPe | Associative STN | Channel-wise | 1 |
| Associative GPe | Associative GPi | Channel-wise | 0.3 |
| Associative GPi | Thalamus | Channel-wise | 1 |
| Dopaminergic modulation | Caudate | *NA* | 0.2 |

##### B2.2 Table Motor loop scalar synaptic weight values and type

| Source | Target | Type | Weight |
|---|---|---|---|
| PFC | Putamen | Action rep → channel | 0.3 |
| PFC | Motor STN | Diffuse | 0.01 |
| Action affordances | Pre/motor cortex | Channel-wise | 0.3 |
| Action affordances | Putamen | Channel-wise | 0.3 |
| Motor thalamus | Pre/motor cortex | Channel-wise | 1 |
| Pre/motor cortex | Putamen | Channel-wise | 0.2 |
| Pre/motor cortex | Motor Thalamus | Channel-wise | 1 |
| Pre/motor cortex | Motor STN | Channel-wise | 1 |
| Putamen (D1) | Motor GPi | Channel-wise | 1 |
| Putamen (D2) | Motor GPe | Channel-wise | 1 |
| Motor STN | Motor GPe | Diffuse | 0.8 |
| Motor STN | Motor GPi | Diffuse | 0.8 |
| Motor GPe | Motor STN | Channel-wise | 1 |
| Motor GPe | Motor GPi | Channel-wise | 0.3 |
| Motor GPi | Motor Thalamus | Channel-wise | 1 |
| Dopaminergic modulation | Putamen | *NA* | 0.2 |

##### B2.3 Table: Additional regions scalar synaptic weight values and type

| Source | Target | Type | Weight |
|---|---|---|---|
| Goal | Transition nodes | Scalar → first task node | 1 |
| Transition nodes | Transition nodes | All to all | 0.5 |
| Object representations | Object representations | All to all | 0.5 |
| Object representations | Action affordances | All to all | 0.3 |

##### B2.4 Table: Thalamus → PFC weight matrix

Rows = Source thalamus channels; Columns = Target PFC nodes

Continued below

| | 1 | 2 | 3 | 4 | 5 | 6 | 7 | 8 | 9 | 10 | 11 | 12 | 13 | 14 | 15 | 16 |
|---|---|---|---|---|---|---|---|---|---|---|---|---|---|---|---|---|
| 1 | 0.6 | 0.6 | 0 | 0 | 0.6 | 0.6 | 0 | 0 | 0 | 0 | 0 | 0 | 0 | 0 | 0.6 | 0.6 |
| 2 | 0 | 0 | 0.6 | 0.6 | 0 | 0 | 0.6 | 0.6 | 0 | 0 | 0 | 0 | 0 | 0 | 0 | 0 |
| 3 | 0.6 | 0.6 | 0.6 | 0.6 | 0 | 0 | 0 | 0 | 0.6 | 0.6 | 0 | 0 | 0 | 0 | 0 | 0 |
| 4 | 0 | 0 | 0.6 | 0.6 | 0 | 0 | 0 | 0 | 0 | 0 | 0.6 | 0.6 | 0 | 0 | 0 | 0 |
| 5 | 0.6 | 0.6 | 0.6 | 0.6 | 0 | 0 | 0 | 0 | 0 | 0 | 0 | 0 | 0.6 | 0.6 | 0 | 0 |

##### B2.4 Table: Thalamus → PFC weight matrix

| | 17 | 18 | 19 | 20 | 21 | 22 | 23 | 24 | 25 | 26 | 27 | 28 | 29 | 30 | 31 | 32 |
|---|---|---|---|---|---|---|---|---|---|---|---|---|---|---|---|---|
| 1 | 0 | 0 | 0 | 0 | 0 | 0 | 0 | 0 | 0 | 0 | 0.6 | 0.6 | 0.6 | 0.6 | 0.6 | 0.6 |
| 2 | 0.6 | 0.6 | 0.6 | 0.6 | 0 | 0 | 0 | 0 | 0 | 0 | 0.6 | 0.6 | 0.6 | 0.6 | 0 | 0 |
| 3 | 0 | 0 | 0 | 0 | 0.6 | 0.6 | 0 | 0 | 0 | 0 | 0.6 | 0.6 | 0.6 | 0.6 | 0 | 0 |
| 4 | 0 | 0 | 0 | 0 | 0 | 0 | 0.6 | 0.6 | 0 | 0 | 0.6 | 0.6 | 0.6 | 0.6 | 0 | 0 |
| 5 | 0.6 | 0.6 | 0 | 0 | 0 | 0 | 0 | 0 | 0.6 | 0.6 | 0.6 | 0.6 | 0.6 | 0.6 | 0 | 0 |

##### B2.4 Table: Thalamus → PFC weight matrix

| | 33 | 34 | 35 | 36 | 37 | 38 | 39 | 40 | 41 | 42 | 43 | 44 | 45 | 46 | 47 | 48 | 49 |
|---|---|---|---|---|---|---|---|---|---|---|---|---|---|---|---|---|---|
| 1 | 0 | 0 | 0 | 0 | 0 | 0 | 0 | 0 | 0 | 0 | 0 | 0 | 0 | 0 | 0 | 0 | 0 |
| 2 | 0.6 | 0.6 | 0.6 | 0.6 | 0 | 0 | 0 | 0 | 0 | 0 | 0 | 0 | 0 | 0 | 0 | 0 | 0 |
| 3 | 0.6 | 0.6 | 0 | 0 | 0 | 0 | 0 | 0 | 0 | 0 | 0 | 0 | 0 | 0 | 0 | 0 | 0 |
| 4 | 0.6 | 0.6 | 0 | 0 | 0.6 | 0.6 | 0 | 0 | 0.6 | 0.6 | 0 | 0 | 0 | 0 | 0 | 0 | 0 |
| 5 | 0.6 | 0.6 | 0.6 | 0.6 | 0 | 0 | 0.6 | 0.6 | 0 | 0 | 0.6 | 0.6 | 0 | 0 | 0 | 0 | 0 |

**B2.5 Table PFC → Object representations weight matrix**

Rows = Source PFC channels; Columns = Target object nodes

| | 1 | 2 | 3 | 4 | 5 | 6 | 7 |
|---|---|---|---|---|---|---|---|
| 1 | 0 | 0 | 0 | 0 | 0 | 0 | 0 |
| 2 | 0 | 0 | 0 | 0 | 0 | 0 | 0 |
| 3 | 0 | 0 | 0 | 0 | 0 | 0 | 0 |
| 4 | 0 | 0 | 0 | 0 | 0 | 0 | 0 |
| 5 | 0 | 0 | 0 | 0 | 0 | 0 | 0 |
| 6 | 0 | 0 | 0 | 0 | 0 | 0 | 0 |
| 7 | 0 | 0 | 0 | 0 | 0 | 0 | 0 |
| 8 | 0 | 0 | 0 | 0 | 0 | 0 | 0 |
| 9 | 0 | 0 | 0 | 0 | 0 | 0 | 0 |
| 10 | 0 | 0 | 0 | 0 | 0 | 0 | 0 |
| 11 | 0 | 0 | 0 | 0 | 0 | 0 | 0 |
| 12 | 0 | 0 | 0 | 0 | 0 | 0 | 0 |
| 13 | 0 | 0 | 0 | 0 | 0 | 0 | 0 |
| 14 | 0 | 0 | 0 | 0 | 0 | 0 | 0 |
| 15 | 0.3 | 0 | 0 | 0 | 0 | 0 | 0 |
| 16 | 0.3 | 0 | 0 | 0 | 0 | 0 | 0 |
| 17 | 0 | 0.3 | 0 | 0 | 0 | 0 | 0 |
| 18 | 0 | 0.3 | 0 | 0 | 0 | 0 | 0 |
| 19 | 0 | 0 | 0.3 | 0 | 0 | 0 | 0 |
| 20 | 0 | 0 | 0.3 | 0 | 0 | 0 | 0 |
| 21 | 0 | 0 | 0 | 0.3 | 0 | 0 | 0 |
| 22 | 0 | 0 | 0 | 0.3 | 0 | 0 | 0 |
| 23 | 0 | 0 | 0 | 0 | 0.3 | 0 | 0 |
| 24 | 0 | 0 | 0 | 0 | 0.3 | 0 | 0 |
| 25 | 0 | 0 | 0 | 0 | 0 | 0.3 | 0 |
| 26 | 0 | 0 | 0 | 0 | 0 | 0.3 | 0 |
| 27 | 0 | 0 | 0 | 0 | 0 | 0 | 0.3 |
| 28 | 0 | 0 | 0 | 0 | 0 | 0 | 0.3 |
| 29 | 0.3 | 0.3 | 0 | 0.3 | 0.3 | 0 | 0 |
| 30 | 0.3 | 0.3 | 0 | 0.3 | 0.3 | 0 | 0 |
| 31 | 0 | 0 | 0 | 0 | 0 | 0 | 0.3 |
| 32 | 0 | 0 | 0 | 0 | 0 | 0 | 0.3 |
| 33 | 0 | 0 | 0 | 0 | 0 | 0 | 0.3 |
| 34 | 0 | 0 | 0 | 0 | 0 | 0 | 0.3 |
| 35 | 0 | 0 | 0.3 | 0 | 0 | 0.3 | 0 |
| 36 | 0 | 0 | 0.3 | 0 | 0 | 0.3 | 0 |
| 37 | 0 | 0 | 0 | 0 | 0 | 0 | 0 |
| 38 | 0 | 0 | 0 | 0 | 0 | 0 | 0 |
| 39 | 0 | 0 | 0 | 0 | 0 | 0 | 0 |
| 40 | 0 | 0 | 0 | 0 | 0 | 0 | 0 |
| 41 | 0 | 0 | 0 | 0 | 0 | 0 | 0 |
| 42 | 0 | 0 | 0 | 0 | 0 | 0 | 0 |
| 43 | 0 | 0 | 0 | 0 | 0 | 0 | 0 |
| 44 | 0 | 0 | 0 | 0 | 0 | 0 | 0 |
| 45 | 0 | 0 | 0 | 0 | 0 | 0 | 0 |
| 46 | 0 | 0 | 0 | 0 | 0 | 0 | 0 |
| 47 | 0 | 0 | 0 | 0 | 0 | 0 | 0 |
| 48 | 0 | 0 | 0 | 0 | 0 | 0 | 0 |
| 49 | 0 | 0 | 0 | 0 | 0 | 0 | 0 |

**B2.6 Table: PFC → PFC Recurrent weight matrix**

Rows = Source PFC nodes; Columns = Target PFC nodes.

Continued on next page

| | 1 | 2 | 3 | 4 | 5 | 6 | 7 | 8 | 9 | 10 | 11 | 12 | 13 | 14 | 15 | 16 |
|---|---|---|---|---|---|---|---|---|---|---|---|---|---|---|---|---|
| 1 | 0.05 | 0.05 | -1 | -0.5 | 0.05 | 0.05 | -0.5 | -0.5 | 0.05 | 0.05 | -0.5 | -0.5 | 0.05 | 0.05 | 0.05 | 0.05 |
| 2 | 0.05 | 0.05 | -0.5 | -1 | 0.05 | 0.05 | -0.5 | -0.5 | 0.05 | 0.05 | -0.5 | -0.5 | 0.05 | 0.05 | 0.05 | 0.05 |
| 3 | -1 | -0.5 | 0.05 | 0.05 | -0.5 | -0.5 | 0.05 | 0.05 | 0.05 | 0.05 | 0.05 | 0.05 | 0.05 | 0.05 | -0.5 | -0.5 |
| 4 | -0.5 | -1 | 0.05 | 0.05 | -0.5 | -0.5 | 0.05 | 0.05 | 0.05 | 0.05 | 0.05 | 0.05 | 0.05 | 0.05 | -0.5 | -0.5 |
| 5 | 0.05 | 0.05 | -0.5 | -0.5 | 0.05 | 0.05 | -0.5 | -0.5 | -0.5 | -0.5 | -0.5 | -0.5 | -0.5 | -0.5 | 0.05 | 0.05 |
| 6 | 0.05 | 0.05 | -0.5 | -0.5 | 0.05 | 0.05 | -0.5 | -0.5 | -0.5 | -0.5 | -0.5 | -0.5 | -0.5 | -0.5 | 0.05 | 0.05 |
| 7 | -0.5 | -0.5 | 0.05 | 0.05 | -0.5 | -0.5 | 0.05 | 0.05 | -0.5 | -0.5 | -0.5 | -0.5 | -0.5 | -0.5 | -0.5 | -0.5 |
| 8 | -0.5 | -0.5 | 0.05 | 0.05 | -0.5 | -0.5 | 0.05 | 0.05 | -0.5 | -0.5 | -0.5 | -0.5 | -0.5 | -0.5 | -0.5 | -0.5 |
| 9 | 0.05 | 0.05 | 0.05 | 0.05 | -0.5 | -0.5 | -0.5 | -0.5 | 0.05 | 0.05 | -0.5 | -0.5 | -0.5 | -0.5 | -0.5 | -0.5 |
| 10 | 0.05 | 0.05 | 0.05 | 0.05 | -0.5 | -0.5 | -0.5 | -0.5 | 0.05 | 0.05 | -0.5 | -0.5 | -0.5 | -0.5 | -0.5 | -0.5 |
| 11 | -0.5 | -0.5 | 0.05 | 0.05 | -0.5 | -0.5 | -0.5 | -0.5 | -0.5 | -0.5 | 0.05 | 0.05 | -0.5 | -0.5 | -0.5 | -0.5 |
| 12 | -0.5 | -0.5 | 0.05 | 0.05 | -0.5 | -0.5 | -0.5 | -0.5 | -0.5 | -0.5 | 0.05 | 0.05 | -0.5 | -0.5 | -0.5 | -0.5 |
| 13 | 0.05 | 0.05 | 0.05 | 0.05 | -0.5 | -0.5 | -0.5 | -0.5 | -0.5 | -0.5 | -0.5 | -0.5 | 0.05 | 0.05 | -0.5 | -0.5 |
| 14 | 0.05 | 0.05 | 0.05 | 0.05 | -0.5 | -0.5 | -0.5 | -0.5 | -0.5 | -0.5 | -0.5 | -0.5 | 0.05 | 0.05 | -0.5 | -0.5 |
| 15 | 0.05 | 0.05 | -0.5 | -0.5 | 0.05 | 0.05 | -0.5 | -0.5 | -0.5 | -0.5 | -0.5 | -0.5 | -0.5 | -0.5 | 0.05 | 0.05 |
| 16 | 0.05 | 0.05 | -0.5 | -0.5 | 0.05 | 0.05 | -0.5 | -0.5 | -0.5 | -0.5 | -0.5 | -0.5 | -0.5 | -0.5 | 0.05 | 0.05 |
| 17 | 0.05 | 0.05 | 0.05 | 0.05 | -0.5 | -0.5 | 0.05 | 0.05 | -0.5 | -0.5 | -0.5 | -0.5 | 0.05 | 0.05 | -0.5 | -0.5 |
| 18 | 0.05 | 0.05 | 0.05 | 0.05 | -0.5 | -0.5 | 0.05 | 0.05 | -0.5 | -0.5 | -0.5 | -0.5 | 0.05 | 0.05 | -0.5 | -0.5 |
| 19 | -0.5 | -0.5 | 0.05 | 0.05 | -0.5 | -0.5 | 0.05 | 0.05 | -0.5 | -0.5 | -0.5 | -0.5 | -0.5 | -0.5 | -0.5 | -0.5 |
| 20 | -0.5 | -0.5 | 0.05 | 0.05 | -0.5 | -0.5 | 0.05 | 0.05 | -0.5 | -0.5 | -0.5 | -0.5 | -0.5 | -0.5 | -0.5 | -0.5 |
| 21 | 0.05 | 0.05 | 0.05 | 0.05 | -0.5 | -0.5 | -0.5 | -0.5 | 0.05 | 0.05 | -0.5 | -0.5 | -0.5 | -0.5 | -0.5 | -0.5 |
| 22 | 0.05 | 0.05 | 0.05 | 0.05 | -0.5 | -0.5 | -0.5 | -0.5 | 0.05 | 0.05 | -0.5 | -0.5 | -0.5 | -0.5 | -0.5 | -0.5 |
| 23 | -0.5 | -0.5 | 0.05 | 0.05 | -0.5 | -0.5 | -0.5 | -0.5 | -0.5 | -0.5 | 0.05 | 0.05 | -0.5 | -0.5 | -0.5 | -0.5 |
| 24 | -0.5 | -0.5 | 0.05 | 0.05 | -0.5 | -0.5 | -0.5 | -0.5 | -0.5 | -0.5 | 0.05 | 0.05 | -0.5 | -0.5 | -0.5 | -0.5 |
| 25 | 0.05 | 0.05 | 0.05 | 0.05 | -0.5 | -0.5 | -0.5 | -0.5 | -0.5 | -0.5 | -0.5 | -0.5 | 0.05 | 0.05 | -0.5 | -0.5 |
| 26 | 0.05 | 0.05 | 0.05 | 0.05 | -0.5 | -0.5 | -0.5 | -0.5 | -0.5 | -0.5 | -0.5 | -0.5 | 0.05 | 0.05 | -0.5 | -0.5 |
| 27 | 0.05 | 0.05 | 0.05 | 0.05 | 0.05 | 0.05 | 0.05 | 0.05 | 0.05 | 0.05 | 0.05 | 0.05 | 0.05 | 0.05 | 0.05 | 0.05 |
| 28 | 0.05 | 0.05 | 0.05 | 0.05 | 0.05 | 0.05 | 0.05 | 0.05 | 0.05 | 0.05 | 0.05 | 0.05 | 0.05 | 0.05 | 0.05 | 0.05 |
| 29 | 0.05 | 0.05 | 0.05 | 0.05 | 0.05 | 0.05 | 0.05 | 0.05 | 0.05 | 0.05 | 0.05 | 0.05 | 0.05 | 0.05 | 0.05 | 0.05 |
| 30 | 0.05 | 0.05 | 0.05 | 0.05 | 0.05 | 0.05 | 0.05 | 0.05 | 0.05 | 0.05 | 0.05 | 0.05 | 0.05 | 0.05 | 0.05 | 0.05 |
| 31 | 0.05 | 0.05 | -0.5 | -0.5 | 0.05 | 0.05 | -0.5 | -0.5 | -0.5 | -0.5 | -0.5 | -0.5 | -0.5 | -0.5 | 0.05 | 0.05 |
| 32 | 0.05 | 0.05 | -0.5 | -0.5 | 0.05 | 0.05 | -0.5 | -0.5 | -0.5 | -0.5 | -0.5 | -0.5 | -0.5 | -0.5 | 0.05 | 0.05 |
| 33 | 0.05 | 0.05 | 0.05 | 0.05 | -0.5 | -0.5 | 0.05 | 0.05 | 0.05 | 0.05 | 0.05 | 0.05 | 0.05 | 0.05 | -0.5 | -0.5 |
| 34 | 0.05 | 0.05 | 0.05 | 0.05 | -0.5 | -0.5 | 0.05 | 0.05 | 0.05 | 0.05 | 0.05 | 0.05 | 0.05 | 0.05 | -0.5 | -0.5 |
| 35 | 0.05 | 0.05 | 0.05 | 0.05 | -0.5 | -0.5 | 0.05 | 0.05 | -0.5 | -0.5 | -0.5 | -0.5 | 0.05 | 0.05 | -0.5 | -0.5 |
| 36 | 0.05 | 0.05 | 0.05 | 0.05 | -0.5 | -0.5 | 0.05 | 0.05 | -0.5 | -0.5 | -0.5 | -0.5 | 0.05 | 0.05 | -0.5 | -0.5 |
| 37 | -0.5 | -0.5 | 0.05 | 0.05 | -0.5 | -0.5 | -0.5 | -0.5 | -0.5 | -0.5 | 0.05 | 0.05 | -0.5 | -0.5 | -0.5 | -0.5 |
| 38 | -0.5 | -0.5 | 0.05 | 0.05 | -0.5 | -0.5 | -0.5 | -0.5 | -0.5 | -0.5 | 0.05 | 0.05 | -0.5 | -0.5 | -0.5 | -0.5 |
| 39 | -0.5 | -0.5 | 0.05 | 0.05 | -0.5 | -0.5 | -0.5 | -0.5 | -0.5 | -0.5 | -0.5 | -0.5 | 0.05 | 0.05 | -0.5 | -0.5 |
| 40 | -0.5 | -0.5 | 0.05 | 0.05 | -0.5 | -0.5 | -0.5 | -0.5 | -0.5 | -0.5 | -0.5 | -0.5 | 0.05 | 0.05 | -0.5 | -0.5 |
| 41 | -0.5 | -0.5 | 0.05 | 0.05 | -0.5 | -0.5 | -0.5 | -0.5 | -0.5 | -0.5 | 0.05 | 0.05 | -0.5 | -0.5 | -0.5 | -0.5 |
| 42 | -0.5 | -0.5 | 0.05 | 0.05 | -0.5 | -0.5 | -0.5 | -0.5 | -0.5 | -0.5 | 0.05 | 0.05 | -0.5 | -0.5 | -0.5 | -0.5 |
| 43 | -0.5 | -0.5 | 0.05 | 0.05 | -0.5 | -0.5 | -0.5 | -0.5 | -0.5 | -0.5 | -0.5 | -0.5 | 0.05 | 0.05 | -0.5 | -0.5 |
| 44 | -0.5 | -0.5 | 0.05 | 0.05 | -0.5 | -0.5 | -0.5 | -0.5 | -0.5 | -0.5 | -0.5 | -0.5 | 0.05 | 0.05 | -0.5 | -0.5 |
| 45 | -0.5 | -0.5 | -0.5 | -0.5 | -0.5 | -0.5 | -0.5 | -0.5 | -0.5 | -0.5 | -0.5 | -0.5 | -0.5 | -0.5 | -0.5 | -0.5 |
| 46 | -0.5 | -0.5 | -0.5 | -0.5 | -0.5 | -0.5 | -0.5 | -0.5 | -0.5 | -0.5 | -0.5 | -0.5 | -0.5 | -0.5 | -0.5 | -0.5 |
| 47 | -0.5 | -0.5 | -0.5 | -0.5 | -0.5 | -0.5 | -0.5 | -0.5 | -0.5 | -0.5 | -0.5 | -0.5 | -0.5 | -0.5 | -0.5 | -0.5 |
| 48 | -0.5 | -0.5 | -0.5 | -0.5 | -0.5 | -0.5 | -0.5 | -0.5 | -0.5 | -0.5 | -0.5 | -0.5 | -0.5 | -0.5 | -0.5 | -0.5 |
| 49 | -0.5 | -0.5 | -0.5 | -0.5 | -0.5 | -0.5 | -0.5 | -0.5 | -0.5 | -0.5 | -0.5 | -0.5 | -0.5 | -0.5 | -0.5 | -0.5 |

**B2.6 Table: PFC → PFC Recurrent weight matrix**

Continued from previous page. Continued on next page.

| | 17 | 18 | 19 | 20 | 21 | 22 | 23 | 24 | 25 | 26 | 27 | 28 | 29 | 30 | 31 | 32 | 33 |
|---|---|---|---|---|---|---|---|---|---|---|---|---|---|---|---|---|---|
| 1 | 0.05 | 0.05 | -0.5 | -0.5 | 0.05 | 0.05 | -0.5 | -0.5 | 0.05 | 0.05 | 0.05 | 0.05 | 0.05 | 0.05 | 0.05 | 0.05 | 0.05 |
| 2 | 0.05 | 0.05 | -0.5 | -0.5 | 0.05 | 0.05 | -0.5 | -0.5 | 0.05 | 0.05 | 0.05 | 0.05 | 0.05 | 0.05 | 0.05 | 0.05 | 0.05 |
| 3 | 0.05 | 0.05 | 0.05 | 0.05 | 0.05 | 0.05 | 0.05 | 0.05 | 0.05 | 0.05 | 0.05 | 0.05 | 0.05 | 0.05 | -0.5 | -0.5 | 0.05 |
| 4 | 0.05 | 0.05 | 0.05 | 0.05 | 0.05 | 0.05 | 0.05 | 0.05 | 0.05 | 0.05 | 0.05 | 0.05 | 0.05 | 0.05 | -0.5 | -0.5 | 0.05 |
| 5 | -0.5 | -0.5 | -0.5 | -0.5 | -0.5 | -0.5 | -0.5 | -0.5 | -0.5 | -0.5 | 0.05 | 0.05 | 0.05 | 0.05 | 0.05 | 0.05 | -0.5 |
| 6 | -0.5 | -0.5 | -0.5 | -0.5 | -0.5 | -0.5 | -0.5 | -0.5 | -0.5 | -0.5 | 0.05 | 0.05 | 0.05 | 0.05 | 0.05 | 0.05 | -0.5 |
| 7 | 0.05 | 0.05 | 0.05 | 0.05 | -0.5 | -0.5 | -0.5 | -0.5 | -0.5 | -0.5 | 0.05 | 0.05 | 0.05 | 0.05 | -0.5 | -0.5 | 0.05 |
| 8 | 0.05 | 0.05 | 0.05 | 0.05 | -0.5 | -0.5 | -0.5 | -0.5 | -0.5 | -0.5 | 0.05 | 0.05 | 0.05 | 0.05 | -0.5 | -0.5 | 0.05 |
| 9 | -0.5 | -0.5 | -0.5 | -0.5 | 0.05 | 0.05 | -0.5 | -0.5 | -0.5 | -0.5 | 0.05 | 0.05 | 0.05 | 0.05 | -0.5 | -0.5 | 0.05 |
| 10 | -0.5 | -0.5 | -0.5 | -0.5 | 0.05 | 0.05 | -0.5 | -0.5 | -0.5 | -0.5 | 0.05 | 0.05 | 0.05 | 0.05 | -0.5 | -0.5 | 0.05 |
| 11 | -0.5 | -0.5 | -0.5 | -0.5 | -0.5 | -0.5 | 0.05 | 0.05 | -0.5 | -0.5 | 0.05 | 0.05 | 0.05 | 0.05 | -0.5 | -0.5 | 0.05 |
| 12 | -0.5 | -0.5 | -0.5 | -0.5 | -0.5 | -0.5 | 0.05 | 0.05 | -0.5 | -0.5 | 0.05 | 0.05 | 0.05 | 0.05 | -0.5 | -0.5 | 0.05 |
| 13 | 0.05 | 0.05 | -0.5 | -0.5 | -0.5 | -0.5 | -0.5 | -0.5 | 0.05 | 0.05 | 0.05 | 0.05 | 0.05 | 0.05 | -0.5 | -0.5 | 0.05 |
| 14 | 0.05 | 0.05 | -0.5 | -0.5 | -0.5 | -0.5 | -0.5 | -0.5 | 0.05 | 0.05 | 0.05 | 0.05 | 0.05 | 0.05 | -0.5 | -0.5 | 0.05 |
| 15 | -0.5 | -0.5 | -0.5 | -0.5 | -0.5 | -0.5 | -0.5 | -0.5 | -0.5 | -0.5 | 0.05 | 0.05 | 0.05 | 0.05 | 0.05 | 0.05 | -0.5 |
| 16 | -0.5 | -0.5 | -0.5 | -0.5 | -0.5 | -0.5 | -0.5 | -0.5 | -0.5 | -0.5 | 0.05 | 0.05 | 0.05 | 0.05 | 0.05 | 0.05 | -0.5 |
| 17 | 0.05 | 0.05 | 0.05 | 0.05 | -0.5 | -0.5 | -0.5 | -0.5 | 0.05 | 0.05 | 0.05 | 0.05 | 0.05 | 0.05 | -0.5 | -0.5 | 0.05 |
| 18 | 0.05 | 0.05 | 0.05 | 0.05 | -0.5 | -0.5 | -0.5 | -0.5 | 0.05 | 0.05 | 0.05 | 0.05 | 0.05 | 0.05 | -0.5 | -0.5 | 0.05 |
| 19 | 0.05 | 0.05 | 0.05 | 0.05 | -0.5 | -0.5 | -0.5 | -0.5 | -0.5 | -0.5 | 0.05 | 0.05 | -0.5 | -0.5 | -0.5 | -0.5 | 0.05 |
| 20 | 0.05 | 0.05 | 0.05 | 0.05 | -0.5 | -0.5 | -0.5 | -0.5 | -0.5 | -0.5 | 0.05 | 0.05 | -0.5 | -0.5 | -0.5 | -0.5 | 0.05 |
| 21 | -0.5 | -0.5 | -0.5 | -0.5 | 0.05 | 0.05 | -0.5 | -0.5 | -0.5 | -0.5 | 0.05 | 0.05 | 0.05 | 0.05 | -0.5 | -0.5 | 0.05 |
| 22 | -0.5 | -0.5 | -0.5 | -0.5 | 0.05 | 0.05 | -0.5 | -0.5 | -0.5 | -0.5 | 0.05 | 0.05 | 0.05 | 0.05 | -0.5 | -0.5 | 0.05 |
| 23 | -0.5 | -0.5 | -0.5 | -0.5 | -0.5 | -0.5 | 0.05 | 0.05 | -0.5 | -0.5 | 0.05 | 0.05 | 0.05 | 0.05 | -0.5 | -0.5 | 0.05 |
| 24 | -0.5 | -0.5 | -0.5 | -0.5 | -0.5 | -0.5 | 0.05 | 0.05 | -0.5 | -0.5 | 0.05 | 0.05 | 0.05 | 0.05 | -0.5 | -0.5 | 0.05 |
| 25 | 0.05 | 0.05 | -0.5 | -0.5 | -0.5 | -0.5 | -0.5 | -0.5 | 0.05 | 0.05 | 0.05 | 0.05 | -0.5 | -0.5 | -0.5 | -0.5 | 0.05 |
| 26 | 0.05 | 0.05 | -0.5 | -0.5 | -0.5 | -0.5 | -0.5 | -0.5 | 0.05 | 0.05 | 0.05 | 0.05 | -0.5 | -0.5 | -0.5 | -0.5 | 0.05 |
| 27 | 0.05 | 0.05 | 0.05 | 0.05 | 0.05 | 0.05 | 0.05 | 0.05 | 0.05 | 0.05 | 0.05 | 0.05 | -0.5 | -0.5 | 0.05 | 0.05 | 0.05 |
| 28 | 0.05 | 0.05 | 0.05 | 0.05 | 0.05 | 0.05 | 0.05 | 0.05 | 0.05 | 0.05 | 0.05 | 0.05 | -0.5 | -0.5 | 0.05 | 0.05 | 0.05 |
| 29 | 0.05 | 0.05 | -0.5 | -0.5 | 0.05 | 0.05 | 0.05 | 0.05 | -0.5 | -0.5 | -1 | -1 | 0.05 | 0.05 | -0.5 | -0.5 | -1 |
| 30 | 0.05 | 0.05 | -0.5 | -0.5 | 0.05 | 0.05 | 0.05 | 0.05 | -0.5 | -0.5 | -1 | -1 | 0.05 | 0.05 | -0.5 | -0.5 | -1 |
| 31 | -0.5 | -0.5 | -0.5 | -0.5 | -0.5 | -0.5 | -0.5 | -0.5 | -0.5 | -0.5 | 0.05 | 0.05 | -0.5 | -0.5 | 0.05 | 0.05 | -0.5 |
| 32 | -0.5 | -0.5 | -0.5 | -0.5 | -0.5 | -0.5 | -0.5 | -0.5 | -0.5 | -0.5 | 0.05 | 0.05 | -0.5 | -0.5 | 0.05 | 0.05 | -0.5 |
| 33 | 0.05 | 0.05 | 0.05 | 0.05 | 0.05 | 0.05 | 0.05 | 0.05 | 0.05 | 0.05 | 0.05 | 0.05 | -0.5 | -0.5 | -0.5 | -0.5 | 0.05 |
| 34 | 0.05 | 0.05 | 0.05 | 0.05 | 0.05 | 0.05 | 0.05 | 0.05 | 0.05 | 0.05 | 0.05 | 0.05 | -0.5 | -0.5 | -0.5 | -0.5 | 0.05 |
| 35 | 0.05 | 0.05 | 0.05 | 0.05 | -0.5 | -0.5 | -0.5 | -0.5 | 0.05 | 0.05 | -1 | -1 | -0.5 | -0.5 | -0.5 | -0.5 | -1 |
| 36 | 0.05 | 0.05 | 0.05 | 0.05 | -0.5 | -0.5 | -0.5 | -0.5 | 0.05 | 0.05 | -1 | -1 | -0.5 | -0.5 | -0.5 | -0.5 | -1 |
| 37 | -0.5 | -0.5 | -0.5 | -0.5 | -0.5 | -0.5 | 0.05 | 0.05 | -0.5 | -0.5 | 0.05 | 0.05 | 0.05 | 0.05 | -0.5 | -0.5 | 0.05 |
| 38 | -0.5 | -0.5 | -0.5 | -0.5 | -0.5 | -0.5 | 0.05 | 0.05 | -0.5 | -0.5 | 0.05 | 0.05 | 0.05 | 0.05 | -0.5 | -0.5 | 0.05 |
| 39 | 0.05 | 0.05 | -0.5 | -0.5 | -0.5 | -0.5 | -0.5 | -0.5 | 0.05 | 0.05 | 0.05 | 0.05 | 0.05 | 0.05 | -0.5 | -0.5 | 0.05 |
| 40 | 0.05 | 0.05 | -0.5 | -0.5 | -0.5 | -0.5 | -0.5 | -0.5 | 0.05 | 0.05 | 0.05 | 0.05 | 0.05 | 0.05 | -0.5 | -0.5 | 0.05 |
| 41 | -0.5 | -0.5 | -0.5 | -0.5 | -0.5 | -0.5 | 0.05 | 0.05 | -0.5 | -0.5 | 0.05 | 0.05 | 0.05 | 0.05 | -0.5 | -0.5 | 0.05 |
| 42 | -0.5 | -0.5 | -0.5 | -0.5 | -0.5 | -0.5 | 0.05 | 0.05 | -0.5 | -0.5 | 0.05 | 0.05 | 0.05 | 0.05 | -0.5 | -0.5 | 0.05 |
| 43 | 0.05 | 0.05 | -0.5 | -0.5 | -0.5 | -0.5 | -0.5 | -0.5 | 0.05 | 0.05 | 0.05 | 0.05 | 0.05 | 0.05 | -0.5 | -0.5 | 0.05 |
| 44 | 0.05 | 0.05 | -0.5 | -0.5 | -0.5 | -0.5 | -0.5 | -0.5 | 0.05 | 0.05 | 0.05 | 0.05 | 0.05 | 0.05 | -0.5 | -0.5 | 0.05 |
| 45 | -0.5 | -0.5 | -0.5 | -0.5 | -0.5 | -0.5 | -0.5 | -0.5 | -0.5 | -0.5 | -0.5 | -0.5 | -0.5 | -0.5 | -0.5 | -0.5 | -0.5 |
| 46 | -0.5 | -0.5 | -0.5 | -0.5 | -0.5 | -0.5 | -0.5 | -0.5 | -0.5 | -0.5 | -0.5 | -0.5 | -0.5 | -0.5 | -0.5 | -0.5 | -0.5 |
| 47 | -0.5 | -0.5 | -0.5 | -0.5 | -0.5 | -0.5 | -0.5 | -0.5 | -0.5 | -0.5 | -0.5 | -0.5 | -0.5 | -0.5 | -0.5 | -0.5 | -0.5 |
| 48 | -0.5 | -0.5 | -0.5 | -0.5 | -0.5 | -0.5 | -0.5 | -0.5 | -0.5 | -0.5 | -0.5 | -0.5 | -0.5 | -0.5 | -0.5 | -0.5 | -0.5 |
| 49 | -0.5 | -0.5 | -0.5 | -0.5 | -0.5 | -0.5 | -0.5 | -0.5 | -0.5 | -0.5 | -0.5 | -0.5 | -0.5 | -0.5 | -0.5 | -0.5 | -0.5 |

**B2.6 Table: PFC → PFC Recurrent weight matrix**

Continued from previous page.

| | 34 | 35 | 36 | 37 | 38 | 39 | 40 | 41 | 42 | 43 | 44 | 45 | 46 | 47 | 48 | 49 |
|---|---|---|---|---|---|---|---|---|---|---|---|---|---|---|---|---|
| 1 | 0.05 | 0.05 | 0.05 | -1 | -1 | -1 | -1 | -1 | -1 | -1 | -1 | -0.5 | -0.5 | -0.5 | -0.5 | -0.5 |
| 2 | 0.05 | 0.05 | 0.05 | -1 | -1 | -1 | -1 | -1 | -1 | -1 | -1 | -0.5 | -0.5 | -0.5 | -0.5 | -0.5 |
| 3 | 0.05 | 0.05 | 0.05 | 0.05 | 0.05 | 0.05 | 0.05 | 0.05 | 0.05 | 0.05 | 0.05 | -0.5 | -0.5 | -0.5 | -0.5 | -0.5 |
| 4 | 0.05 | 0.05 | 0.05 | 0.05 | 0.05 | 0.05 | 0.05 | 0.05 | 0.05 | 0.05 | 0.05 | -0.5 | -0.5 | -0.5 | -0.5 | -0.5 |
| 5 | -0.5 | -0.5 | -0.5 | -0.5 | -0.5 | -0.5 | -0.5 | -0.5 | -0.5 | -0.5 | -0.5 | -0.5 | -0.5 | -0.5 | -0.5 | -0.5 |
| 6 | -0.5 | -0.5 | -0.5 | -0.5 | -0.5 | -0.5 | -0.5 | -0.5 | -0.5 | -0.5 | -0.5 | -0.5 | -0.5 | -0.5 | -0.5 | -0.5 |
| 7 | 0.05 | 0.05 | 0.05 | -0.5 | -0.5 | -0.5 | -0.5 | -0.5 | -0.5 | -0.5 | -0.5 | -0.5 | -0.5 | -0.5 | -0.5 | -0.5 |
| 8 | 0.05 | 0.05 | 0.05 | -0.5 | -0.5 | -0.5 | -0.5 | -0.5 | -0.5 | -0.5 | -0.5 | -0.5 | -0.5 | -0.5 | -0.5 | -0.5 |
| 9 | 0.05 | -0.5 | -0.5 | -0.5 | -0.5 | -0.5 | -0.5 | -0.5 | -0.5 | -0.5 | -0.5 | -0.5 | -0.5 | -0.5 | -0.5 | -0.5 |
| 10 | 0.05 | -0.5 | -0.5 | -0.5 | -0.5 | -0.5 | -0.5 | -0.5 | -0.5 | -0.5 | -0.5 | -0.5 | -0.5 | -0.5 | -0.5 | -0.5 |
| 11 | 0.05 | -0.5 | -0.5 | 0.05 | 0.05 | -0.5 | -0.5 | 0.05 | 0.05 | -0.5 | -0.5 | -0.5 | -0.5 | -0.5 | -0.5 | -0.5 |
| 12 | 0.05 | -0.5 | -0.5 | 0.05 | 0.05 | -0.5 | -0.5 | 0.05 | 0.05 | -0.5 | -0.5 | -0.5 | -0.5 | -0.5 | -0.5 | -0.5 |
| 13 | 0.05 | 0.05 | 0.05 | -0.5 | -0.5 | 0.05 | 0.05 | -0.5 | -0.5 | 0.05 | 0.05 | -0.5 | -0.5 | -0.5 | -0.5 | -0.5 |
| 14 | 0.05 | 0.05 | 0.05 | -0.5 | -0.5 | 0.05 | 0.05 | -0.5 | -0.5 | 0.05 | 0.05 | -0.5 | -0.5 | -0.5 | -0.5 | -0.5 |
| 15 | -0.5 | -0.5 | -0.5 | -0.5 | -0.5 | -0.5 | -0.5 | -0.5 | -0.5 | -0.5 | -0.5 | -0.5 | -0.5 | -0.5 | -0.5 | -0.5 |
| 16 | -0.5 | -0.5 | -0.5 | -0.5 | -0.5 | -0.5 | -0.5 | -0.5 | -0.5 | -0.5 | -0.5 | -0.5 | -0.5 | -0.5 | -0.5 | -0.5 |
| 17 | 0.05 | 0.05 | 0.05 | -0.5 | -0.5 | 0.05 | 0.05 | -0.5 | -0.5 | 0.05 | 0.05 | -0.5 | -0.5 | -0.5 | -0.5 | -0.5 |
| 18 | 0.05 | 0.05 | 0.05 | -0.5 | -0.5 | 0.05 | 0.05 | -0.5 | -0.5 | 0.05 | 0.05 | -0.5 | -0.5 | -0.5 | -0.5 | -0.5 |
| 19 | 0.05 | 0.05 | 0.05 | -0.5 | -0.5 | -0.5 | -0.5 | -0.5 | -0.5 | -0.5 | -0.5 | -0.5 | -0.5 | -0.5 | -0.5 | -0.5 |
| 20 | 0.05 | 0.05 | 0.05 | -0.5 | -0.5 | -0.5 | -0.5 | -0.5 | -0.5 | -0.5 | -0.5 | -0.5 | -0.5 | -0.5 | -0.5 | -0.5 |
| 21 | 0.05 | -0.5 | -0.5 | -0.5 | -0.5 | -0.5 | -0.5 | -0.5 | -0.5 | -0.5 | -0.5 | -0.5 | -0.5 | -0.5 | -0.5 | -0.5 |
| 22 | 0.05 | -0.5 | -0.5 | -0.5 | -0.5 | -0.5 | -0.5 | -0.5 | -0.5 | -0.5 | -0.5 | -0.5 | -0.5 | -0.5 | -0.5 | -0.5 |
| 23 | 0.05 | -0.5 | -0.5 | 0.05 | 0.05 | -0.5 | -0.5 | 0.05 | 0.05 | -0.5 | -0.5 | -0.5 | -0.5 | -0.5 | -0.5 | -0.5 |
| 24 | 0.05 | -0.5 | -0.5 | 0.05 | 0.05 | -0.5 | -0.5 | 0.05 | 0.05 | -0.5 | -0.5 | -0.5 | -0.5 | -0.5 | -0.5 | -0.5 |
| 25 | 0.05 | 0.05 | 0.05 | -0.5 | -0.5 | 0.05 | 0.05 | -0.5 | -0.5 | 0.05 | 0.05 | -0.5 | -0.5 | -0.5 | -0.5 | -0.5 |
| 26 | 0.05 | 0.05 | 0.05 | -0.5 | -0.5 | 0.05 | 0.05 | -0.5 | -0.5 | 0.05 | 0.05 | -0.5 | -0.5 | -0.5 | -0.5 | -0.5 |
| 27 | 0.05 | -0.5 | -0.5 | 0.05 | 0.05 | 0.05 | 0.05 | 0.05 | 0.05 | 0.05 | 0.05 | -0.5 | -0.5 | -0.5 | -0.5 | -0.5 |
| 28 | 0.05 | -0.5 | -0.5 | 0.05 | 0.05 | 0.05 | 0.05 | 0.05 | 0.05 | 0.05 | 0.05 | -0.5 | -0.5 | -0.5 | -0.5 | -0.5 |
| 29 | -1 | -1 | -0.5 | 0.05 | 0.05 | 0.05 | 0.05 | 0.05 | 0.05 | 0.05 | 0.05 | -0.5 | -0.5 | -0.5 | -0.5 | -0.5 |
| 30 | -1 | -0.5 | -1 | 0.05 | 0.05 | 0.05 | 0.05 | 0.05 | 0.05 | 0.05 | 0.05 | -0.5 | -0.5 | -0.5 | -0.5 | -0.5 |
| 31 | -0.5 | -0.5 | -0.5 | -0.5 | -0.5 | -0.5 | -0.5 | -0.5 | -0.5 | -0.5 | -0.5 | -0.5 | -0.5 | -0.5 | -0.5 | -0.5 |
| 32 | -0.5 | -0.5 | -0.5 | -0.5 | -0.5 | -0.5 | -0.5 | -0.5 | -0.5 | -0.5 | -0.5 | -0.5 | -0.5 | -0.5 | -0.5 | -0.5 |
| 33 | 0.05 | -0.5 | -0.5 | 0.05 | 0.05 | 0.05 | 0.05 | 0.05 | 0.05 | 0.05 | 0.05 | -0.5 | -0.5 | -0.5 | -0.5 | -0.5 |
| 34 | 0.05 | -0.5 | -0.5 | 0.05 | 0.05 | 0.05 | 0.05 | 0.05 | 0.05 | 0.05 | 0.05 | -0.5 | -0.5 | -0.5 | -0.5 | -0.5 |
| 35 | -1 | 0.05 | 0.05 | -0.5 | -0.5 | 0.05 | 0.05 | -0.5 | -0.5 | 0.05 | 0.05 | -0.5 | -0.5 | -0.5 | -0.5 | -0.5 |
| 36 | -1 | 0.05 | 0.05 | -0.5 | -0.5 | 0.05 | 0.05 | -0.5 | -0.5 | 0.05 | 0.05 | -0.5 | -0.5 | -0.5 | -0.5 | -0.5 |
| 37 | 0.05 | -0.5 | -0.5 | 0.05 | 0.05 | -0.5 | -0.5 | -1 | -0.5 | -0.5 | -0.5 | -0.5 | -0.5 | -0.5 | -0.5 | -0.5 |
| 38 | 0.05 | -0.5 | -0.5 | 0.05 | 0.05 | -0.5 | -0.5 | -0.5 | -1 | -0.5 | -0.5 | -0.5 | -0.5 | -0.5 | -0.5 | -0.5 |
| 39 | 0.05 | 0.05 | 0.05 | -0.5 | -0.5 | 0.05 | 0.05 | -0.5 | -0.5 | -1 | -0.5 | -0.5 | -0.5 | -0.5 | -0.5 | -0.5 |
| 40 | 0.05 | 0.05 | 0.05 | -0.5 | -0.5 | 0.05 | 0.05 | -0.5 | -0.5 | -0.5 | -1 | -0.5 | -0.5 | -0.5 | -0.5 | -0.5 |
| 41 | 0.05 | -0.5 | -0.5 | -1 | -0.5 | -0.5 | -0.5 | 0.05 | 0.05 | -0.5 | -0.5 | -0.5 | -0.5 | -0.5 | -0.5 | -0.5 |
| 42 | 0.05 | -0.5 | -0.5 | -0.5 | -1 | -0.5 | -0.5 | 0.05 | 0.05 | -0.5 | -0.5 | -0.5 | -0.5 | -0.5 | -0.5 | -0.5 |
| 43 | 0.05 | 0.05 | 0.05 | -0.5 | -0.5 | -1 | -0.5 | -0.5 | -0.5 | 0.05 | 0.05 | -0.5 | -0.5 | -0.5 | -0.5 | -0.5 |
| 44 | 0.05 | 0.05 | 0.05 | -0.5 | -0.5 | -0.5 | -1 | -0.5 | -0.5 | 0.05 | 0.05 | -0.5 | -0.5 | -0.5 | -0.5 | -0.5 |
| 45 | -0.5 | -0.5 | -0.5 | -0.5 | -0.5 | -0.5 | -0.5 | -0.5 | -0.5 | -0.5 | -0.5 | -0.5 | -0.5 | -0.5 | -0.5 | -0.5 |
| 46 | -0.5 | -0.5 | -0.5 | -0.5 | -0.5 | -0.5 | -0.5 | -0.5 | -0.5 | -0.5 | -0.5 | -0.5 | -0.5 | -0.5 | -0.5 | -0.5 |
| 47 | -0.5 | -0.5 | -0.5 | -0.5 | -0.5 | -0.5 | -0.5 | -0.5 | -0.5 | -0.5 | -0.5 | -0.5 | -0.5 | -0.5 | -0.5 | -0.5 |
| 48 | -0.5 | -0.5 | -0.5 | -0.5 | -0.5 | -0.5 | -0.5 | -0.5 | -0.5 | -0.5 | -0.5 | -0.5 | -0.5 | -0.5 | -0.5 | -0.5 |
| 49 | -0.5 | -0.5 | -0.5 | -0.5 | -0.5 | -0.5 | -0.5 | -0.5 | -0.5 | -0.5 | -0.5 | -0.5 | -0.5 | -0.5 | -0.5 | -0.5 |

**B2.7 Table: PFC → Transition node weight matrix**

Rows = Source PFC nodes; Columns = Target transition nodes.

Continued on next page.

| | 1 | 2 | 3 | 4 | 5 | 6 | 7 | 8 | 9 | 10 | 11 |
|---|---|---|---|---|---|---|---|---|---|---|---|
| 1 | -0.005 | 0.03 | -0.005 | -0.005 | -0.005 | 0.03 | 0.03 | -0.005 | -0.005 | -0.005 | -0.005 |
| 2 | -0.005 | 0.03 | -0.005 | -0.005 | -0.005 | 0.03 | 0.03 | -0.005 | -0.005 | -0.005 | -0.005 |
| 3 | -0.005 | -0.005 | -0.005 | 0.03 | 0.03 | -0.005 | -0.005 | 0.03 | 0.03 | 0.03 | 0.03 |
| 4 | -0.005 | -0.005 | -0.005 | 0.03 | 0.03 | -0.005 | -0.005 | 0.03 | 0.03 | 0.03 | 0.03 |
| 5 | -0.005 | 0.03 | -0.005 | -0.005 | -0.005 | 0.03 | -0.005 | -0.005 | -0.005 | -0.005 | -0.005 |
| 6 | -0.005 | 0.03 | -0.005 | -0.005 | -0.005 | 0.03 | -0.005 | -0.005 | -0.005 | -0.005 | -0.005 |
| 7 | -0.005 | -0.005 | -0.005 | 0.03 | 0.03 | -0.005 | -0.005 | 0.03 | -0.005 | -0.005 | -0.005 |
| 8 | -0.005 | -0.005 | -0.005 | 0.03 | 0.03 | -0.005 | -0.005 | 0.03 | -0.005 | -0.005 | -0.005 |
| 9 | -0.005 | -0.005 | -0.005 | -0.005 | -0.005 | -0.005 | 0.03 | -0.005 | 0.03 | 0.03 | -0.005 |
| 10 | -0.005 | -0.005 | -0.005 | -0.005 | -0.005 | -0.005 | 0.03 | -0.005 | 0.03 | 0.03 | -0.005 |
| 11 | -0.005 | -0.005 | -0.005 | -0.005 | -0.005 | -0.005 | -0.005 | -0.005 | -0.005 | -0.005 | 0.03 |
| 12 | -0.005 | -0.005 | -0.005 | -0.005 | -0.005 | -0.005 | -0.005 | -0.005 | -0.005 | -0.005 | 0.03 |
| 13 | -0.005 | -0.005 | -0.005 | -0.005 | -0.005 | -0.005 | -0.005 | -0.005 | -0.005 | -0.005 | -0.005 |
| 14 | -0.005 | -0.005 | -0.005 | -0.005 | -0.005 | -0.005 | -0.005 | -0.005 | -0.005 | -0.005 | -0.005 |
| 15 | -0.005 | 0.03 | -0.005 | -0.005 | -0.005 | 0.03 | -0.005 | -0.005 | -0.005 | -0.005 | -0.005 |
| 16 | -0.005 | 0.03 | -0.005 | -0.005 | -0.005 | 0.03 | -0.005 | -0.005 | -0.005 | -0.005 | -0.005 |
| 17 | -0.005 | -0.005 | -0.005 | 0.03 | 0.03 | -0.005 | -0.005 | 0.03 | -0.005 | -0.005 | -0.005 |
| 18 | -0.005 | -0.005 | -0.005 | 0.03 | 0.03 | -0.005 | -0.005 | 0.03 | -0.005 | -0.005 | -0.005 |
| 19 | -0.005 | -0.005 | -0.005 | -0.005 | 0.03 | -0.005 | -0.005 | 0.03 | -0.005 | -0.005 | -0.005 |
| 20 | -0.005 | -0.005 | -0.005 | -0.005 | 0.03 | -0.005 | -0.005 | 0.03 | -0.005 | -0.005 | -0.005 |
| 21 | -0.005 | -0.005 | -0.005 | -0.005 | -0.005 | -0.005 | 0.03 | -0.005 | 0.03 | 0.03 | -0.005 |
| 22 | -0.005 | -0.005 | -0.005 | -0.005 | -0.005 | -0.005 | 0.03 | -0.005 | 0.03 | 0.03 | -0.005 |
| 23 | -0.005 | -0.005 | -0.005 | -0.005 | -0.005 | -0.005 | -0.005 | -0.005 | -0.005 | -0.005 | 0.03 |
| 24 | -0.005 | -0.005 | -0.005 | -0.005 | -0.005 | -0.005 | -0.005 | -0.005 | -0.005 | -0.005 | 0.03 |
| 25 | -0.005 | -0.005 | -0.005 | -0.005 | -0.005 | -0.005 | -0.005 | -0.005 | -0.005 | -0.005 | -0.005 |
| 26 | -0.005 | -0.005 | -0.005 | -0.005 | -0.005 | -0.005 | -0.005 | -0.005 | -0.005 | -0.005 | -0.005 |
| 27 | -0.005 | -0.005 | -0.005 | -0.005 | -0.005 | 0.03 | -0.005 | 0.03 | -0.005 | 0.03 | -0.005 |
| 28 | -0.005 | -0.005 | -0.005 | -0.005 | -0.005 | 0.03 | -0.005 | 0.03 | -0.005 | 0.03 | -0.005 |
| 29 | -0.005 | 0.03 | -0.005 | 0.03 | -0.005 | -0.005 | 0.03 | -0.005 | 0.03 | -0.005 | 0.03 |
| 30 | -0.005 | 0.03 | -0.005 | 0.03 | -0.005 | -0.005 | 0.03 | -0.005 | 0.03 | -0.005 | 0.03 |
| 31 | -0.005 | -0.005 | -0.005 | -0.005 | -0.005 | 0.03 | -0.005 | -0.005 | -0.005 | -0.005 | -0.005 |
| 32 | -0.005 | -0.005 | -0.005 | -0.005 | -0.005 | 0.03 | -0.005 | -0.005 | -0.005 | -0.005 | -0.005 |
| 33 | -0.005 | -0.005 | -0.005 | -0.005 | -0.005 | -0.005 | -0.005 | 0.03 | -0.005 | 0.03 | -0.005 |
| 34 | -0.005 | -0.005 | -0.005 | -0.005 | -0.005 | -0.005 | -0.005 | 0.03 | -0.005 | 0.03 | -0.005 |
| 35 | -0.005 | -0.005 | -0.005 | -0.005 | 0.03 | -0.005 | -0.005 | -0.005 | -0.005 | -0.005 | -0.005 |
| 36 | -0.005 | -0.005 | -0.005 | -0.005 | 0.03 | -0.005 | -0.005 | -0.005 | -0.005 | -0.005 | -0.005 |
| 37 | -0.005 | -0.005 | -0.005 | -0.005 | -0.005 | -0.005 | -0.005 | -0.005 | -0.005 | -0.005 | 0.03 |
| 38 | -0.005 | -0.005 | -0.005 | -0.005 | -0.005 | -0.005 | -0.005 | -0.005 | -0.005 | -0.005 | 0.03 |
| 39 | -0.005 | -0.005 | -0.005 | -0.005 | -0.005 | -0.005 | -0.005 | -0.005 | -0.005 | -0.005 | -0.005 |
| 40 | -0.005 | -0.005 | -0.005 | -0.005 | -0.005 | -0.005 | -0.005 | -0.005 | -0.005 | -0.005 | -0.005 |
| 41 | -0.005 | -0.005 | -0.005 | -0.005 | -0.005 | -0.005 | -0.005 | -0.005 | -0.005 | -0.005 | -0.005 |
| 42 | -0.005 | -0.005 | -0.005 | -0.005 | -0.005 | -0.005 | -0.005 | -0.005 | -0.005 | -0.005 | -0.005 |
| 43 | -0.005 | -0.005 | -0.005 | -0.005 | -0.005 | -0.005 | -0.005 | -0.005 | -0.005 | -0.005 | -0.005 |
| 44 | -0.005 | -0.005 | -0.005 | -0.005 | -0.005 | -0.005 | -0.005 | -0.005 | -0.005 | -0.005 | -0.005 |
| 45 | -0.005 | -0.005 | -0.005 | -0.005 | -0.005 | -0.005 | -0.005 | -0.005 | -0.005 | -0.005 | -0.005 |
| 46 | -0.005 | -0.005 | -0.005 | -0.005 | -0.005 | -0.005 | -0.005 | -0.005 | -0.005 | -0.005 | -0.005 |
| 47 | -0.005 | -0.005 | -0.005 | -0.005 | -0.005 | -0.005 | -0.005 | -0.005 | -0.005 | -0.005 | -0.005 |
| 48 | -0.005 | -0.005 | -0.005 | -0.005 | -0.005 | -0.005 | -0.005 | -0.005 | -0.005 | -0.005 | -0.005 |
| 49 | -0.005 | -0.005 | -0.005 | -0.005 | -0.005 | -0.005 | -0.005 | -0.005 | -0.005 | -0.005 | -0.005 |

**B2.7 Table: PFC → Transition node weight matrix**

Continued from previous page.

| | 12 | 13 | 14 | 15 | 16 | 17 | 18 | 19 | 20 | 21 | 22 |
|---|---|---|---|---|---|---|---|---|---|---|---|
| 1 | -0.005 | -0.005 | 0.03 | 0.03 | 0.03 | -0.005 | -0.005 | -0.005 | -0.005 | -0.005 | -0.005 |
| 2 | -0.005 | -0.005 | 0.03 | 0.03 | 0.03 | -0.005 | -0.005 | -0.005 | -0.005 | -0.005 | -0.005 |
| 3 | 0.03 | 0.03 | -0.005 | -0.005 | -0.005 | 0.03 | 0.03 | 0.03 | 0.03 | 0.03 | 0.03 |
| 4 | 0.03 | 0.03 | -0.005 | -0.005 | -0.005 | 0.03 | 0.03 | 0.03 | 0.03 | 0.03 | 0.03 |
| 5 | -0.005 | -0.005 | -0.005 | -0.005 | -0.005 | -0.005 | -0.005 | -0.005 | -0.005 | -0.005 | -0.005 |
| 6 | -0.005 | -0.005 | -0.005 | -0.005 | -0.005 | -0.005 | -0.005 | -0.005 | -0.005 | -0.005 | -0.005 |
| 7 | -0.005 | -0.005 | -0.005 | -0.005 | -0.005 | -0.005 | -0.005 | -0.005 | -0.005 | -0.005 | -0.005 |
| 8 | -0.005 | -0.005 | -0.005 | -0.005 | -0.005 | -0.005 | -0.005 | -0.005 | -0.005 | -0.005 | -0.005 |
| 9 | -0.005 | -0.005 | 0.03 | -0.005 | -0.005 | 0.03 | -0.005 | -0.005 | -0.005 | -0.005 | -0.005 |
| 10 | -0.005 | -0.005 | 0.03 | -0.005 | -0.005 | 0.03 | -0.005 | -0.005 | -0.005 | -0.005 | -0.005 |
| 11 | -0.005 | 0.03 | -0.005 | -0.005 | -0.005 | -0.005 | -0.005 | -0.005 | 0.03 | -0.005 | -0.005 |
| 12 | -0.005 | 0.03 | -0.005 | -0.005 | -0.005 | -0.005 | -0.005 | -0.005 | 0.03 | -0.005 | -0.005 |
| 13 | 0.03 | -0.005 | -0.005 | 0.03 | 0.03 | -0.005 | 0.03 | 0.03 | -0.005 | 0.03 | 0.03 |
| 14 | 0.03 | -0.005 | -0.005 | 0.03 | 0.03 | -0.005 | 0.03 | 0.03 | -0.005 | 0.03 | 0.03 |
| 15 | -0.005 | -0.005 | -0.005 | -0.005 | -0.005 | -0.005 | -0.005 | -0.005 | -0.005 | -0.005 | -0.005 |
| 16 | -0.005 | -0.005 | -0.005 | -0.005 | -0.005 | -0.005 | -0.005 | -0.005 | -0.005 | -0.005 | -0.005 |
| 17 | 0.03 | -0.005 | -0.005 | 0.03 | 0.03 | -0.005 | 0.03 | 0.03 | -0.005 | 0.03 | 0.03 |
| 18 | 0.03 | -0.005 | -0.005 | 0.03 | 0.03 | -0.005 | 0.03 | 0.03 | -0.005 | 0.03 | 0.03 |
| 19 | -0.005 | -0.005 | -0.005 | -0.005 | -0.005 | -0.005 | -0.005 | -0.005 | -0.005 | -0.005 | -0.005 |
| 20 | -0.005 | -0.005 | -0.005 | -0.005 | -0.005 | -0.005 | -0.005 | -0.005 | -0.005 | -0.005 | -0.005 |
| 21 | -0.005 | -0.005 | 0.03 | -0.005 | -0.005 | 0.03 | -0.005 | -0.005 | -0.005 | -0.005 | -0.005 |
| 22 | -0.005 | -0.005 | 0.03 | -0.005 | -0.005 | 0.03 | -0.005 | -0.005 | -0.005 | -0.005 | -0.005 |
| 23 | -0.005 | 0.03 | -0.005 | -0.005 | -0.005 | -0.005 | -0.005 | -0.005 | 0.03 | -0.005 | -0.005 |
| 24 | -0.005 | 0.03 | -0.005 | -0.005 | -0.005 | -0.005 | -0.005 | -0.005 | 0.03 | -0.005 | -0.005 |
| 25 | 0.03 | -0.005 | -0.005 | -0.005 | 0.03 | -0.005 | -0.005 | 0.03 | -0.005 | -0.005 | 0.03 |
| 26 | 0.03 | -0.005 | -0.005 | -0.005 | 0.03 | -0.005 | -0.005 | 0.03 | -0.005 | -0.005 | 0.03 |
| 27 | 0.03 | -0.005 | 0.03 | -0.005 | -0.005 | 0.03 | -0.005 | -0.005 | 0.03 | -0.005 | -0.005 |
| 28 | 0.03 | -0.005 | 0.03 | -0.005 | -0.005 | 0.03 | -0.005 | -0.005 | 0.03 | -0.005 | -0.005 |
| 29 | -0.005 | 0.03 | -0.005 | 0.03 | -0.005 | -0.005 | 0.03 | -0.005 | -0.005 | 0.03 | -0.005 |
| 30 | -0.005 | 0.03 | -0.005 | 0.03 | -0.005 | -0.005 | 0.03 | -0.005 | -0.005 | 0.03 | -0.005 |
| 31 | -0.005 | -0.005 | -0.005 | -0.005 | -0.005 | -0.005 | -0.005 | -0.005 | -0.005 | -0.005 | -0.005 |
| 32 | -0.005 | -0.005 | -0.005 | -0.005 | -0.005 | -0.005 | -0.005 | -0.005 | -0.005 | -0.005 | -0.005 |
| 33 | 0.03 | -0.005 | 0.03 | -0.005 | -0.005 | 0.03 | -0.005 | -0.005 | 0.03 | -0.005 | -0.005 |
| 34 | 0.03 | -0.005 | 0.03 | -0.005 | -0.005 | 0.03 | -0.005 | -0.005 | 0.03 | -0.005 | -0.005 |
| 35 | -0.005 | -0.005 | -0.005 | -0.005 | 0.03 | -0.005 | -0.005 | 0.03 | -0.005 | -0.005 | 0.03 |
| 36 | -0.005 | -0.005 | -0.005 | -0.005 | 0.03 | -0.005 | -0.005 | 0.03 | -0.005 | -0.005 | 0.03 |
| 37 | -0.005 | -0.005 | -0.005 | -0.005 | -0.005 | -0.005 | -0.005 | -0.005 | 0.03 | -0.005 | -0.005 |
| 38 | -0.005 | -0.005 | -0.005 | -0.005 | -0.005 | -0.005 | -0.005 | -0.005 | 0.03 | -0.005 | -0.005 |
| 39 | -0.005 | -0.005 | -0.005 | -0.005 | -0.005 | -0.005 | -0.005 | -0.005 | -0.005 | 0.03 | 0.03 |
| 40 | -0.005 | -0.005 | -0.005 | -0.005 | -0.005 | -0.005 | -0.005 | -0.005 | -0.005 | 0.03 | 0.03 |
| 41 | -0.005 | 0.03 | -0.005 | -0.005 | -0.005 | -0.005 | -0.005 | -0.005 | -0.005 | -0.005 | -0.005 |
| 42 | -0.005 | 0.03 | -0.005 | -0.005 | -0.005 | -0.005 | -0.005 | -0.005 | -0.005 | -0.005 | -0.005 |
| 43 | 0.03 | -0.005 | -0.005 | -0.005 | -0.005 | -0.005 | 0.03 | 0.03 | -0.005 | -0.005 | -0.005 |
| 44 | 0.03 | -0.005 | -0.005 | -0.005 | -0.005 | -0.005 | 0.03 | 0.03 | -0.005 | -0.005 | -0.005 |
| 45 | -0.005 | -0.005 | -0.005 | -0.005 | -0.005 | -0.005 | -0.005 | -0.005 | -0.005 | -0.005 | -0.005 |
| 46 | -0.005 | -0.005 | -0.005 | -0.005 | -0.005 | -0.005 | -0.005 | -0.005 | -0.005 | -0.005 | -0.005 |
| 47 | -0.005 | -0.005 | -0.005 | -0.005 | -0.005 | -0.005 | -0.005 | -0.005 | -0.005 | -0.005 | -0.005 |
| 48 | -0.005 | -0.005 | -0.005 | -0.005 | -0.005 | -0.005 | -0.005 | -0.005 | -0.005 | -0.005 | -0.005 |
| 49 | -0.005 | -0.005 | -0.005 | -0.005 | -0.005 | -0.005 | -0.005 | -0.005 | -0.005 | -0.005 | -0.005 |

**B2.8 Table: Transition node → PFC weight matrix**

Rows = Source transition nodes. Columns = Target PFC nodes. Continued below.

| | 1 | 2 | 3 | 4 | 5 | 6 | 7 | 8 | 9 | 10 | 11 | 12 | 13 | 14 | 15 | 16 | 17 | 18 | 19 | 20 | 21 | 22 | 23 | 24 | 25 |
|---|---|---|---|---|---|---|---|---|---|---|---|---|---|---|---|---|---|---|---|---|---|---|---|---|---|
| 1 | 2 | 2 | -0.3 | -0.3 | 2 | 2 | -0.3 | -0.3 | -0.3 | -0.3 | -0.3 | -0.3 | -0.3 | -0.3 | 2 | 2 | -0.3 | -0.3 | -0.3 | -0.3 | -0.3 | -0.3 | -0.3 | -0.3 | -0.3 |
| 2 | 2 | 2 | -0.3 | -0.3 | 2 | 2 | -0.3 | -0.3 | -0.3 | -0.3 | -0.3 | -0.3 | -0.3 | -0.3 | 2 | 2 | -0.3 | -0.3 | -0.3 | -0.3 | -0.3 | -0.3 | -0.3 | -0.3 | -0.3 |
| 3 | -0.3 | -0.3 | 2 | 2 | -0.3 | -0.3 | 2 | 2 | -0.3 | -0.3 | -0.3 | -0.3 | -0.3 | -0.3 | -0.3 | -0.3 | 2 | 2 | -0.3 | -0.3 | -0.3 | -0.3 | -0.3 | -0.3 | -0.3 |
| 4 | -0.3 | -0.3 | 2 | 2 | -0.3 | -0.3 | 2 | 2 | -0.3 | -0.3 | -0.3 | -0.3 | -0.3 | -0.3 | -0.3 | -0.3 | 2 | 2 | 2 | 2 | -0.3 | -0.3 | -0.3 | -0.3 | -0.3 |
| 5 | -0.3 | -0.3 | 2 | 2 | -0.3 | -0.3 | 2 | 2 | -0.3 | -0.3 | -0.3 | -0.3 | -0.3 | -0.3 | -0.3 | -0.3 | 2 | 2 | 2 | 2 | -0.3 | -0.3 | -0.3 | -0.3 | -0.3 |
| 6 | 3 | 3 | -0.45 | -0.45 | -0.45 | -0.45 | -0.45 | -0.45 | 3 | 3 | -0.45 | -0.45 | -0.45 | -0.45 | -0.45 | -0.45 | -0.45 | -0.45 | -0.45 | -0.45 | 3 | 3 | -0.45 | -0.45 | -0.45 |
| 7 | 2 | 2 | -0.3 | -0.3 | -0.3 | -0.3 | -0.3 | -0.3 | 2 | 2 | -0.3 | -0.3 | -0.3 | -0.3 | -0.3 | -0.3 | -0.3 | -0.3 | -0.3 | -0.3 | 2 | 2 | -0.3 | -0.3 | -0.3 |
| 8 | -0.45 | -0.45 | 3 | 3 | -0.45 | -0.45 | -0.45 | -0.45 | 3 | 3 | -0.45 | -0.45 | -0.45 | -0.45 | -0.45 | -0.45 | -0.45 | -0.45 | -0.45 | -0.45 | 3 | 3 | -0.45 | -0.45 | -0.45 |
| 9 | -0.3 | -0.3 | 2 | 2 | -0.3 | -0.3 | -0.3 | -0.3 | 2 | 2 | -0.3 | -0.3 | -0.3 | -0.3 | -0.3 | -0.3 | -0.3 | -0.3 | -0.3 | -0.3 | 2 | 2 | -0.3 | -0.3 | -0.3 |
| 10 | -0.3 | -0.3 | 2 | 2 | -0.3 | -0.3 | -0.3 | -0.3 | -0.3 | -0.3 | 2 | 2 | -0.3 | -0.3 | -0.3 | -0.3 | -0.3 | -0.3 | -0.3 | -0.3 | -0.3 | -0.3 | 2 | 2 | -0.3 |
| 11 | -0.3 | -0.3 | 2 | 2 | -0.3 | -0.3 | -0.3 | -0.3 | -0.3 | -0.3 | 2 | 2 | -0.3 | -0.3 | -0.3 | -0.3 | -0.3 | -0.3 | -0.3 | -0.3 | -0.3 | -0.3 | 2 | 2 | -0.3 |
| 12 | -0.45 | -0.45 | 3 | 3 | -0.45 | -0.45 | -0.45 | -0.45 | -0.45 | -0.45 | 3 | 3 | -0.45 | -0.45 | -0.45 | -0.45 | -0.45 | -0.45 | -0.45 | -0.45 | -0.45 | -0.45 | 3 | 3 | -0.45 |
| 13 | -0.3 | -0.3 | 2 | 2 | -0.3 | -0.3 | -0.3 | -0.3 | -0.3 | -0.3 | 2 | 2 | -0.3 | -0.3 | -0.3 | -0.3 | -0.3 | -0.3 | -0.3 | -0.3 | -0.3 | -0.3 | 2 | 2 | -0.3 |
| 14 | 3 | 3 | -0.45 | -0.45 | -0.45 | -0.45 | -0.45 | -0.45 | -0.45 | -0.45 | -0.45 | -0.45 | 3 | 3 | -0.45 | -0.45 | 3 | 3 | -0.45 | -0.45 | -0.45 | -0.45 | -0.45 | -0.45 | -0.45 |
| 15 | 2 | 2 | -0.3 | -0.3 | -0.3 | -0.3 | -0.3 | -0.3 | -0.3 | -0.3 | -0.3 | -0.3 | 2 | 2 | -0.3 | -0.3 | 2 | 2 | -0.3 | -0.3 | -0.3 | -0.3 | -0.3 | -0.3 | 2 |
| 16 | 2 | 2 | -0.3 | -0.3 | -0.3 | -0.3 | -0.3 | -0.3 | -0.3 | -0.3 | -0.3 | -0.3 | 2 | 2 | -0.3 | -0.3 | 2 | 2 | -0.3 | -0.3 | -0.3 | -0.3 | -0.3 | -0.3 | 2 |
| 17 | -0.3 | -0.3 | 2 | 2 | -0.3 | -0.3 | -0.3 | -0.3 | -0.3 | -0.3 | -0.3 | -0.3 | 2 | 2 | -0.3 | -0.3 | 2 | 2 | -0.3 | -0.3 | -0.3 | -0.3 | -0.3 | -0.3 | -0.3 |
| 18 | -0.3 | -0.3 | 2 | 2 | -0.3 | -0.3 | -0.3 | -0.3 | -0.3 | -0.3 | -0.3 | -0.3 | 2 | 2 | -0.3 | -0.3 | 2 | 2 | -0.3 | -0.3 | -0.3 | -0.3 | -0.3 | -0.3 | 2 |
| 19 | -0.3 | -0.3 | 2 | 2 | -0.3 | -0.3 | -0.3 | -0.3 | -0.3 | -0.3 | -0.3 | -0.3 | 2 | 2 | -0.3 | -0.3 | 2 | 2 | -0.3 | -0.3 | -0.3 | -0.3 | -0.3 | -0.3 | 2 |
| 20 | -0.45 | -0.45 | 3 | 3 | -0.45 | -0.45 | -0.45 | -0.45 | -0.45 | -0.45 | -0.45 | -0.45 | 3 | 3 | -0.45 | -0.45 | 3 | 3 | -0.45 | -0.45 | -0.45 | -0.45 | -0.45 | -0.45 | -0.45 |
| 21 | -0.3 | -0.3 | 2 | 2 | -0.3 | -0.3 | -0.3 | -0.3 | -0.3 | -0.3 | -0.3 | -0.3 | 2 | 2 | -0.3 | -0.3 | 2 | 2 | -0.3 | -0.3 | -0.3 | -0.3 | -0.3 | -0.3 | 2 |
| 22 | -0.3 | -0.3 | 2 | 2 | -0.3 | -0.3 | -0.3 | -0.3 | -0.3 | -0.3 | -0.3 | -0.3 | 2 | 2 | -0.3 | -0.3 | 2 | 2 | -0.3 | -0.3 | -0.3 | -0.3 | -0.3 | -0.3 | 2 |

**B2.8 Table: Transition node → PFC weight matrix**

| | 26 | 27 | 28 | 29 | 30 | 31 | 32 | 33 | 34 | 35 | 36 | 37 | 38 | 39 | 40 | 41 | 42 | 43 | 44 | 45 | 46 | 47 | 48 | 49 |
|---|---|---|---|---|---|---|---|---|---|---|---|---|---|---|---|---|---|---|---|---|---|---|---|---|
| 1 | -0.3 | -0.3 | -0.3 | 2 | 2 | -0.3 | -0.3 | -0.3 | -0.3 | -0.3 | -0.3 | -0.3 | -0.3 | -0.3 | -0.3 | -0.3 | -0.3 | -0.3 | -0.3 | -0.3 | -0.3 | -0.3 | -0.3 | -0.3 |
| 2 | -0.3 | 2 | 2 | -0.3 | -0.3 | 2 | 2 | -0.3 | -0.3 | -0.3 | -0.3 | -0.3 | -0.3 | -0.3 | -0.3 | -0.3 | -0.3 | -0.3 | -0.3 | -0.3 | -0.3 | -0.3 | -0.3 | -0.3 |
| 3 | -0.3 | -0.3 | -0.3 | 2 | 2 | -0.3 | -0.3 | -0.3 | -0.3 | -0.3 | -0.3 | -0.3 | -0.3 | -0.3 | -0.3 | -0.3 | -0.3 | -0.3 | -0.3 | -0.3 | -0.3 | -0.3 | -0.3 | -0.3 |
| 4 | -0.3 | -0.3 | -0.3 | -0.3 | -0.3 | -0.3 | -0.3 | -0.3 | -0.3 | 2 | 2 | -0.3 | -0.3 | -0.3 | -0.3 | -0.3 | -0.3 | -0.3 | -0.3 | -0.3 | -0.3 | -0.3 | -0.3 | -0.3 |
| 5 | -0.3 | 2 | 2 | -0.3 | -0.3 | -0.3 | -0.3 | 2 | 2 | -0.3 | -0.3 | -0.3 | -0.3 | -0.3 | -0.3 | -0.3 | -0.3 | -0.3 | -0.3 | -0.3 | -0.3 | -0.3 | -0.3 | -0.3 |
| 6 | -0.45 | -0.45 | -0.45 | 3 | 3 | -0.45 | -0.45 | -0.45 | -0.45 | -0.45 | -0.45 | -0.45 | -0.45 | -0.45 | -0.45 | -0.45 | -0.45 | -0.45 | -0.45 | -0.45 | -0.45 | -0.45 | -0.45 | -0.45 |
| 7 | -0.3 | 2 | 2 | -0.3 | -0.3 | -0.3 | -0.3 | 2 | 2 | -0.3 | -0.3 | -0.3 | -0.3 | -0.3 | -0.3 | -0.3 | -0.3 | -0.3 | -0.3 | -0.3 | -0.3 | -0.3 | -0.3 | -0.3 |
| 8 | -0.45 | -0.45 | -0.45 | 3 | 3 | -0.45 | -0.45 | -0.45 | -0.45 | -0.45 | -0.45 | -0.45 | -0.45 | -0.45 | -0.45 | -0.45 | -0.45 | -0.45 | -0.45 | -0.45 | -0.45 | -0.45 | -0.45 | -0.45 |
| 9 | -0.3 | 2 | 2 | -0.3 | -0.3 | -0.3 | -0.3 | 2 | 2 | -0.3 | -0.3 | -0.3 | -0.3 | -0.3 | -0.3 | -0.3 | -0.3 | -0.3 | -0.3 | -0.3 | -0.3 | -0.3 | -0.3 | -0.3 |
| 10 | -0.3 | -0.3 | -0.3 | 2 | 2 | -0.3 | -0.3 | -0.3 | -0.3 | -0.3 | -0.3 | 2 | 2 | -0.3 | -0.3 | -0.3 | -0.3 | -0.3 | -0.3 | -0.3 | -0.3 | -0.3 | -0.3 | -0.3 |
| 11 | -0.3 | 2 | 2 | -0.3 | -0.3 | -0.3 | -0.3 | 2 | 2 | -0.3 | -0.3 | 2 | 2 | -0.3 | -0.3 | -0.3 | -0.3 | -0.3 | -0.3 | -0.3 | -0.3 | -0.3 | -0.3 | -0.3 |
| 12 | -0.45 | -0.45 | -0.45 | 3 | 3 | -0.45 | -0.45 | -0.45 | -0.45 | -0.45 | -0.45 | -0.45 | -0.45 | -0.45 | -0.45 | 3 | 3 | -0.45 | -0.45 | -0.45 | -0.45 | -0.45 | -0.45 | -0.45 |
| 13 | -0.3 | 2 | 2 | -0.3 | -0.3 | -0.3 | -0.3 | 2 | 2 | -0.3 | -0.3 | -0.3 | -0.3 | -0.3 | -0.3 | 2 | 2 | -0.3 | -0.3 | -0.3 | -0.3 | -0.3 | -0.3 | -0.3 |
| 14 | -0.45 | -0.45 | -0.45 | 3 | 3 | -0.45 | -0.45 | -0.45 | -0.45 | -0.45 | -0.45 | -0.45 | -0.45 | -0.45 | -0.45 | -0.45 | -0.45 | -0.45 | -0.45 | -0.45 | -0.45 | -0.45 | -0.45 | -0.45 |
| 15 | 2 | -0.3 | -0.3 | -0.3 | -0.3 | -0.3 | -0.3 | -0.3 | -0.3 | 2 | 2 | -0.3 | -0.3 | -0.3 | -0.3 | -0.3 | -0.3 | -0.3 | -0.3 | -0.3 | -0.3 | -0.3 | -0.3 | -0.3 |
| 16 | 2 | 2 | 2 | -0.3 | -0.3 | -0.3 | -0.3 | 2 | 2 | -0.3 | -0.3 | -0.3 | -0.3 | -0.3 | -0.3 | -0.3 | -0.3 | -0.3 | -0.3 | -0.3 | -0.3 | -0.3 | -0.3 | -0.3 |
| 17 | -0.3 | -0.3 | -0.3 | 2 | 2 | -0.3 | -0.3 | -0.3 | -0.3 | -0.3 | -0.3 | -0.3 | -0.3 | -0.3 | -0.3 | -0.3 | -0.3 | 2 | 2 | -0.3 | -0.3 | -0.3 | -0.3 | -0.3 |
| 18 | 2 | -0.3 | -0.3 | -0.3 | -0.3 | -0.3 | -0.3 | -0.3 | -0.3 | 2 | 2 | -0.3 | -0.3 | -0.3 | -0.3 | -0.3 | -0.3 | 2 | 2 | -0.3 | -0.3 | -0.3 | -0.3 | -0.3 |
| 19 | 2 | 2 | 2 | -0.3 | -0.3 | -0.3 | -0.3 | 2 | 2 | -0.3 | -0.3 | -0.3 | -0.3 | -0.3 | -0.3 | -0.3 | -0.3 | 2 | 2 | -0.3 | -0.3 | -0.3 | -0.3 | -0.3 |
| 20 | -0.45 | -0.45 | -0.45 | 3 | 3 | -0.45 | -0.45 | -0.45 | -0.45 | -0.45 | -0.45 | -0.45 | -0.45 | 3 | 3 | -0.45 | -0.45 | -0.45 | -0.45 | -0.45 | -0.45 | -0.45 | -0.45 | -0.45 |
| 21 | 2 | -0.3 | -0.3 | -0.3 | -0.3 | -0.3 | -0.3 | -0.3 | -0.3 | 2 | 2 | -0.3 | -0.3 | 2 | 2 | -0.3 | -0.3 | -0.3 | -0.3 | -0.3 | -0.3 | -0.3 | -0.3 | -0.3 |
| 22 | 2 | 2 | 2 | -0.3 | -0.3 | -0.3 | -0.3 | 2 | 2 | -0.3 | -0.3 | -0.3 | -0.3 | 2 | 2 | -0.3 | -0.3 | -0.3 | -0.3 | -0.3 | -0.3 | -0.3 | -0.3 | -0.3 |

**B2.9 Table Environment node → transition node weight matrix**

Rows = Source environment nodes ; Columns = Target transition nodes. Continued on next page.

| | 1 | 2 | 3 | 4 | 5 | 6 | 7 | 8 | 9 | 10 | 11 | 12 | 13 | 14 | 15 | 16 | 17 | 18 | 19 | 20 | 21 | 22 |
|---|---|---|---|---|---|---|---|---|---|---|---|---|---|---|---|---|---|---|---|---|---|---|
| 1 | -0.1 | 0.083 | -0.1 | -0.1 | -0.1 | -0.1 | -0.1 | -0.1 | -0.1 | -0.1 | -0.1 | -0.1 | -0.1 | -0.1 | -0.1 | -0.1 | -0.1 | -0.1 | -0.1 | -0.1 | -0.1 | -0.1 |
| 2 | -0.1 | -0.1 | -0.1 | 0.083 | 0.077 | -0.1 | -0.1 | -0.1 | -0.1 | -0.1 | -0.1 | -0.1 | -0.1 | -0.1 | 0.083 | 0.071 | -0.1 | 0.077 | 0.0667 | -0.1 | 0.083 | 0.071 |
| 3 | -0.1 | -0.1 | -0.1 | -0.1 | -0.1 | -0.1 | -0.1 | -0.1 | -0.1 | -0.1 | -0.1 | -0.1 | -0.1 | -0.1 | -0.1 | -0.1 | -0.1 | -0.1 | -0.1 | -0.1 | -0.1 | -0.1 |
| 4 | -0.1 | -0.1 | -0.1 | -0.1 | -0.1 | -0.1 | 0.077 | -0.1 | 0.071 | -0.1 | -0.1 | -0.1 | -0.1 | -0.1 | -0.1 | -0.1 | -0.1 | -0.1 | -0.1 | -0.1 | -0.1 | -0.1 |
| 5 | -0.1 | -0.1 | -0.1 | -0.1 | -0.1 | -0.1 | -0.1 | -0.1 | -0.1 | -0.1 | 0.077 | -0.1 | 0.077 | -0.1 | -0.1 | -0.1 | -0.1 | -0.1 | -0.1 | -0.1 | -0.1 | -0.1 |
| 6 | -0.1 | -0.1 | -0.1 | -0.1 | -0.1 | -0.1 | -0.1 | -0.1 | -0.1 | -0.1 | -0.1 | -0.1 | -0.1 | -0.1 | -0.1 | -0.1 | -0.1 | -0.1 | -0.1 | -0.1 | -0.1 | -0.1 |
| 7 | -0.1 | -0.1 | -0.1 | -0.1 | -0.1 | -0.1 | -0.1 | -0.1 | -0.1 | -0.1 | -0.1 | -0.1 | -0.1 | -0.1 | -0.1 | -0.1 | -0.1 | -0.1 | -0.1 | -0.1 | -0.1 | -0.1 |
| 8 | -0.1 | -0.1 | -0.1 | -0.1 | -0.1 | -0.1 | -0.1 | -0.1 | -0.1 | -0.1 | -0.1 | -0.1 | -0.1 | -0.1 | -0.1 | -0.1 | -0.1 | -0.1 | -0.1 | -0.1 | -0.1 | -0.1 |
| 9 | -0.1 | -0.1 | -0.1 | -0.1 | -0.1 | -0.1 | -0.1 | -0.1 | -0.1 | -0.1 | -0.1 | -0.1 | -0.1 | -0.1 | -0.1 | -0.1 | -0.1 | -0.1 | -0.1 | -0.1 | -0.1 | -0.1 |
| 10 | -0.1 | -0.1 | -0.1 | -0.1 | -0.1 | -0.1 | -0.1 | -0.1 | -0.1 | -0.1 | -0.1 | -0.1 | -0.1 | -0.1 | -0.1 | -0.1 | -0.1 | -0.1 | -0.1 | -0.1 | -0.1 | -0.1 |
| 11 | -0.1 | -0.1 | -0.1 | -0.1 | -0.1 | -0.1 | -0.1 | -0.1 | -0.1 | -0.1 | -0.1 | -0.1 | -0.1 | -0.1 | -0.1 | -0.1 | -0.1 | -0.1 | -0.1 | -0.1 | -0.1 | -0.1 |
| 12 | -0.1 | -0.1 | -0.1 | -0.1 | -0.1 | -0.1 | -0.1 | -0.1 | -0.1 | -0.1 | -0.1 | -0.1 | -0.1 | -0.1 | -0.1 | -0.1 | -0.1 | -0.1 | -0.1 | -0.1 | -0.1 | -0.1 |
| 13 | -0.1 | -0.1 | -0.1 | -0.1 | -0.1 | -0.1 | -0.1 | -0.1 | -0.1 | -0.1 | -0.1 | -0.1 | -0.1 | -0.1 | -0.1 | -0.1 | -0.1 | -0.1 | -0.1 | -0.1 | -0.1 | -0.1 |
| 14 | 0.045 | -0.05 | 0.045 | 0.042 | 0.038 | -0.05 | -0.05 | 0.038 | 0.036 | 0.042 | 0.038 | 0.042 | 0.038 | -0.05 | -0.05 | -0.05 | 0.042 | 0.038 | 0.033 | 0.045 | 0.042 | 0.036 |
| 15 | 0.0455 | 0.042 | 0.045 | -0.05 | -0.05 | 0.042 | 0.038 | 0.038 | 0.036 | 0.042 | 0.038 | 0.042 | 0.038 | 0.045 | -0.05 | -0.05 | 0.042 | -0.05 | -0.05 | 0.045 | -0.05 | -0.05 |
| 16 | 0.045 | 0.042 | 0.045 | 0.042 | -0.05 | 0.042 | 0.038 | 0.038 | 0.036 | 0.042 | 0.038 | 0.042 | 0.038 | 0.045 | 0.042 | 0.036 | 0.042 | 0.038 | 0.033 | 0.045 | 0.042 | 0.036 |
| 17 | 0.045 | 0.042 | 0.045 | 0.042 | 0.038 | 0.042 | -0.05 | 0.038 | -0.05 | 0.042 | 0.038 | 0.042 | 0.038 | 0.045 | 0.042 | 0.036 | 0.042 | 0.038 | 0.033 | 0.045 | 0.042 | 0.036 |
| 18 | 0.045 | 0.042 | 0.045 | 0.042 | 0.038 | 0.042 | 0.038 | 0.038 | 0.036 | 0.042 | -0.05 | 0.042 | -0.05 | 0.045 | 0.042 | 0.036 | 0.042 | 0.038 | 0.033 | 0.045 | 0.042 | 0.036 |
| 19 | 0.045 | 0.042 | 0.045 | 0.042 | 0.038 | 0.042 | 0.038 | 0.038 | 0.036 | 0.042 | 0.038 | 0.042 | 0.038 | 0.045 | 0.042 | 0.036 | 0.042 | 0.038 | 0.033 | 0.045 | 0.042 | 0.036 |
| 20 | 0.045 | 0.042 | 0.045 | 0.042 | 0.038 | 0.042 | 0.038 | 0.038 | 0.036 | 0.042 | 0.038 | 0.042 | 0.038 | 0.045 | 0.042 | 0.036 | 0.042 | 0.038 | 0.033 | 0.045 | 0.042 | 0.036 |
| 21 | -0.05 | -0.05 | -0.05 | -0.05 | -0.05 | -0.05 | -0.05 | -0.05 | -0.05 | -0.05 | -0.05 | -0.05 | -0.05 | -0.05 | -0.05 | -0.05 | -0.05 | -0.05 | -0.05 | -0.05 | -0.05 | -0.05 |
| 22 | -0.05 | -0.05 | -0.05 | -0.05 | -0.05 | -0.05 | -0.05 | -0.05 | -0.05 | -0.05 | -0.05 | -0.05 | -0.05 | -0.05 | -0.05 | -0.05 | -0.05 | -0.05 | -0.05 | -0.05 | -0.05 | -0.05 |
| 23 | -0.05 | -0.05 | -0.05 | -0.05 | -0.05 | -0.05 | -0.05 | -0.05 | -0.05 | -0.05 | -0.05 | -0.05 | -0.05 | -0.05 | -0.05 | -0.05 | -0.05 | -0.05 | -0.05 | -0.05 | -0.05 | -0.05 |
| 24 | -0.05 | -0.05 | -0.05 | -0.05 | -0.05 | -0.05 | -0.05 | -0.05 | -0.05 | -0.05 | -0.05 | -0.05 | -0.05 | -0.05 | -0.05 | -0.05 | -0.05 | -0.05 | -0.05 | -0.05 | -0.05 | -0.05 |
| 25 | -0.05 | -0.05 | -0.05 | -0.05 | -0.05 | -0.05 | -0.05 | -0.05 | -0.05 | -0.05 | -0.05 | -0.05 | -0.05 | -0.05 | -0.05 | -0.05 | -0.05 | -0.05 | -0.05 | -0.05 | -0.05 | -0.05 |
| 26 | -0.05 | -0.05 | -0.05 | -0.05 | -0.05 | -0.05 | -0.05 | -0.05 | -0.05 | -0.05 | -0.05 | -0.05 | -0.05 | -0.05 | -0.05 | -0.05 | -0.05 | -0.05 | -0.05 | -0.05 | -0.05 | -0.05 |
| 27 | -0.1 | -0.1 | -0.1 | -0.1 | -0.1 | -0.1 | -0.1 | -0.1 | -0.1 | -0.1 | -0.1 | -0.1 | -0.1 | -0.1 | -0.1 | -0.1 | -0.1 | -0.1 | -0.1 | -0.1 | -0.1 | -0.1 |
| 28 | -0.1 | -0.1 | -0.1 | -0.1 | -0.1 | -0.1 | -0.1 | -0.1 | -0.1 | -0.1 | -0.1 | -0.1 | -0.1 | -0.1 | -0.1 | -0.1 | -0.1 | -0.1 | -0.1 | -0.1 | -0.1 | -0.1 |
| 29 | -0.1 | -0.1 | -0.1 | -0.1 | -0.1 | -0.1 | -0.1 | -0.1 | -0.1 | -0.1 | -0.1 | -0.1 | -0.1 | -0.1 | -0.1 | -0.1 | -0.1 | -0.1 | -0.1 | -0.1 | -0.1 | -0.1 |
| 30 | -0.1 | -0.1 | -0.1 | -0.1 | -0.1 | -0.1 | -0.1 | -0.1 | -0.1 | -0.1 | -0.1 | -0.1 | -0.1 | -0.1 | -0.1 | -0.1 | -0.1 | -0.1 | -0.1 | -0.1 | -0.1 | -0.1 |
| 31 | -0.1 | -0.1 | -0.1 | -0.1 | -0.1 | -0.1 | -0.1 | -0.1 | -0.1 | -0.1 | -0.1 | -0.1 | -0.1 | -0.1 | -0.1 | -0.1 | -0.1 | -0.1 | -0.1 | -0.1 | -0.1 | -0.1 |
| 32 | -0.1 | -0.1 | -0.1 | -0.1 | -0.1 | -0.1 | -0.1 | -0.1 | -0.1 | -0.1 | -0.1 | -0.1 | -0.1 | -0.1 | -0.1 | -0.1 | -0.1 | -0.1 | -0.1 | -0.1 | -0.1 | -0.1 |
| 33 | -0.1 | -0.1 | -0.1 | -0.1 | -0.1 | -0.1 | -0.1 | -0.1 | -0.1 | -0.1 | -0.1 | -0.1 | -0.1 | -0.1 | -0.1 | -0.1 | -0.1 | -0.1 | -0.1 | -0.1 | -0.1 | -0.1 |
| 34 | -0.1 | -0.1 | -0.1 | -0.1 | 0.077 | -0.1 | -0.1 | -0.1 | -0.1 | -0.1 | -0.1 | -0.1 | -0.1 | -0.1 | -0.1 | -0.1 | -0.1 | -0.1 | -0.1 | -0.1 | -0.1 | -0.1 |
| 35 | -0.1 | -0.1 | -0.1 | -0.1 | -0.1 | -0.1 | -0.1 | -0.1 | -0.1 | -0.1 | -0.1 | -0.1 | -0.1 | -0.1 | -0.1 | -0.1 | -0.1 | -0.1 | -0.1 | -0.1 | -0.1 | -0.1 |
| 36 | -0.1 | -0.1 | -0.1 | -0.1 | -0.1 | -0.1 | -0.1 | -0.1 | -0.1 | -0.1 | -0.1 | -0.1 | -0.1 | -0.1 | -0.1 | -0.1 | -0.1 | -0.1 | -0.1 | -0.1 | -0.1 | -0.1 |
| 37 | -0.1 | -0.1 | -0.1 | -0.1 | -0.1 | -0.1 | -0.1 | -0.1 | -0.1 | -0.1 | -0.1 | -0.1 | -0.1 | -0.1 | -0.1 | 0.071 | -0.1 | -0.1 | 0.067 | -0.1 | -0.1 | 0.071 |
| 38 | -0.1 | -0.1 | -0.1 | -0.1 | -0.1 | -0.1 | -0.1 | -0.1 | -0.1 | -0.1 | -0.1 | -0.1 | -0.1 | -0.1 | -0.1 | -0.1 | -0.1 | -0.1 | -0.1 | -0.1 | -0.1 | -0.1 |
| 39 | -0.1 | -0.1 | -0.1 | -0.1 | -0.1 | -0.1 | -0.1 | -0.1 | -0.1 | -0.1 | -0.1 | -0.1 | -0.1 | -0.1 | -0.1 | -0.1 | -0.1 | -0.1 | -0.1 | -0.1 | -0.1 | -0.1 |
| 40 | -0.05 | -0.05 | -0.05 | -0.05 | -0.05 | -0.05 | -0.05 | -0.05 | -0.05 | -0.05 | -0.05 | -0.05 | -0.05 | -0.05 | -0.05 | -0.05 | -0.05 | -0.05 | -0.05 | -0.05 | -0.05 | -0.05 |

**B2.9 Table Environment node → transition node weight matrix**

Continued from previous page. Continued on next page.

| | 1 | 2 | 3 | 4 | 5 | 6 | 7 | 8 | 9 | 10 | 11 | 12 | 13 | 14 | 15 | 16 | 17 | 18 | 19 | 20 | 21 | 22 |
|---|---|---|---|---|---|---|---|---|---|---|---|---|---|---|---|---|---|---|---|---|---|---|
| 41 | -0.05 | -0.05 | -0.05 | -0.05 | -0.05 | -0.05 | -0.05 | -0.05 | -0.05 | -0.05 | -0.05 | -0.05 | -0.05 | -0.05 | -0.05 | -0.05 | -0.05 | -0.05 | -0.05 | -0.05 | -0.05 | -0.05 |
| 42 | -0.05 | -0.05 | -0.05 | -0.05 | -0.05 | -0.05 | -0.05 | -0.05 | -0.05 | -0.05 | -0.05 | -0.05 | -0.05 | -0.05 | -0.05 | -0.05 | -0.05 | -0.05 | -0.05 | -0.05 | -0.05 | -0.05 |
| 43 | -0.05 | -0.05 | -0.05 | -0.05 | -0.05 | -0.05 | -0.05 | -0.05 | -0.05 | -0.05 | -0.05 | -0.05 | -0.05 | -0.05 | -0.05 | -0.05 | -0.05 | -0.05 | -0.05 | -0.05 | -0.05 | -0.05 |
| 44 | -0.05 | -0.05 | -0.05 | -0.05 | -0.05 | -0.05 | -0.05 | -0.05 | -0.05 | -0.05 | -0.05 | -0.05 | -0.05 | -0.05 | -0.05 | -0.05 | -0.05 | -0.05 | -0.05 | -0.05 | -0.05 | -0.05 |
| 45 | -0.05 | -0.05 | -0.05 | -0.05 | -0.05 | -0.05 | -0.05 | -0.05 | -0.05 | -0.05 | -0.05 | -0.05 | -0.05 | -0.05 | -0.05 | -0.05 | -0.05 | -0.05 | -0.05 | -0.05 | -0.05 | -0.05 |
| 46 | -0.05 | -0.05 | -0.05 | -0.05 | -0.05 | -0.05 | -0.05 | -0.05 | -0.05 | -0.05 | -0.05 | -0.05 | -0.05 | -0.05 | -0.05 | -0.05 | -0.05 | -0.05 | -0.05 | -0.05 | -0.05 | -0.05 |
| 47 | 0.045 | 0.042 | 0.045 | 0.042 | 0.038 | 0.042 | 0.038 | 0.038 | 0.036 | 0.042 | 0.038 | 0.042 | 0.038 | 0.045 | 0.042 | 0.036 | 0.042 | 0.038 | 0.033 | 0.045 | 0.042 | 0.036 |
| 48 | -0.05 | -0.05 | -0.05 | -0.05 | -0.05 | -0.05 | -0.05 | -0.05 | -0.05 | -0.05 | -0.05 | -0.05 | -0.05 | -0.05 | -0.05 | -0.05 | -0.05 | -0.05 | -0.05 | -0.05 | -0.05 | -0.05 |
| 49 | -0.05 | -0.05 | -0.05 | -0.05 | -0.05 | -0.05 | -0.05 | -0.05 | -0.05 | -0.05 | -0.05 | -0.05 | -0.05 | -0.05 | -0.05 | -0.05 | -0.05 | -0.05 | -0.05 | -0.05 | -0.05 | -0.05 |
| 50 | -0.05 | -0.05 | -0.05 | -0.05 | -0.05 | -0.05 | -0.05 | -0.05 | -0.05 | -0.05 | -0.05 | -0.05 | -0.05 | -0.05 | -0.05 | -0.05 | -0.05 | -0.05 | -0.05 | -0.05 | -0.05 | -0.05 |
| 51 | -0.05 | -0.05 | -0.05 | -0.05 | -0.05 | -0.05 | -0.05 | -0.05 | -0.05 | -0.05 | -0.05 | -0.05 | -0.05 | -0.05 | -0.05 | -0.05 | -0.05 | -0.05 | -0.05 | -0.05 | -0.05 | -0.05 |
| 52 | -0.05 | -0.05 | -0.05 | -0.05 | -0.05 | -0.05 | -0.05 | -0.05 | -0.05 | -0.05 | -0.05 | -0.05 | -0.05 | -0.05 | -0.05 | -0.05 | -0.05 | -0.05 | -0.05 | -0.05 | -0.05 | -0.05 |
| 53 | -0.05 | -0.05 | -0.05 | -0.05 | -0.05 | -0.05 | -0.05 | -0.05 | -0.05 | -0.05 | -0.05 | -0.05 | -0.05 | -0.05 | -0.05 | -0.05 | -0.05 | -0.05 | -0.05 | -0.05 | -0.05 | -0.05 |
| 54 | -0.05 | -0.05 | -0.05 | -0.05 | -0.05 | -0.05 | -0.05 | -0.05 | -0.05 | -0.05 | -0.05 | -0.05 | -0.05 | -0.05 | -0.05 | -0.05 | -0.05 | -0.05 | -0.05 | -0.05 | -0.05 | -0.05 |
| 55 | -0.05 | -0.05 | -0.05 | -0.05 | -0.05 | -0.05 | -0.05 | -0.05 | -0.05 | -0.05 | -0.05 | -0.05 | -0.05 | -0.05 | -0.05 | -0.05 | -0.05 | -0.05 | -0.05 | -0.05 | -0.05 | -0.05 |
| 56 | -0.05 | -0.05 | -0.05 | -0.05 | -0.05 | -0.05 | -0.05 | -0.05 | -0.05 | -0.05 | -0.05 | -0.05 | -0.05 | -0.05 | -0.05 | -0.05 | -0.05 | -0.05 | -0.05 | -0.05 | -0.05 | -0.05 |
| 57 | -0.05 | -0.05 | -0.05 | -0.05 | -0.05 | -0.05 | -0.05 | -0.05 | -0.05 | -0.05 | -0.05 | -0.05 | -0.05 | -0.05 | -0.05 | -0.05 | -0.05 | -0.05 | -0.05 | -0.05 | -0.05 | -0.05 |
| 58 | -0.05 | -0.05 | -0.05 | -0.05 | -0.05 | -0.05 | -0.05 | -0.05 | -0.05 | -0.05 | -0.05 | -0.05 | -0.05 | -0.05 | -0.05 | -0.05 | -0.05 | -0.05 | -0.05 | -0.05 | -0.05 | -0.05 |
| 59 | -0.05 | -0.05 | -0.05 | -0.05 | -0.05 | -0.05 | -0.05 | -0.05 | -0.05 | -0.05 | -0.05 | -0.05 | -0.05 | -0.05 | -0.05 | -0.05 | -0.05 | -0.05 | -0.05 | -0.05 | -0.05 | -0.05 |
| 50 | -0.05 | -0.05 | -0.05 | -0.05 | -0.05 | -0.05 | -0.05 | -0.05 | -0.05 | -0.05 | -0.05 | -0.05 | -0.05 | -0.05 | -0.05 | -0.05 | -0.05 | -0.05 | -0.05 | -0.05 | -0.05 | -0.05 |
| 61 | 0.045 | 0.042 | 0.045 | 0.042 | 0.038 | 0.042 | 0.038 | 0.038 | 0.036 | -0.05 | -0.05 | -0.05 | -0.05 | -0.05 | -0.05 | -0.05 | -0.05 | -0.05 | -0.05 | -0.05 | -0.05 | -0.05 |
| 62 | -0.05 | -0.05 | -0.05 | -0.05 | -0.05 | -0.05 | -0.05 | -0.05 | -0.05 | -0.05 | -0.05 | -0.05 | -0.05 | -0.05 | -0.05 | -0.05 | -0.05 | -0.05 | -0.05 | -0.05 | -0.05 | -0.05 |
| 63 | -0.05 | -0.05 | -0.05 | -0.05 | -0.05 | -0.05 | -0.05 | -0.05 | -0.05 | -0.05 | -0.05 | -0.05 | -0.05 | -0.05 | -0.05 | -0.05 | -0.05 | -0.05 | -0.05 | -0.05 | -0.05 | -0.05 |
| 64 | -0.05 | -0.05 | -0.05 | -0.05 | -0.05 | -0.05 | -0.05 | -0.05 | -0.05 | -0.05 | -0.05 | -0.05 | -0.05 | -0.05 | -0.05 | -0.05 | -0.05 | -0.05 | -0.05 | -0.05 | -0.05 | -0.05 |
| 65 | -0.05 | -0.05 | -0.05 | -0.05 | -0.05 | -0.05 | -0.05 | -0.05 | -0.05 | -0.05 | -0.05 | -0.05 | -0.05 | -0.05 | -0.05 | -0.05 | -0.05 | -0.05 | -0.05 | -0.05 | -0.05 | -0.05 |
| 66 | -0.05 | -0.05 | -0.05 | -0.05 | -0.05 | -0.05 | -0.05 | -0.05 | -0.05 | -0.05 | -0.05 | -0.05 | -0.05 | -0.05 | -0.05 | -0.05 | -0.05 | -0.05 | -0.05 | -0.05 | -0.05 | -0.05 |
| 67 | -0.05 | -0.05 | -0.05 | -0.05 | -0.05 | -0.05 | -0.05 | -0.05 | -0.05 | -0.05 | -0.05 | -0.05 | -0.05 | -0.05 | -0.05 | -0.05 | -0.05 | -0.05 | -0.05 | -0.05 | -0.05 | -0.05 |
| 68 | -0.05 | -0.05 | -0.05 | -0.05 | -0.05 | -0.05 | -0.05 | -0.05 | -0.05 | -0.05 | -0.05 | -0.05 | -0.05 | -0.05 | -0.05 | -0.05 | -0.05 | -0.05 | -0.05 | -0.05 | -0.05 | -0.05 |
| 69 | -0.05 | -0.05 | -0.05 | -0.05 | -0.05 | -0.05 | -0.05 | -0.05 | -0.05 | -0.05 | -0.05 | -0.05 | -0.05 | -0.05 | -0.05 | -0.05 | -0.05 | -0.05 | -0.05 | -0.05 | -0.05 | -0.05 |
| 70 | -0.05 | -0.05 | -0.05 | -0.05 | -0.05 | -0.05 | -0.05 | -0.05 | -0.05 | -0.05 | -0.05 | -0.05 | -0.05 | -0.05 | -0.05 | -0.05 | -0.05 | -0.05 | -0.05 | -0.05 | -0.05 | -0.05 |
| 71 | -0.05 | -0.05 | -0.05 | -0.05 | -0.05 | -0.05 | -0.05 | -0.05 | -0.05 | -0.05 | -0.05 | -0.05 | -0.05 | -0.05 | -0.05 | -0.05 | -0.05 | -0.05 | -0.05 | -0.05 | -0.05 | -0.05 |
| 72 | -0.05 | -0.05 | -0.05 | -0.05 | -0.05 | -0.05 | -0.05 | -0.05 | -0.05 | -0.05 | -0.05 | -0.05 | -0.05 | -0.05 | -0.05 | -0.05 | -0.05 | -0.05 | -0.05 | -0.05 | -0.05 | -0.05 |
| 73 | -0.05 | -0.05 | -0.05 | -0.05 | -0.05 | -0.05 | -0.05 | -0.05 | -0.05 | -0.05 | -0.05 | -0.05 | -0.05 | -0.05 | -0.05 | -0.05 | -0.05 | -0.05 | -0.05 | -0.05 | -0.05 | -0.05 |
| 74 | -0.05 | -0.05 | -0.05 | -0.05 | -0.05 | -0.05 | -0.05 | -0.05 | -0.05 | -0.05 | -0.05 | -0.05 | -0.05 | -0.05 | -0.05 | -0.05 | -0.05 | -0.05 | -0.05 | -0.05 | -0.05 | -0.05 |
| 75 | 0.045 | 0.042 | 0.045 | 0.042 | 0.038 | 0.042 | 0.038 | 0.038 | 0.036 | 0.042 | 0.038 | 0.042 | 0.038 | 0.045 | 0.042 | 0.036 | 0.042 | 0.038 | 0.033 | -0.05 | -0.05 | -0.05 |
| 76 | -0.05 | -0.05 | -0.05 | -0.05 | -0.05 | -0.05 | -0.05 | -0.05 | -0.05 | -0.05 | -0.05 | -0.05 | -0.05 | -0.05 | -0.05 | -0.05 | -0.05 | -0.05 | -0.05 | -0.05 | -0.05 | -0.05 |
| 77 | -0.05 | -0.05 | -0.05 | -0.05 | -0.05 | -0.05 | -0.05 | -0.05 | -0.05 | -0.05 | -0.05 | -0.05 | -0.05 | -0.05 | -0.05 | -0.05 | -0.05 | -0.05 | -0.05 | -0.05 | -0.05 | -0.05 |
| 78 | -0.05 | -0.05 | -0.05 | -0.05 | -0.05 | -0.05 | -0.05 | -0.05 | -0.05 | -0.05 | -0.05 | -0.05 | -0.05 | -0.05 | -0.05 | -0.05 | -0.05 | -0.05 | -0.05 | -0.05 | -0.05 | -0.05 |
| 79 | -0.05 | -0.05 | -0.05 | -0.05 | -0.05 | -0.05 | -0.05 | -0.05 | -0.05 | -0.05 | -0.05 | -0.05 | -0.05 | -0.05 | -0.05 | -0.05 | -0.05 | -0.05 | -0.05 | -0.05 | -0.05 | -0.05 |
| 80 | -0.05 | -0.05 | -0.05 | -0.05 | -0.05 | -0.05 | -0.05 | -0.05 | -0.05 | -0.05 | -0.05 | -0.05 | -0.05 | -0.05 | -0.05 | -0.05 | -0.05 | -0.05 | -0.05 | -0.05 | -0.05 | -0.05 |

**B2.9 Table Environment node → transition node weight matrix**

Continued from previous page.

| | 1 | 2 | 3 | 4 | 5 | 6 | 7 | 8 | 9 | 10 | 11 | 12 | 13 | 14 | 15 | 16 | 17 | 18 | 19 | 20 | 21 | 22 |
|---|---|---|---|---|---|---|---|---|---|---|---|---|---|---|---|---|---|---|---|---|---|---|
| 81 | -0.05 | -0.05 | -0.05 | -0.05 | -0.05 | -0.05 | -0.05 | -0.05 | -0.05 | -0.05 | -0.05 | -0.05 | -0.05 | -0.05 | -0.05 | -0.05 | -0.05 | -0.05 | -0.05 | -0.05 | -0.05 | -0.05 |
| 82 | -0.05 | -0.05 | -0.05 | -0.05 | -0.05 | -0.05 | -0.05 | -0.05 | -0.05 | -0.05 | -0.05 | -0.05 | -0.05 | -0.05 | -0.05 | -0.05 | -0.05 | -0.05 | -0.05 | -0.05 | -0.05 | -0.05 |
| 83 | -0.05 | -0.05 | -0.05 | -0.05 | -0.05 | -0.05 | -0.05 | -0.05 | -0.05 | -0.05 | -0.05 | -0.05 | -0.05 | -0.05 | -0.05 | -0.05 | -0.05 | -0.05 | -0.05 | -0.05 | -0.05 | -0.05 |
| 84 | -0.05 | -0.05 | -0.05 | -0.05 | -0.05 | -0.05 | -0.05 | -0.05 | -0.05 | -0.05 | -0.05 | -0.05 | -0.05 | -0.05 | -0.05 | -0.05 | -0.05 | -0.05 | -0.05 | -0.05 | -0.05 | -0.05 |
| 85 | -0.05 | -0.05 | -0.05 | -0.05 | -0.05 | -0.05 | -0.05 | -0.05 | -0.05 | -0.05 | -0.05 | -0.05 | -0.05 | -0.05 | -0.05 | -0.05 | -0.05 | -0.05 | -0.05 | -0.05 | -0.05 | -0.05 |
| 86 | -0.05 | -0.05 | -0.05 | -0.05 | -0.05 | -0.05 | -0.05 | -0.05 | -0.05 | -0.05 | -0.05 | -0.05 | -0.05 | -0.05 | -0.05 | -0.05 | -0.05 | -0.05 | -0.05 | -0.05 | -0.05 | -0.05 |
| 87 | -0.05 | -0.05 | -0.05 | -0.05 | -0.05 | -0.05 | -0.05 | -0.05 | -0.05 | -0.05 | -0.05 | -0.05 | -0.05 | -0.05 | -0.05 | -0.05 | -0.05 | -0.05 | -0.05 | -0.05 | -0.05 | -0.05 |
| 88 | -0.05 | -0.05 | -0.05 | -0.05 | -0.05 | -0.05 | -0.05 | -0.05 | -0.05 | -0.05 | -0.05 | -0.05 | -0.05 | -0.05 | -0.05 | -0.05 | -0.05 | -0.05 | -0.05 | -0.05 | -0.05 | -0.05 |
| 89 | 0.045 | 0.042 | 0.045 | 0.042 | 0.038 | 0.042 | 0.038 | 0.038 | 0.036 | 0.042 | 0.038 | 0.042 | 0.038 | 0.045 | 0.042 | 0.036 | 0.042 | 0.038 | 0.033 | 0.045 | 0.042 | 0.036 |
| 90 | -0.05 | -0.05 | -0.05 | -0.05 | -0.05 | -0.05 | -0.05 | -0.05 | -0.05 | -0.05 | -0.05 | -0.05 | -0.05 | -0.05 | -0.05 | -0.05 | -0.05 | -0.05 | -0.05 | -0.05 | -0.05 | -0.05 |
| 91 | -0.05 | -0.05 | -0.05 | -0.05 | -0.05 | -0.05 | -0.05 | -0.05 | -0.05 | -0.05 | -0.05 | -0.05 | -0.05 | -0.05 | -0.05 | -0.05 | -0.05 | -0.05 | -0.05 | -0.05 | -0.05 | -0.05 |
| 92 | -0.1 | -0.1 | -0.1 | -0.1 | -0.1 | 0.083 | 0.077 | -0.1 | -0.1 | -0.1 | -0.1 | -0.1 | -0.1 | -0.1 | -0.1 | -0.1 | -0.1 | -0.1 | -0.1 | -0.1 | -0.1 | -0.1 |
| 93 | -0.1 | -0.1 | -0.1 | -0.1 | -0.1 | -0.1 | -0.1 | -0.1 | -0.1 | -0.1 | -0.1 | -0.1 | -0.1 | -0.1 | -0.1 | -0.1 | -0.1 | -0.1 | -0.1 | -0.1 | -0.1 | -0.1 |
| 94 | -0.1 | -0.1 | -0.1 | -0.1 | -0.1 | -0.1 | -0.1 | -0.1 | -0.1 | -0.1 | -0.1 | -0.1 | -0.1 | -0.1 | -0.1 | -0.1 | -0.1 | -0.1 | -0.1 | -0.1 | -0.1 | -0.1 |
| 95 | -0.1 | -0.1 | -0.1 | -0.1 | -0.1 | -0.1 | -0.1 | -0.1 | -0.1 | -0.1 | -0.1 | -0.1 | -0.1 | -0.1 | -0.1 | -0.1 | -0.1 | -0.1 | -0.1 | -0.1 | -0.1 | -0.1 |
| 96 | -0.1 | -0.1 | -0.1 | -0.1 | -0.1 | -0.1 | -0.1 | -0.1 | -0.1 | -0.1 | -0.1 | -0.1 | -0.1 | -0.1 | -0.1 | -0.1 | -0.1 | -0.1 | -0.1 | -0.1 | -0.1 | -0.1 |
| 97 | -0.1 | -0.1 | -0.1 | -0.1 | -0.1 | -0.1 | -0.1 | -0.1 | -0.1 | -0.1 | -0.1 | -0.1 | -0.1 | -0.1 | -0.1 | -0.1 | -0.1 | -0.1 | -0.1 | -0.1 | -0.1 | -0.1 |
| 98 | -0.1 | -0.1 | -0.1 | -0.1 | -0.1 | -0.1 | -0.1 | -0.1 | -0.1 | -0.1 | -0.1 | -0.1 | -0.1 | -0.1 | -0.1 | -0.1 | -0.1 | -0.1 | -0.1 | -0.1 | -0.1 | -0.1 |
| 99 | -0.1 | -0.1 | -0.1 | -0.1 | -0.1 | -0.1 | -0.1 | 0.077 | 0.071 | -0.1 | -0.1 | -0.1 | -0.1 | -0.1 | -0.1 | -0.1 | -0.1 | -0.1 | -0.1 | -0.1 | -0.1 | -0.1 |
| 100 | -0.1 | -0.1 | -0.1 | -0.1 | -0.1 | -0.1 | -0.1 | -0.1 | -0.1 | -0.1 | -0.1 | -0.1 | -0.1 | -0.1 | -0.1 | -0.1 | -0.1 | -0.1 | -0.1 | -0.1 | -0.1 | -0.1 |
| 101 | -0.1 | -0.1 | -0.1 | -0.1 | -0.1 | -0.1 | -0.1 | -0.1 | -0.1 | -0.1 | -0.1 | -0.1 | -0.1 | -0.1 | -0.1 | -0.1 | -0.1 | -0.1 | -0.1 | -0.1 | -0.1 | -0.1 |
| 102 | -0.1 | -0.1 | -0.1 | -0.1 | -0.1 | -0.1 | -0.1 | -0.1 | -0.1 | -0.1 | -0.1 | -0.1 | -0.1 | -0.1 | -0.1 | -0.1 | -0.1 | -0.1 | -0.1 | -0.1 | -0.1 | -0.1 |
| 103 | -0.1 | -0.1 | -0.1 | -0.1 | -0.1 | -0.1 | -0.1 | -0.1 | -0.1 | 0.083 | 0.077 | 0.083 | 0.077 | 0.091 | 0.083 | 0.071 | 0.083 | 0.077 | 0.067 | -0.1 | -0.1 | -0.1 |
| 104 | -0.1 | -0.1 | -0.1 | -0.1 | -0.1 | -0.1 | -0.1 | -0.1 | -0.1 | -0.1 | -0.1 | -0.1 | -0.1 | -0.1 | -0.1 | -0.1 | -0.1 | -0.1 | -0.1 | 0.091 | 0.083 | 0.071 |

#### B3: Model weights: Waiter scenario

This section details the model weights for the waiter scenario simulation in section 3.5. For this simulation, new representations were designed, as illustrated in figure C.3.1. To select and support these new representations, amendments were made to weights between associative thalamus and PFC, between PFC and object representations, recurrent connections within PFC, weights between PFC and the transition layer, and those between the environment representation and the transition layer. These are detailed below.

The number of associative BG channels was reduced to 3 as only three subtasks were required for this task. Additionally, the number of transition nodes was reduced to 21 to reflect fewer overall PFC representations, and the number of object representations reduced to 5 to account for fewer required objects.

All other parameter values were as detailed in sections B.1 & B.2 above.

**B3.1 Table: Thalamus → PFC weight matrix**
Rows = source thalamus channels; Columns = Target PFC nodes.
Continued below

| | 1 | 2 | 3 | 4 | 5 | 6 | 7 | 8 | 9 | 10 | 11 | 12 | 13 | 14 | 15 | 16 |
|---|---|---|---|---|---|---|---|---|---|---|---|---|---|---|---|---|
| 1 | 0.6 | 0.6 | 0.6 | 0.6 | 0.6 | 0.6 | 0.6 | 0.6 | 0 | 0 | 0.6 | 0.6 | 0 | 0 | 0.6 | 0.6 |
| 2 | 0.6 | 0.6 | 0.6 | 0.6 | 0 | 0 | 0.6 | 0.6 | 0.6 | 0.6 | 0.6 | 0.6 | 0 | 0 | 0 | 0 |
| 3 | 0.6 | 0.6 | 0 | 0 | 0 | 0 | 0.6 | 0.6 | 0 | 0 | 0.6 | 0.6 | 0.6 | 0.6 | 0 | 0 |

**B3.1 Table: Thalamus → PFC weight matrix**

| | 17 | 18 | 19 | 20 | 21 | 22 | 23 | 24 | 25 | 26 | 27 | 28 | 29 | 30 | 31 | 32 |
|---|---|---|---|---|---|---|---|---|---|---|---|---|---|---|---|---|
| 1 | 0 | 0 | 0 | 0 | 0 | 0 | 0 | 0 | 0 | 0 | 0.6 | 0.6 | 0.6 | 0.6 | 0.6 | 0.6 |
| 2 | 0 | 0 | 0.6 | 0.6 | 0.6 | 0.6 | 0 | 0 | 0 | 0 | 0.6 | 0.6 | 0.6 | 0.6 | 0 | 0 |
| 3 | 0.6 | 0.6 | 0 | 0 | 0 | 0 | 0 | 0 | 0.6 | 0.6 | 0.6 | 0.6 | 0.6 | 0.6 | 0 | 0 |

**B3.1 Table: Thalamus → PFC weight matrix**

| | 33 | 34 | 35 | 36 | 37 | 38 | 39 | 40 | 41 | 42 | 43 | 44 | 45 | 46 | 47 | 48 | 49 |
|---|---|---|---|---|---|---|---|---|---|---|---|---|---|---|---|---|---|
| 1 | 0 | 0 | 0 | 0 | 0 | 0 | 0 | 0 | 0 | 0 | 0 | 0 | 0 | 0 | 0 | 0 | 0 |
| 2 | 0.6 | 0.6 | 0 | 0 | 0 | 0 | 0 | 0 | 0 | 0 | 0 | 0 | 0 | 0 | 0 | 0 | 0 |
| 3 | 0.6 | 0.6 | 0.6 | 0.6 | 0 | 0 | 0.6 | 0.6 | 0 | 0 | 0.6 | 0.6 | 0 | 0 | 0 | 0 | 0 |

**B3.2 Table: PFC → PFC Recurrent weight matrix**

Rows = Source PFC nodes; Columns = Target PFC nodes.

Continued on next page.

| | 1 | 2 | 3 | 4 | 5 | 6 | 7 | 8 | 9 | 10 | 11 | 12 | 13 | 14 | 15 | 16 |
|---|---|---|---|---|---|---|---|---|---|---|---|---|---|---|---|---|
| 1 | 0.05 | 0.05 | 0.05 | 0.05 | 0.05 | 0.05 | 0.05 | 0.05 | 0.05 | 0.05 | 0.05 | 0.05 | 0.05 | 0.05 | 0.05 | 0.05 |
| 2 | 0.05 | 0.05 | 0.05 | 0.05 | 0.05 | 0.05 | 0.05 | 0.05 | 0.05 | 0.05 | 0.05 | 0.05 | 0.05 | 0.05 | 0.05 | 0.05 |
| 3 | 0.05 | 0.05 | 0.05 | 0.05 | 0.05 | 0.05 | -1 | -0.5 | 0.05 | 0.05 | -1 | -0.5 | -0.5 | -0.5 | 0.05 | 0.05 |
| 4 | 0.05 | 0.05 | 0.05 | 0.05 | 0.05 | 0.05 | -0.5 | -1 | 0.05 | 0.05 | -0.5 | -1 | -0.5 | -0.5 | 0.05 | 0.05 |
| 5 | 0.05 | 0.05 | 0.05 | 0.05 | 0.05 | 0.05 | 0.05 | 0.05 | -0.5 | -0.5 | 0.05 | 0.05 | -0.5 | -0.5 | 0.05 | 0.05 |
| 6 | 0.05 | 0.05 | 0.05 | 0.05 | 0.05 | 0.05 | 0.05 | 0.05 | -0.5 | -0.5 | 0.05 | 0.05 | -0.5 | -0.5 | 0.05 | 0.05 |
| 7 | 0.05 | 0.05 | -1 | -0.5 | 0.05 | 0.05 | 0.05 | 0.05 | 0.05 | 0.05 | -1 | -0.5 | 0.05 | 0.05 | 0.05 | 0.05 |
| 8 | 0.05 | 0.05 | -0.5 | -1 | 0.05 | 0.05 | 0.05 | 0.05 | 0.05 | 0.05 | -0.5 | -1 | 0.05 | 0.05 | 0.05 | 0.05 |
| 9 | 0.05 | 0.05 | 0.05 | 0.05 | -0.5 | -0.5 | 0.05 | 0.05 | 0.05 | 0.05 | 0.05 | 0.05 | -0.5 | -0.5 | -0.5 | -0.5 |
| 10 | 0.05 | 0.05 | 0.05 | 0.05 | -0.5 | -0.5 | 0.05 | 0.05 | 0.05 | 0.05 | 0.05 | 0.05 | -0.5 | -0.5 | -0.5 | -0.5 |
| 11 | 0.05 | 0.05 | -1 | -0.5 | 0.05 | 0.05 | -1 | -0.5 | 0.05 | 0.05 | 0.05 | 0.05 | 0.05 | 0.05 | 0.05 | 0.05 |
| 12 | 0.05 | 0.05 | -0.5 | -1 | 0.05 | 0.05 | -0.5 | -1 | 0.05 | 0.05 | 0.05 | 0.05 | 0.05 | 0.05 | 0.05 | 0.05 |
| 13 | 0.05 | 0.05 | -0.5 | -0.5 | -0.5 | -0.5 | 0.05 | 0.05 | -0.5 | -0.5 | 0.05 | 0.05 | 0.05 | 0.05 | -0.5 | -0.5 |
| 14 | 0.05 | 0.05 | -0.5 | -0.5 | -0.5 | -0.5 | 0.05 | 0.05 | -0.5 | -0.5 | 0.05 | 0.05 | 0.05 | 0.05 | -0.5 | -0.5 |
| 15 | 0.05 | 0.05 | 0.05 | 0.05 | 0.05 | 0.05 | 0.05 | 0.05 | -0.5 | -0.5 | 0.05 | 0.05 | -0.5 | -0.5 | 0.05 | 0.05 |
| 16 | 0.05 | 0.05 | 0.05 | 0.05 | 0.05 | 0.05 | 0.05 | 0.05 | -0.5 | -0.5 | 0.05 | 0.05 | -0.5 | -0.5 | 0.05 | 0.05 |
| 17 | 0.05 | 0.05 | -0.5 | -0.5 | -0.5 | -0.5 | 0.05 | 0.05 | -0.5 | -0.5 | 0.05 | 0.05 | 0.05 | 0.05 | -0.5 | -0.5 |
| 18 | 0.05 | 0.05 | -0.5 | -0.5 | -0.5 | -0.5 | 0.05 | 0.05 | -0.5 | -0.5 | 0.05 | 0.05 | 0.05 | 0.05 | -0.5 | -0.5 |
| 19 | 0.05 | 0.05 | -0.5 | -0.5 | -0.5 | -0.5 | 0.05 | 0.05 | 0.05 | 0.05 | 0.05 | 0.05 | -0.5 | -0.5 | -0.5 | -0.5 |
| 20 | 0.05 | 0.05 | -0.5 | -0.5 | -0.5 | -0.5 | 0.05 | 0.05 | 0.05 | 0.05 | 0.05 | 0.05 | -0.5 | -0.5 | -0.5 | -0.5 |
| 21 | 0.05 | 0.05 | 0.05 | 0.05 | -0.5 | -0.5 | 0.05 | 0.05 | 0.05 | 0.05 | 0.05 | 0.05 | -0.5 | -0.5 | -0.5 | -0.5 |
| 22 | 0.05 | 0.05 | 0.05 | 0.05 | -0.5 | -0.5 | 0.05 | 0.05 | 0.05 | 0.05 | 0.05 | 0.05 | -0.5 | -0.5 | -0.5 | -0.5 |
| 23 | 0.05 | 0.05 | -0.5 | -0.5 | -0.5 | -0.5 | -0.5 | -0.5 | -0.5 | -0.5 | 0.05 | 0.05 | 0.05 | 0.05 | -0.5 | -0.5 |
| 24 | 0.05 | 0.05 | -0.5 | -0.5 | -0.5 | -0.5 | -0.5 | -0.5 | -0.5 | -0.5 | 0.05 | 0.05 | 0.05 | 0.05 | -0.5 | -0.5 |
| 25 | 0.05 | 0.05 | -0.5 | -0.5 | -0.5 | -0.5 | 0.05 | 0.05 | -0.5 | -0.5 | 0.05 | 0.05 | 0.05 | 0.05 | -0.5 | -0.5 |
| 26 | 0.05 | 0.05 | -0.5 | -0.5 | -0.5 | -0.5 | 0.05 | 0.05 | -0.5 | -0.5 | 0.05 | 0.05 | 0.05 | 0.05 | -0.5 | -0.5 |
| 27 | 0.05 | 0.05 | 0.05 | 0.05 | 0.05 | 0.05 | 0.05 | 0.05 | 0.05 | 0.05 | 0.05 | 0.05 | 0.05 | 0.05 | 0.05 | 0.05 |
| 28 | 0.05 | 0.05 | 0.05 | 0.05 | 0.05 | 0.05 | 0.05 | 0.05 | 0.05 | 0.05 | 0.05 | 0.05 | 0.05 | 0.05 | 0.05 | 0.05 |
| 29 | 0.05 | 0.05 | 0.05 | 0.05 | 0.05 | 0.05 | 0.05 | 0.05 | 0.05 | 0.05 | 0.05 | 0.05 | 0.05 | 0.05 | 0.05 | 0.05 |
| 30 | 0.05 | 0.05 | 0.05 | 0.05 | 0.05 | 0.05 | 0.05 | 0.05 | 0.05 | 0.05 | 0.05 | 0.05 | 0.05 | 0.05 | 0.05 | 0.05 |
| 31 | 0.05 | 0.05 | 0.05 | 0.05 | 0.05 | 0.05 | 0.05 | 0.05 | -0.5 | -0.5 | 0.05 | 0.05 | -0.5 | -0.5 | 0.05 | 0.05 |
| 32 | 0.05 | 0.05 | 0.05 | 0.05 | 0.05 | 0.05 | 0.05 | 0.05 | -0.5 | -0.5 | 0.05 | 0.05 | -0.5 | -0.5 | 0.05 | 0.05 |
| 33 | 0.05 | 0.05 | 0.05 | 0.05 | -0.5 | -0.5 | 0.05 | 0.05 | 0.05 | 0.05 | 0.05 | 0.05 | 0.05 | 0.05 | -0.5 | -0.5 |
| 34 | 0.05 | 0.05 | 0.05 | 0.05 | -0.5 | -0.5 | 0.05 | 0.05 | 0.05 | 0.05 | 0.05 | 0.05 | 0.05 | 0.05 | -0.5 | -0.5 |
| 35 | 0.05 | 0.05 | -0.5 | -0.5 | -0.5 | -0.5 | 0.05 | 0.05 | -0.5 | -0.5 | 0.05 | 0.05 | 0.05 | 0.05 | -0.5 | -0.5 |
| 36 | 0.05 | 0.05 | -0.5 | -0.5 | -0.5 | -0.5 | 0.05 | 0.05 | -0.5 | -0.5 | 0.05 | 0.05 | 0.05 | 0.05 | -0.5 | -0.5 |
| 37 | -0.5 | -0.5 | -0.5 | -0.5 | -0.5 | -0.5 | -0.5 | -0.5 | -0.5 | -0.5 | -0.5 | -0.5 | -0.5 | -0.5 | -0.5 | -0.5 |
| 38 | -0.5 | -0.5 | -0.5 | -0.5 | -0.5 | -0.5 | -0.5 | -0.5 | -0.5 | -0.5 | -0.5 | -0.5 | -0.5 | -0.5 | -0.5 | -0.5 |
| 39 | 0.05 | 0.05 | -0.5 | -0.5 | -0.5 | -0.5 | -0.5 | -0.5 | -0.5 | -0.5 | 0.05 | 0.05 | 0.05 | 0.05 | -0.5 | -0.5 |
| 40 | 0.05 | 0.05 | -0.5 | -0.5 | -0.5 | -0.5 | -0.5 | -0.5 | -0.5 | -0.5 | 0.05 | 0.05 | 0.05 | 0.05 | -0.5 | -0.5 |
| 41 | -0.5 | -0.5 | -0.5 | -0.5 | -0.5 | -0.5 | -0.5 | -0.5 | -0.5 | -0.5 | -0.5 | -0.5 | -0.5 | -0.5 | -0.5 | -0.5 |
| 42 | -0.5 | -0.5 | -0.5 | -0.5 | -0.5 | -0.5 | -0.5 | -0.5 | -0.5 | -0.5 | -0.5 | -0.5 | -0.5 | -0.5 | -0.5 | -0.5 |
| 43 | 0.05 | 0.05 | -0.5 | -0.5 | -0.5 | -0.5 | 0.05 | 0.05 | -0.5 | -0.5 | 0.05 | 0.05 | 0.05 | 0.05 | -0.5 | -0.5 |
| 44 | 0.05 | 0.05 | -0.5 | -0.5 | -0.5 | -0.5 | 0.05 | 0.05 | -0.5 | -0.5 | 0.05 | 0.05 | 0.05 | 0.05 | -0.5 | -0.5 |
| 45 | -0.5 | -0.5 | -0.5 | -0.5 | -0.5 | -0.5 | -0.5 | -0.5 | -0.5 | -0.5 | -0.5 | -0.5 | -0.5 | -0.5 | -0.5 | -0.5 |
| 46 | -0.5 | -0.5 | -0.5 | -0.5 | -0.5 | -0.5 | -0.5 | -0.5 | -0.5 | -0.5 | -0.5 | -0.5 | -0.5 | -0.5 | -0.5 | -0.5 |
| 47 | -0.5 | -0.5 | -0.5 | -0.5 | -0.5 | -0.5 | -0.5 | -0.5 | -0.5 | -0.5 | -0.5 | -0.5 | -0.5 | -0.5 | -0.5 | -0.5 |
| 48 | -0.5 | -0.5 | -0.5 | -0.5 | -0.5 | -0.5 | -0.5 | -0.5 | -0.5 | -0.5 | -0.5 | -0.5 | -0.5 | -0.5 | -0.5 | -0.5 |
| 49 | -0.5 | -0.5 | -0.5 | -0.5 | -0.5 | -0.5 | -0.5 | -0.5 | -0.5 | -0.5 | -0.5 | -0.5 | -0.5 | -0.5 | -0.5 | -0.5 |

**B3.2 Table: PFC → PFC Recurrent weight matrix**

Continued from previous page. Continued further on next page.

| | 17 | 18 | 19 | 20 | 21 | 22 | 23 | 24 | 25 | 26 | 27 | 28 | 29 | 30 | 31 | 32 | 33 |
|---|---|---|---|---|---|---|---|---|---|---|---|---|---|---|---|---|---|
| 1 | 0.05 | 0.05 | 0.05 | 0.05 | 0.05 | 0.05 | 0.05 | 0.05 | 0.05 | 0.05 | 0.05 | 0.05 | 0.05 | 0.05 | 0.05 | 0.05 | 0.05 |
| 2 | 0.05 | 0.05 | 0.05 | 0.05 | 0.05 | 0.05 | 0.05 | 0.05 | 0.05 | 0.05 | 0.05 | 0.05 | 0.05 | 0.05 | 0.05 | 0.05 | 0.05 |
| 3 | -0.5 | -0.5 | -0.5 | -0.5 | 0.05 | 0.05 | -0.5 | -0.5 | -0.5 | -0.5 | 0.05 | 0.05 | 0.05 | 0.05 | 0.05 | 0.05 | 0.05 |
| 4 | -0.5 | -0.5 | -0.5 | -0.5 | 0.05 | 0.05 | -0.5 | -0.5 | -0.5 | -0.5 | 0.05 | 0.05 | 0.05 | 0.05 | 0.05 | 0.05 | 0.05 |
| 5 | -0.5 | -0.5 | -0.5 | -0.5 | -0.5 | -0.5 | -0.5 | -0.5 | -0.5 | -0.5 | 0.05 | 0.05 | 0.05 | 0.05 | 0.05 | 0.05 | -0.5 |
| 6 | -0.5 | -0.5 | -0.5 | -0.5 | -0.5 | -0.5 | -0.5 | -0.5 | -0.5 | -0.5 | 0.05 | 0.05 | 0.05 | 0.05 | 0.05 | 0.05 | -0.5 |
| 7 | 0.05 | 0.05 | 0.05 | 0.05 | 0.05 | 0.05 | -0.5 | -0.5 | 0.05 | 0.05 | 0.05 | 0.05 | 0.05 | 0.05 | 0.05 | 0.05 | 0.05 |
| 8 | 0.05 | 0.05 | 0.05 | 0.05 | 0.05 | 0.05 | -0.5 | -0.5 | 0.05 | 0.05 | 0.05 | 0.05 | 0.05 | 0.05 | 0.05 | 0.05 | 0.05 |
| 9 | -0.5 | -0.5 | 0.05 | 0.05 | 0.05 | 0.05 | -0.5 | -0.5 | -0.5 | -0.5 | 0.05 | 0.05 | 0.05 | 0.05 | -0.5 | -0.5 | 0.05 |
| 10 | -0.5 | -0.5 | 0.05 | 0.05 | 0.05 | 0.05 | -0.5 | -0.5 | -0.5 | -0.5 | 0.05 | 0.05 | 0.05 | 0.05 | -0.5 | -0.5 | 0.05 |
| 11 | 0.05 | 0.05 | 0.05 | 0.05 | 0.05 | 0.05 | 0.05 | 0.05 | 0.05 | 0.05 | 0.05 | 0.05 | 0.05 | 0.05 | 0.05 | 0.05 | 0.05 |
| 12 | 0.05 | 0.05 | 0.05 | 0.05 | 0.05 | 0.05 | 0.05 | 0.05 | 0.05 | 0.05 | 0.05 | 0.05 | 0.05 | 0.05 | 0.05 | 0.05 | 0.05 |
| 13 | 0.05 | 0.05 | -0.5 | -0.5 | -0.5 | -0.5 | 0.05 | 0.05 | 0.05 | 0.05 | 0.05 | 0.05 | 0.05 | 0.05 | -0.5 | -0.5 | 0.05 |
| 14 | 0.05 | 0.05 | -0.5 | -0.5 | -0.5 | -0.5 | 0.05 | 0.05 | 0.05 | 0.05 | 0.05 | 0.05 | 0.05 | 0.05 | -0.5 | -0.5 | 0.05 |
| 15 | -0.5 | -0.5 | -0.5 | -0.5 | -0.5 | -0.5 | -0.5 | -0.5 | -0.5 | -0.5 | 0.05 | 0.05 | 0.05 | 0.05 | 0.05 | 0.05 | -0.5 |
| 16 | -0.5 | -0.5 | -0.5 | -0.5 | -0.5 | -0.5 | -0.5 | -0.5 | -0.5 | -0.5 | 0.05 | 0.05 | 0.05 | 0.05 | 0.05 | 0.05 | -0.5 |
| 17 | 0.05 | 0.05 | -0.5 | -0.5 | -0.5 | -0.5 | 0.05 | 0.05 | 0.05 | 0.05 | 0.05 | 0.05 | 0.05 | 0.05 | -0.5 | -0.5 | 0.05 |
| 18 | 0.05 | 0.05 | -0.5 | -0.5 | -0.5 | -0.5 | 0.05 | 0.05 | 0.05 | 0.05 | 0.05 | 0.05 | 0.05 | 0.05 | -0.5 | -0.5 | 0.05 |
| 19 | -0.5 | -0.5 | 0.05 | 0.05 | 0.05 | 0.05 | -0.5 | -0.5 | -0.5 | -0.5 | 0.05 | 0.05 | -0.5 | -0.5 | -0.5 | -0.5 | 0.05 |
| 20 | -0.5 | -0.5 | 0.05 | 0.05 | 0.05 | 0.05 | -0.5 | -0.5 | -0.5 | -0.5 | 0.05 | 0.05 | -0.5 | -0.5 | -0.5 | -0.5 | 0.05 |
| 21 | -0.5 | -0.5 | 0.05 | 0.05 | 0.05 | 0.05 | -0.5 | -0.5 | -0.5 | -0.5 | 0.05 | 0.05 | 0.05 | 0.05 | -0.5 | -0.5 | 0.05 |
| 22 | -0.5 | -0.5 | 0.05 | 0.05 | 0.05 | 0.05 | -0.5 | -0.5 | -0.5 | -0.5 | 0.05 | 0.05 | 0.05 | 0.05 | -0.5 | -0.5 | 0.05 |
| 23 | 0.05 | 0.05 | -0.5 | -0.5 | -0.5 | -0.5 | 0.05 | 0.05 | 0.05 | 0.05 | 0.05 | 0.05 | -0.5 | -0.5 | -0.5 | -0.5 | 0.05 |
| 24 | 0.05 | 0.05 | -0.5 | -0.5 | -0.5 | -0.5 | 0.05 | 0.05 | 0.05 | 0.05 | 0.05 | 0.05 | -0.5 | -0.5 | -0.5 | -0.5 | 0.05 |
| 25 | 0.05 | 0.05 | -0.5 | -0.5 | -0.5 | -0.5 | 0.05 | 0.05 | 0.05 | 0.05 | 0.05 | 0.05 | -0.5 | -0.5 | -0.5 | -0.5 | 0.05 |
| 26 | 0.05 | 0.05 | -0.5 | -0.5 | -0.5 | -0.5 | 0.05 | 0.05 | 0.05 | 0.05 | 0.05 | 0.05 | -0.5 | -0.5 | -0.5 | -0.5 | 0.05 |
| 27 | 0.05 | 0.05 | 0.05 | 0.05 | 0.05 | 0.05 | 0.05 | 0.05 | 0.05 | 0.05 | 0.05 | 0.05 | -0.5 | -0.5 | 0.05 | 0.05 | 0.05 |
| 28 | 0.05 | 0.05 | 0.05 | 0.05 | 0.05 | 0.05 | 0.05 | 0.05 | 0.05 | 0.05 | 0.05 | 0.05 | -0.5 | -0.5 | 0.05 | 0.05 | 0.05 |
| 29 | 0.05 | 0.05 | -0.5 | -0.5 | 0.05 | 0.05 | -0.5 | -0.5 | -0.5 | -0.5 | -1 | -1 | 0.05 | 0.05 | -0.5 | -0.5 | -1 |
| 30 | 0.05 | 0.05 | -0.5 | -0.5 | 0.05 | 0.05 | -0.5 | -0.5 | -0.5 | -0.5 | -1 | -1 | 0.05 | 0.05 | -0.5 | -0.5 | -1 |
| 31 | -0.5 | -0.5 | -0.5 | -0.5 | -0.5 | -0.5 | -0.5 | -0.5 | -0.5 | -0.5 | 0.05 | 0.05 | -0.5 | -0.5 | 0.05 | 0.05 | -0.5 |
| 32 | -0.5 | -0.5 | -0.5 | -0.5 | -0.5 | -0.5 | -0.5 | -0.5 | -0.5 | -0.5 | 0.05 | 0.05 | -0.5 | -0.5 | 0.05 | 0.05 | -0.5 |
| 33 | 0.05 | 0.05 | 0.05 | 0.05 | 0.05 | 0.05 | 0.05 | 0.05 | 0.05 | 0.05 | 0.05 | 0.05 | -0.5 | -0.5 | -0.5 | -0.5 | 0.05 |
| 34 | 0.05 | 0.05 | 0.05 | 0.05 | 0.05 | 0.05 | 0.05 | 0.05 | 0.05 | 0.05 | 0.05 | 0.05 | -0.5 | -0.5 | -0.5 | -0.5 | 0.05 |
| 35 | 0.05 | 0.05 | -0.5 | -0.5 | -0.5 | -0.5 | -0.5 | -0.5 | 0.05 | 0.05 | -1 | -1 | -0.5 | -0.5 | -0.5 | -0.5 | -1 |
| 36 | 0.05 | 0.05 | -0.5 | -0.5 | -0.5 | -0.5 | -0.5 | -0.5 | 0.05 | 0.05 | -1 | -1 | -0.5 | -0.5 | -0.5 | -0.5 | -1 |
| 37 | -0.5 | -0.5 | -0.5 | -0.5 | -0.5 | -0.5 | -0.5 | -0.5 | -0.5 | -0.5 | -0.5 | -0.5 | -0.5 | -0.5 | -0.5 | -0.5 | -0.5 |
| 38 | -0.5 | -0.5 | -0.5 | -0.5 | -0.5 | -0.5 | -0.5 | -0.5 | -0.5 | -0.5 | -0.5 | -0.5 | -0.5 | -0.5 | -0.5 | -0.5 | -0.5 |
| 39 | 0.05 | 0.05 | -0.5 | -0.5 | -0.5 | -0.5 | -0.5 | -0.5 | 0.05 | 0.05 | 0.05 | 0.05 | 0.05 | 0.05 | -0.5 | -0.5 | 0.05 |
| 40 | 0.05 | 0.05 | -0.5 | -0.5 | -0.5 | -0.5 | -0.5 | -0.5 | 0.05 | 0.05 | 0.05 | 0.05 | 0.05 | 0.05 | -0.5 | -0.5 | 0.05 |
| 41 | -0.5 | -0.5 | -0.5 | -0.5 | -0.5 | -0.5 | -0.5 | -0.5 | -0.5 | -0.5 | -0.5 | -0.5 | -0.5 | -0.5 | -0.5 | -0.5 | -0.5 |
| 42 | -0.5 | -0.5 | -0.5 | -0.5 | -0.5 | -0.5 | -0.5 | -0.5 | -0.5 | -0.5 | -0.5 | -0.5 | -0.5 | -0.5 | -0.5 | -0.5 | -0.5 |
| 43 | 0.05 | 0.05 | -0.5 | -0.5 | -0.5 | -0.5 | 0.05 | 0.05 | 0.05 | 0.05 | 0.05 | 0.05 | 0.05 | 0.05 | -0.5 | -0.5 | 0.05 |
| 44 | 0.05 | 0.05 | -0.5 | -0.5 | -0.5 | -0.5 | 0.05 | 0.05 | 0.05 | 0.05 | 0.05 | 0.05 | 0.05 | 0.05 | -0.5 | -0.5 | 0.05 |
| 45 | -0.5 | -0.5 | -0.5 | -0.5 | -0.5 | -0.5 | -0.5 | -0.5 | -0.5 | -0.5 | -0.5 | -0.5 | -0.5 | -0.5 | -0.5 | -0.5 | -0.5 |
| 46 | -0.5 | -0.5 | -0.5 | -0.5 | -0.5 | -0.5 | -0.5 | -0.5 | -0.5 | -0.5 | -0.5 | -0.5 | -0.5 | -0.5 | -0.5 | -0.5 | -0.5 |
| 47 | -0.5 | -0.5 | -0.5 | -0.5 | -0.5 | -0.5 | -0.5 | -0.5 | -0.5 | -0.5 | -0.5 | -0.5 | -0.5 | -0.5 | -0.5 | -0.5 | -0.5 |
| 48 | -0.5 | -0.5 | -0.5 | -0.5 | -0.5 | -0.5 | -0.5 | -0.5 | -0.5 | -0.5 | -0.5 | -0.5 | -0.5 | -0.5 | -0.5 | -0.5 | -0.5 |
| 49 | -0.5 | -0.5 | -0.5 | -0.5 | -0.5 | -0.5 | -0.5 | -0.5 | -0.5 | -0.5 | -0.5 | -0.5 | -0.5 | -0.5 | -0.5 | -0.5 | -0.5 |

**B3.2 Table: PFC → PFC Recurrent weight matrix**

Continued from previous page.

| | 34 | 35 | 36 | 37 | 38 | 39 | 40 | 41 | 42 | 43 | 44 | 45 | 46 | 47 | 48 | 49 |
|---|---|---|---|---|---|---|---|---|---|---|---|---|---|---|---|---|
| 1 | 0.05 | 0.05 | 0.05 | -0.5 | -0.5 | 0.05 | 0.05 | -0.5 | -0.5 | 0.05 | 0.05 | -0.5 | -0.5 | -0.5 | -0.5 | -0.5 |
| 2 | 0.05 | 0.05 | 0.05 | -0.5 | -0.5 | 0.05 | 0.05 | -0.5 | -0.5 | 0.05 | 0.05 | -0.5 | -0.5 | -0.5 | -0.5 | -0.5 |
| 3 | 0.05 | -0.5 | -0.5 | -0.5 | -0.5 | -0.5 | -0.5 | -0.5 | -0.5 | -0.5 | -0.5 | -0.5 | -0.5 | -0.5 | -0.5 | -0.5 |
| 4 | 0.05 | -0.5 | -0.5 | -0.5 | -0.5 | -0.5 | -0.5 | -0.5 | -0.5 | -0.5 | -0.5 | -0.5 | -0.5 | -0.5 | -0.5 | -0.5 |
| 5 | -0.5 | -0.5 | -0.5 | -0.5 | -0.5 | -0.5 | -0.5 | -0.5 | -0.5 | -0.5 | -0.5 | -0.5 | -0.5 | -0.5 | -0.5 | -0.5 |
| 6 | -0.5 | -0.5 | -0.5 | -0.5 | -0.5 | -0.5 | -0.5 | -0.5 | -0.5 | -0.5 | -0.5 | -0.5 | -0.5 | -0.5 | -0.5 | -0.5 |
| 7 | 0.05 | 0.05 | 0.05 | -0.5 | -0.5 | -0.5 | -0.5 | -0.5 | -0.5 | 0.05 | 0.05 | -0.5 | -0.5 | -0.5 | -0.5 | -0.5 |
| 8 | 0.05 | 0.05 | 0.05 | -0.5 | -0.5 | -0.5 | -0.5 | -0.5 | -0.5 | 0.05 | 0.05 | -0.5 | -0.5 | -0.5 | -0.5 | -0.5 |
| 9 | 0.05 | -0.5 | -0.5 | -0.5 | -0.5 | -0.5 | -0.5 | -0.5 | -0.5 | -0.5 | -0.5 | -0.5 | -0.5 | -0.5 | -0.5 | -0.5 |
| 10 | 0.05 | -0.5 | -0.5 | -0.5 | -0.5 | -0.5 | -0.5 | -0.5 | -0.5 | -0.5 | -0.5 | -0.5 | -0.5 | -0.5 | -0.5 | -0.5 |
| 11 | 0.05 | 0.05 | 0.05 | -0.5 | -0.5 | 0.05 | 0.05 | -0.5 | -0.5 | 0.05 | 0.05 | -0.5 | -0.5 | -0.5 | -0.5 | -0.5 |
| 12 | 0.05 | 0.05 | 0.05 | -0.5 | -0.5 | 0.05 | 0.05 | -0.5 | -0.5 | 0.05 | 0.05 | -0.5 | -0.5 | -0.5 | -0.5 | -0.5 |
| 13 | 0.05 | 0.05 | 0.05 | -0.5 | -0.5 | 0.05 | 0.05 | -0.5 | -0.5 | 0.05 | 0.05 | -0.5 | -0.5 | -0.5 | -0.5 | -0.5 |
| 14 | 0.05 | 0.05 | 0.05 | -0.5 | -0.5 | 0.05 | 0.05 | -0.5 | -0.5 | 0.05 | 0.05 | -0.5 | -0.5 | -0.5 | -0.5 | -0.5 |
| 15 | -0.5 | -0.5 | -0.5 | -0.5 | -0.5 | -0.5 | -0.5 | -0.5 | -0.5 | -0.5 | -0.5 | -0.5 | -0.5 | -0.5 | -0.5 | -0.5 |
| 16 | -0.5 | -0.5 | -0.5 | -0.5 | -0.5 | -0.5 | -0.5 | -0.5 | -0.5 | -0.5 | -0.5 | -0.5 | -0.5 | -0.5 | -0.5 | -0.5 |
| 17 | 0.05 | 0.05 | 0.05 | -0.5 | -0.5 | 0.05 | 0.05 | -0.5 | -0.5 | 0.05 | 0.05 | -0.5 | -0.5 | -0.5 | -0.5 | -0.5 |
| 18 | 0.05 | 0.05 | 0.05 | -0.5 | -0.5 | 0.05 | 0.05 | -0.5 | -0.5 | 0.05 | 0.05 | -0.5 | -0.5 | -0.5 | -0.5 | -0.5 |
| 19 | 0.05 | -0.5 | -0.5 | -0.5 | -0.5 | -0.5 | -0.5 | -0.5 | -0.5 | -0.5 | -0.5 | -0.5 | -0.5 | -0.5 | -0.5 | -0.5 |
| 20 | 0.05 | -0.5 | -0.5 | -0.5 | -0.5 | -0.5 | -0.5 | -0.5 | -0.5 | -0.5 | -0.5 | -0.5 | -0.5 | -0.5 | -0.5 | -0.5 |
| 21 | 0.05 | -0.5 | -0.5 | -0.5 | -0.5 | -0.5 | -0.5 | -0.5 | -0.5 | -0.5 | -0.5 | -0.5 | -0.5 | -0.5 | -0.5 | -0.5 |
| 22 | 0.05 | -0.5 | -0.5 | -0.5 | -0.5 | -0.5 | -0.5 | -0.5 | -0.5 | -0.5 | -0.5 | -0.5 | -0.5 | -0.5 | -0.5 | -0.5 |
| 23 | 0.05 | -0.5 | -0.5 | -0.5 | -0.5 | -0.5 | -0.5 | -0.5 | -0.5 | 0.05 | 0.05 | -0.5 | -0.5 | -0.5 | -0.5 | -0.5 |
| 24 | 0.05 | -0.5 | -0.5 | -0.5 | -0.5 | -0.5 | -0.5 | -0.5 | -0.5 | 0.05 | 0.05 | -0.5 | -0.5 | -0.5 | -0.5 | -0.5 |
| 25 | 0.05 | 0.05 | 0.05 | -0.5 | -0.5 | 0.05 | 0.05 | -0.5 | -0.5 | 0.05 | 0.05 | -0.5 | -0.5 | -0.5 | -0.5 | -0.5 |
| 26 | 0.05 | 0.05 | 0.05 | -0.5 | -0.5 | 0.05 | 0.05 | -0.5 | -0.5 | 0.05 | 0.05 | -0.5 | -0.5 | -0.5 | -0.5 | -0.5 |
| 27 | 0.05 | -0.5 | -0.5 | -0.5 | -0.5 | 0.05 | 0.05 | -0.5 | -0.5 | 0.05 | 0.05 | -0.5 | -0.5 | -0.5 | -0.5 | -0.5 |
| 28 | 0.05 | -0.5 | -0.5 | -0.5 | -0.5 | 0.05 | 0.05 | -0.5 | -0.5 | 0.05 | 0.05 | -0.5 | -0.5 | -0.5 | -0.5 | -0.5 |
| 29 | -1 | -1 | -0.5 | -0.5 | -0.5 | 0.05 | 0.05 | -0.5 | -0.5 | 0.05 | 0.05 | -0.5 | -0.5 | -0.5 | -0.5 | -0.5 |
| 30 | -1 | -0.5 | -1 | -0.5 | -0.5 | 0.05 | 0.05 | -0.5 | -0.5 | 0.05 | 0.05 | -0.5 | -0.5 | -0.5 | -0.5 | -0.5 |
| 31 | -0.5 | -0.5 | -0.5 | -0.5 | -0.5 | -0.5 | -0.5 | -0.5 | -0.5 | -0.5 | -0.5 | -0.5 | -0.5 | -0.5 | -0.5 | -0.5 |
| 32 | -0.5 | -0.5 | -0.5 | -0.5 | -0.5 | -0.5 | -0.5 | -0.5 | -0.5 | -0.5 | -0.5 | -0.5 | -0.5 | -0.5 | -0.5 | -0.5 |
| 33 | 0.05 | -0.5 | -0.5 | -0.5 | -0.5 | 0.05 | 0.05 | -0.5 | -0.5 | 0.05 | 0.05 | -0.5 | -0.5 | -0.5 | -0.5 | -0.5 |
| 34 | 0.05 | -0.5 | -0.5 | -0.5 | -0.5 | 0.05 | 0.05 | -0.5 | -0.5 | 0.05 | 0.05 | -0.5 | -0.5 | -0.5 | -0.5 | -0.5 |
| 35 | -1 | 0.05 | 0.05 | -0.5 | -0.5 | 0.05 | 0.05 | -0.5 | -0.5 | 0.05 | 0.05 | -0.5 | -0.5 | -0.5 | -0.5 | -0.5 |
| 36 | -1 | 0.05 | 0.05 | -0.5 | -0.5 | 0.05 | 0.05 | -0.5 | -0.5 | 0.05 | 0.05 | -0.5 | -0.5 | -0.5 | -0.5 | -0.5 |
| 37 | -0.5 | -0.5 | -0.5 | -0.5 | -0.5 | -0.5 | -0.5 | -0.5 | -0.5 | -0.5 | -0.5 | -0.5 | -0.5 | -0.5 | -0.5 | -0.5 |
| 38 | -0.5 | -0.5 | -0.5 | -0.5 | -0.5 | -0.5 | -0.5 | -0.5 | -0.5 | -0.5 | -0.5 | -0.5 | -0.5 | -0.5 | -0.5 | -0.5 |
| 39 | 0.05 | 0.05 | 0.05 | -0.5 | -0.5 | 0.05 | 0.05 | -0.5 | -0.5 | -1 | -0.5 | -0.5 | -0.5 | -0.5 | -0.5 | -0.5 |
| 40 | 0.05 | 0.05 | 0.05 | -0.5 | -0.5 | 0.05 | 0.05 | -0.5 | -0.5 | -0.5 | -1 | -0.5 | -0.5 | -0.5 | -0.5 | -0.5 |
| 41 | -0.5 | -0.5 | -0.5 | -0.5 | -0.5 | -0.5 | -0.5 | -0.5 | -0.5 | -0.5 | -0.5 | -0.5 | -0.5 | -0.5 | -0.5 | -0.5 |
| 42 | -0.5 | -0.5 | -0.5 | -0.5 | -0.5 | -0.5 | -0.5 | -0.5 | -0.5 | -0.5 | -0.5 | -0.5 | -0.5 | -0.5 | -0.5 | -0.5 |
| 43 | 0.05 | 0.05 | 0.05 | -0.5 | -0.5 | -1 | -0.5 | -0.5 | -0.5 | 0.05 | 0.05 | -0.5 | -0.5 | -0.5 | -0.5 | -0.5 |
| 44 | 0.05 | 0.05 | 0.05 | -0.5 | -0.5 | -0.5 | -1 | -0.5 | -0.5 | 0.05 | 0.05 | -0.5 | -0.5 | -0.5 | -0.5 | -0.5 |
| 45 | -0.5 | -0.5 | -0.5 | -0.5 | -0.5 | -0.5 | -0.5 | -0.5 | -0.5 | -0.5 | -0.5 | -0.5 | -0.5 | -0.5 | -0.5 | -0.5 |
| 46 | -0.5 | -0.5 | -0.5 | -0.5 | -0.5 | -0.5 | -0.5 | -0.5 | -0.5 | -0.5 | -0.5 | -0.5 | -0.5 | -0.5 | -0.5 | -0.5 |
| 47 | -0.5 | -0.5 | -0.5 | -0.5 | -0.5 | -0.5 | -0.5 | -0.5 | -0.5 | -0.5 | -0.5 | -0.5 | -0.5 | -0.5 | -0.5 | -0.5 |
| 48 | -0.5 | -0.5 | -0.5 | -0.5 | -0.5 | -0.5 | -0.5 | -0.5 | -0.5 | -0.5 | -0.5 | -0.5 | -0.5 | -0.5 | -0.5 | -0.5 |
| 49 | -0.5 | -0.5 | -0.5 | -0.5 | -0.5 | -0.5 | -0.5 | -0.5 | -0.5 | -0.5 | -0.5 | -0.5 | -0.5 | -0.5 | -0.5 | -0.5 |

**B3.3 Table: PFC → Transition node matrix**

Rows = Source PFC nodes; Columns = Target transition nodes.

Continued on next page.

| | 1 | 2 | 3 | 4 | 5 | 6 | 7 | 8 | 9 | 10 | 11 |
|---|---|---|---|---|---|---|---|---|---|---|---|
| 1 | -0.01 | 0.025 | -0.01 | 0.025 | -0.01 | 0.025 | 0.025 | 0.025 | 0.025 | 0.025 | 0.025 |
| 2 | -0.01 | 0.025 | -0.01 | 0.025 | -0.01 | 0.025 | 0.025 | 0.025 | 0.025 | 0.025 | 0.025 |
| 3 | -0.01 | 0.025 | -0.01 | -0.01 | -0.01 | -0.01 | 0.025 | 0.025 | -0.01 | -0.01 | -0.01 |
| 4 | -0.01 | 0.025 | -0.01 | -0.01 | -0.01 | -0.01 | 0.025 | 0.025 | -0.01 | -0.01 | -0.01 |
| 5 | -0.01 | 0.025 | -0.01 | 0.025 | -0.01 | 0.025 | 0.025 | -0.01 | 0.025 | -0.01 | 0.025 |
| 6 | -0.01 | 0.025 | -0.01 | 0.025 | -0.01 | 0.025 | 0.025 | -0.01 | 0.025 | -0.01 | 0.025 |
| 7 | -0.01 | -0.01 | -0.01 | 0.025 | -0.01 | -0.01 | -0.01 | -0.01 | 0.025 | 0.025 | -0.01 |
| 8 | -0.01 | -0.01 | -0.01 | 0.025 | -0.01 | -0.01 | -0.01 | -0.01 | 0.025 | 0.025 | -0.01 |
| 9 | -0.01 | -0.01 | -0.01 | -0.01 | -0.01 | -0.01 | -0.01 | 0.025 | -0.01 | 0.025 | -0.01 |
| 10 | -0.01 | -0.01 | -0.01 | -0.01 | -0.01 | -0.01 | -0.01 | 0.025 | -0.01 | 0.025 | -0.01 |
| 11 | -0.01 | -0.01 | -0.01 | -0.01 | -0.01 | 0.025 | -0.01 | -0.01 | -0.01 | -0.01 | 0.025 |
| 12 | -0.01 | -0.01 | -0.01 | -0.01 | -0.01 | 0.025 | -0.01 | -0.01 | -0.01 | -0.01 | 0.025 |
| 13 | -0.01 | -0.01 | -0.01 | -0.01 | -0.01 | -0.01 | -0.01 | -0.01 | -0.01 | -0.01 | -0.01 |
| 14 | -0.01 | -0.01 | -0.01 | -0.01 | -0.01 | -0.01 | -0.01 | -0.01 | -0.01 | -0.01 | -0.01 |
| 15 | -0.01 | 0.025 | -0.01 | 0.025 | -0.01 | 0.025 | 0.025 | -0.01 | 0.025 | -0.01 | 0.025 |
| 16 | -0.01 | 0.025 | -0.01 | 0.025 | -0.01 | 0.025 | 0.025 | -0.01 | 0.025 | -0.01 | 0.025 |
| 17 | -0.01 | -0.01 | -0.01 | -0.01 | -0.01 | -0.01 | -0.01 | -0.01 | -0.01 | -0.01 | -0.01 |
| 18 | -0.01 | -0.01 | -0.01 | -0.01 | -0.01 | -0.01 | -0.01 | -0.01 | -0.01 | -0.01 | -0.01 |
| 19 | -0.01 | -0.01 | -0.01 | -0.01 | -0.01 | -0.01 | -0.01 | -0.01 | -0.01 | -0.01 | -0.01 |
| 20 | -0.01 | -0.01 | -0.01 | -0.01 | -0.01 | -0.01 | -0.01 | -0.01 | -0.01 | -0.01 | -0.01 |
| 21 | -0.01 | -0.01 | -0.01 | -0.01 | -0.01 | -0.01 | -0.01 | 0.025 | -0.01 | 0.025 | -0.01 |
| 22 | -0.01 | -0.01 | -0.01 | -0.01 | -0.01 | -0.01 | -0.01 | 0.025 | -0.01 | 0.025 | -0.01 |
| 23 | -0.01 | -0.01 | -0.01 | -0.01 | -0.01 | -0.01 | -0.01 | -0.01 | -0.01 | -0.01 | -0.01 |
| 24 | -0.01 | -0.01 | -0.01 | -0.01 | -0.01 | -0.01 | -0.01 | -0.01 | -0.01 | -0.01 | -0.01 |
| 25 | -0.01 | -0.01 | -0.01 | -0.01 | -0.01 | -0.01 | -0.01 | -0.01 | -0.01 | -0.01 | -0.01 |
| 26 | -0.01 | -0.01 | -0.01 | -0.01 | -0.01 | -0.01 | -0.01 | -0.01 | -0.01 | -0.01 | -0.01 |
| 27 | -0.01 | -0.01 | -0.01 | -0.01 | -0.01 | -0.01 | 0.025 | -0.01 | 0.025 | -0.01 | 0.025 |
| 28 | -0.01 | -0.01 | -0.01 | -0.01 | -0.01 | -0.01 | 0.025 | -0.01 | 0.025 | -0.01 | 0.025 |
| 29 | -0.01 | 0.025 | -0.01 | 0.025 | -0.01 | 0.025 | -0.01 | 0.025 | -0.01 | 0.025 | -0.01 |
| 30 | -0.01 | 0.025 | -0.01 | 0.025 | -0.01 | 0.025 | -0.01 | 0.025 | -0.01 | 0.025 | -0.01 |
| 31 | -0.01 | -0.01 | -0.01 | -0.01 | -0.01 | -0.01 | 0.025 | -0.01 | 0.025 | -0.01 | 0.025 |
| 32 | -0.01 | -0.01 | -0.01 | -0.01 | -0.01 | -0.01 | 0.025 | -0.01 | 0.025 | -0.01 | 0.025 |
| 33 | -0.01 | -0.01 | -0.01 | -0.01 | -0.01 | -0.01 | -0.01 | -0.01 | -0.01 | -0.01 | -0.01 |
| 34 | -0.01 | -0.01 | -0.01 | -0.01 | -0.01 | -0.01 | -0.01 | -0.01 | -0.01 | -0.01 | -0.01 |
| 35 | -0.01 | -0.01 | -0.01 | -0.01 | -0.01 | -0.01 | -0.01 | -0.01 | -0.01 | -0.01 | -0.01 |
| 36 | -0.01 | -0.01 | -0.01 | -0.01 | -0.01 | -0.01 | -0.01 | -0.01 | -0.01 | -0.01 | -0.01 |
| 37 | -0.01 | -0.01 | -0.01 | -0.01 | -0.01 | -0.01 | -0.01 | -0.01 | -0.01 | -0.01 | -0.01 |
| 38 | -0.01 | -0.01 | -0.01 | -0.01 | -0.01 | -0.01 | -0.01 | -0.01 | -0.01 | -0.01 | -0.01 |
| 39 | -0.01 | -0.01 | -0.01 | -0.01 | -0.01 | -0.01 | -0.01 | -0.01 | -0.01 | -0.01 | -0.01 |
| 40 | -0.01 | -0.01 | -0.01 | -0.01 | -0.01 | -0.01 | -0.01 | -0.01 | -0.01 | -0.01 | -0.01 |
| 41 | -0.01 | -0.01 | -0.01 | -0.01 | -0.01 | -0.01 | -0.01 | -0.01 | -0.01 | -0.01 | -0.01 |
| 42 | -0.01 | -0.01 | -0.01 | -0.01 | -0.01 | -0.01 | -0.01 | -0.01 | -0.01 | -0.01 | -0.01 |
| 43 | -0.01 | -0.01 | -0.01 | -0.01 | -0.01 | -0.01 | -0.01 | -0.01 | -0.01 | -0.01 | -0.01 |
| 44 | -0.01 | -0.01 | -0.01 | -0.01 | -0.01 | -0.01 | -0.01 | -0.01 | -0.01 | -0.01 | -0.01 |
| 45 | -0.01 | -0.01 | -0.01 | -0.01 | -0.01 | -0.01 | -0.01 | -0.01 | -0.01 | -0.01 | -0.01 |
| 46 | -0.01 | -0.01 | -0.01 | -0.01 | -0.01 | -0.01 | -0.01 | -0.01 | -0.01 | -0.01 | -0.01 |
| 47 | -0.01 | -0.01 | -0.01 | -0.01 | -0.01 | -0.01 | -0.01 | -0.01 | -0.01 | -0.01 | -0.01 |
| 48 | -0.01 | -0.01 | -0.01 | -0.01 | -0.01 | -0.01 | -0.01 | -0.01 | -0.01 | -0.01 | -0.01 |
| 49 | -0.01 | -0.01 | -0.01 | -0.01 | -0.01 | -0.01 | -0.01 | -0.01 | -0.01 | -0.01 | -0.01 |

**B3.3 Table: PFC → Transition node matrix**

Continued from previous page.

| | 12 | 13 | 14 | 15 | 16 | 17 | 18 | 19 | 20 | 21 |
|---|---|---|---|---|---|---|---|---|---|---|
| 1 | 0.025 | 0.025 | 0.025 | 0.025 | 0.025 | 0.025 | 0.025 | -0.01 | 0.025 | 0.025 |
| 2 | 0.025 | 0.025 | 0.025 | 0.025 | 0.025 | 0.025 | 0.025 | -0.01 | 0.025 | 0.025 |
| 3 | -0.01 | -0.01 | -0.01 | -0.01 | -0.01 | -0.01 | -0.01 | -0.01 | -0.01 | -0.01 |
| 4 | -0.01 | -0.01 | -0.01 | -0.01 | -0.01 | -0.01 | -0.01 | -0.01 | -0.01 | -0.01 |
| 5 | -0.01 | -0.01 | -0.01 | -0.01 | -0.01 | -0.01 | -0.01 | -0.01 | -0.01 | -0.01 |
| 6 | -0.01 | -0.01 | -0.01 | -0.01 | -0.01 | -0.01 | -0.01 | -0.01 | -0.01 | -0.01 |
| 7 | -0.01 | 0.025 | 0.025 | 0.025 | -0.01 | -0.01 | -0.01 | -0.01 | -0.01 | -0.01 |
| 8 | -0.01 | 0.025 | 0.025 | 0.025 | -0.01 | -0.01 | -0.01 | -0.01 | -0.01 | -0.01 |
| 9 | 0.025 | 0.025 | -0.01 | -0.01 | 0.025 | -0.01 | -0.01 | -0.01 | -0.01 | -0.01 |
| 10 | 0.025 | 0.025 | -0.01 | -0.01 | 0.025 | -0.01 | -0.01 | -0.01 | -0.01 | -0.01 |
| 11 | 0.025 | -0.01 | -0.01 | -0.01 | 0.025 | 0.025 | 0.025 | -0.01 | 0.025 | 0.025 |
| 12 | 0.025 | -0.01 | -0.01 | -0.01 | 0.025 | 0.025 | 0.025 | -0.01 | 0.025 | 0.025 |
| 13 | -0.01 | -0.01 | 0.025 | 0.025 | -0.01 | 0.025 | 0.025 | -0.01 | 0.025 | 0.025 |
| 14 | -0.01 | -0.01 | 0.025 | 0.025 | -0.01 | 0.025 | 0.025 | -0.01 | 0.025 | 0.025 |
| 15 | -0.01 | -0.01 | -0.01 | -0.01 | -0.01 | -0.01 | -0.01 | -0.01 | -0.01 | -0.01 |
| 16 | -0.01 | -0.01 | -0.01 | -0.01 | -0.01 | -0.01 | -0.01 | -0.01 | -0.01 | -0.01 |
| 17 | -0.01 | -0.01 | 0.025 | 0.025 | -0.01 | 0.025 | 0.025 | -0.01 | 0.025 | 0.025 |
| 18 | -0.01 | -0.01 | 0.025 | 0.025 | -0.01 | 0.025 | 0.025 | -0.01 | 0.025 | 0.025 |
| 19 | -0.01 | 0.025 | -0.01 | -0.01 | 0.025 | -0.01 | -0.01 | -0.01 | -0.01 | -0.01 |
| 20 | -0.01 | 0.025 | -0.01 | -0.01 | 0.025 | -0.01 | -0.01 | -0.01 | -0.01 | -0.01 |
| 21 | 0.025 | 0.025 | -0.01 | -0.01 | 0.025 | -0.01 | -0.01 | -0.01 | -0.01 | -0.01 |
| 22 | 0.025 | 0.025 | -0.01 | -0.01 | 0.025 | -0.01 | -0.01 | -0.01 | -0.01 | -0.01 |
| 23 | -0.01 | -0.01 | -0.01 | -0.01 | -0.01 | -0.01 | -0.01 | -0.01 | 0.025 | -0.01 |
| 24 | -0.01 | -0.01 | -0.01 | -0.01 | -0.01 | -0.01 | -0.01 | -0.01 | 0.025 | -0.01 |
| 25 | -0.01 | -0.01 | -0.01 | 0.025 | -0.01 | -0.01 | 0.025 | -0.01 | 0.025 | 0.025 |
| 26 | -0.01 | -0.01 | -0.01 | 0.025 | -0.01 | -0.01 | 0.025 | -0.01 | 0.025 | 0.025 |
| 27 | -0.01 | 0.025 | -0.01 | -0.01 | 0.025 | -0.01 | -0.01 | -0.01 | 0.025 | -0.01 |
| 28 | -0.01 | 0.025 | -0.01 | -0.01 | 0.025 | -0.01 | -0.01 | -0.01 | 0.025 | -0.01 |
| 29 | 0.025 | -0.01 | 0.025 | -0.01 | -0.01 | 0.025 | -0.01 | -0.01 | -0.01 | -0.01 |
| 30 | 0.025 | -0.01 | 0.025 | -0.01 | -0.01 | 0.025 | -0.01 | -0.01 | -0.01 | -0.01 |
| 31 | -0.01 | -0.01 | -0.01 | -0.01 | -0.01 | -0.01 | -0.01 | -0.01 | -0.01 | -0.01 |
| 32 | -0.01 | -0.01 | -0.01 | -0.01 | -0.01 | -0.01 | -0.01 | -0.01 | -0.01 | -0.01 |
| 33 | -0.01 | 0.025 | -0.01 | -0.01 | 0.025 | -0.01 | -0.01 | -0.01 | 0.025 | -0.01 |
| 34 | -0.01 | 0.025 | -0.01 | -0.01 | 0.025 | -0.01 | -0.01 | -0.01 | 0.025 | -0.01 |
| 35 | -0.01 | -0.01 | -0.01 | 0.025 | -0.01 | -0.01 | 0.025 | -0.01 | -0.01 | 0.025 |
| 36 | -0.01 | -0.01 | -0.01 | 0.025 | -0.01 | -0.01 | 0.025 | -0.01 | -0.01 | 0.025 |
| 37 | -0.01 | -0.01 | -0.01 | -0.01 | -0.01 | -0.01 | -0.01 | -0.01 | -0.01 | -0.01 |
| 38 | -0.01 | -0.01 | -0.01 | -0.01 | -0.01 | -0.01 | -0.01 | -0.01 | -0.01 | -0.01 |
| 39 | -0.01 | -0.01 | -0.01 | -0.01 | -0.01 | -0.01 | -0.01 | -0.01 | -0.01 | 0.025 |
| 40 | -0.01 | -0.01 | -0.01 | -0.01 | -0.01 | -0.01 | -0.01 | -0.01 | -0.01 | 0.025 |
| 41 | -0.01 | -0.01 | -0.01 | -0.01 | -0.01 | -0.01 | -0.01 | -0.01 | -0.01 | -0.01 |
| 42 | -0.01 | -0.01 | -0.01 | -0.01 | -0.01 | -0.01 | -0.01 | -0.01 | -0.01 | -0.01 |
| 43 | -0.01 | -0.01 | 0.025 | 0.025 | -0.01 | 0.025 | 0.025 | -0.01 | 0.025 | -0.01 |
| 44 | -0.01 | -0.01 | 0.025 | 0.025 | -0.01 | 0.025 | 0.025 | -0.01 | 0.025 | -0.01 |
| 45 | -0.01 | -0.01 | -0.01 | -0.01 | -0.01 | -0.01 | -0.01 | -0.01 | -0.01 | -0.01 |
| 46 | -0.01 | -0.01 | -0.01 | -0.01 | -0.01 | -0.01 | -0.01 | -0.01 | -0.01 | -0.01 |
| 47 | -0.01 | -0.01 | -0.01 | -0.01 | -0.01 | -0.01 | -0.01 | -0.01 | -0.01 | -0.01 |
| 48 | -0.01 | -0.01 | -0.01 | -0.01 | -0.01 | -0.01 | -0.01 | -0.01 | -0.01 | -0.01 |
| 49 | -0.01 | -0.01 | -0.01 | -0.01 | -0.01 | -0.01 | -0.01 | -0.01 | -0.01 | -0.01 |

**B3.4 Table: Transition node → PFC weight matrix**

Rows = Source transition nodes. Columns = Target PFC nodes. Continued below.

| | 1 | 2 | 3 | 4 | 5 | 6 | 7 | 8 | 9 | 10 | 11 | 12 | 13 | 14 | 15 | 16 | 17 | 18 | 19 | 20 | 21 | 22 | 23 | 24 | 25 |
|---|---|---|---|---|---|---|---|---|---|---|---|---|---|---|---|---|---|---|---|---|---|---|---|---|---|
| 1 | 2 | 2 | 2 | 2 | 2 | 2 | -0.3 | -0.3 | -0.3 | -0.3 | -0.3 | -0.3 | -0.3 | -0.3 | 2 | 2 | -0.3 | -0.3 | -0.3 | -0.3 | -0.3 | -0.3 | -0.3 | -0.3 | -0.3 |
| 2 | 2 | 2 | 2 | 2 | 2 | 2 | -0.3 | -0.3 | -0.3 | -0.3 | -0.3 | -0.3 | -0.3 | -0.3 | 2 | 2 | -0.3 | -0.3 | -0.3 | -0.3 | -0.3 | -0.3 | -0.3 | -0.3 | -0.3 |
| 3 | 2 | 2 | -0.3 | -0.3 | 2 | 2 | 2 | 2 | -0.3 | -0.3 | -0.3 | -0.3 | -0.3 | -0.3 | 2 | 2 | -0.3 | -0.3 | -0.3 | -0.3 | -0.3 | -0.3 | -0.3 | -0.3 | -0.3 |
| 4 | 2 | 2 | -0.3 | -0.3 | 2 | 2 | 2 | 2 | -0.3 | -0.3 | -0.3 | -0.3 | -0.3 | -0.3 | 2 | 2 | -0.3 | -0.3 | -0.3 | -0.3 | -0.3 | -0.3 | -0.3 | -0.3 | -0.3 |
| 5 | 2 | 2 | -0.3 | -0.3 | 2 | 2 | -0.3 | -0.3 | -0.3 | -0.3 | 2 | 2 | -0.3 | -0.3 | 2 | 2 | -0.3 | -0.3 | -0.3 | -0.3 | -0.3 | -0.3 | -0.3 | -0.3 | -0.3 |
| 6 | 2 | 2 | -0.3 | -0.3 | 2 | 2 | -0.3 | -0.3 | -0.3 | -0.3 | 2 | 2 | -0.3 | -0.3 | 2 | 2 | -0.3 | -0.3 | -0.3 | -0.3 | -0.3 | -0.3 | -0.3 | -0.3 | -0.3 |
| 7 | 2 | 2 | 2 | 2 | -0.3 | -0.3 | -0.3 | -0.3 | 2 | 2 | -0.3 | -0.3 | -0.3 | -0.3 | -0.3 | -0.3 | -0.3 | -0.3 | -0.3 | -0.3 | 2 | 2 | -0.3 | -0.3 | -0.3 |
| 8 | 2 | 2 | 2 | 2 | -0.3 | -0.3 | -0.3 | -0.3 | 2 | 2 | -0.3 | -0.3 | -0.3 | -0.3 | -0.3 | -0.3 | -0.3 | -0.3 | -0.3 | -0.3 | 2 | 2 | -0.3 | -0.3 | -0.3 |
| 9 | 2 | 2 | -0.3 | -0.3 | -0.3 | -0.3 | 2 | 2 | 2 | 2 | -0.3 | -0.3 | -0.3 | -0.3 | -0.3 | -0.3 | -0.3 | -0.3 | -0.3 | -0.3 | 2 | 2 | -0.3 | -0.3 | -0.3 |
| 10 | 2 | 2 | -0.3 | -0.3 | -0.3 | -0.3 | 2 | 2 | 2 | 2 | -0.3 | -0.3 | -0.3 | -0.3 | -0.3 | -0.3 | -0.3 | -0.3 | 2 | 2 | 2 | 2 | -0.3 | -0.3 | -0.3 |
| 11 | 2 | 2 | -0.3 | -0.3 | -0.3 | -0.3 | -0.3 | -0.3 | 2 | 2 | 2 | 2 | -0.3 | -0.3 | -0.3 | -0.3 | -0.3 | -0.3 | -0.3 | -0.3 | 2 | 2 | -0.3 | -0.3 | -0.3 |
| 12 | 2 | 2 | -0.3 | -0.3 | -0.3 | -0.3 | -0.3 | -0.3 | 2 | 2 | 2 | 2 | -0.3 | -0.3 | -0.3 | -0.3 | -0.3 | -0.3 | 2 | 2 | 2 | 2 | -0.3 | -0.3 | -0.3 |
| 13 | 2 | 2 | -0.3 | -0.3 | -0.3 | -0.3 | 2 | 2 | -0.3 | -0.3 | -0.3 | -0.3 | 2 | 2 | -0.3 | -0.3 | 2 | 2 | -0.3 | -0.3 | -0.3 | -0.3 | -0.3 | -0.3 | -0.3 |
| 14 | 2 | 2 | -0.3 | -0.3 | -0.3 | -0.3 | 2 | 2 | -0.3 | -0.3 | -0.3 | -0.3 | 2 | 2 | -0.3 | -0.3 | 2 | 2 | -0.3 | -0.3 | -0.3 | -0.3 | -0.3 | -0.3 | 2 |
| 15 | 2 | 2 | -0.3 | -0.3 | -0.3 | -0.3 | 2 | 2 | -0.3 | -0.3 | -0.3 | -0.3 | 2 | 2 | -0.3 | -0.3 | 2 | 2 | -0.3 | -0.3 | -0.3 | -0.3 | -0.3 | -0.3 | 2 |
| 16 | 2 | 2 | -0.3 | -0.3 | -0.3 | -0.3 | -0.3 | -0.3 | -0.3 | -0.3 | 2 | 2 | 2 | 2 | -0.3 | -0.3 | 2 | 2 | -0.3 | -0.3 | -0.3 | -0.3 | -0.3 | -0.3 | -0.3 |
| 17 | 2 | 2 | -0.3 | -0.3 | -0.3 | -0.3 | -0.3 | -0.3 | -0.3 | -0.3 | 2 | 2 | 2 | 2 | -0.3 | -0.3 | 2 | 2 | -0.3 | -0.3 | -0.3 | -0.3 | -0.3 | -0.3 | 2 |
| 18 | 2 | 2 | -0.3 | -0.3 | -0.3 | -0.3 | -0.3 | -0.3 | -0.3 | -0.3 | 2 | 2 | 2 | 2 | -0.3 | -0.3 | 2 | 2 | -0.3 | -0.3 | -0.3 | -0.3 | 2 | 2 | 2 |
| 19 | 2 | 2 | -0.45 | -0.45 | -0.45 | -0.45 | -0.45 | -0.45 | -0.45 | -0.45 | 3 | 3 | 3 | 3 | -0.45 | -0.45 | 3 | 3 | -0.45 | -0.45 | -0.45 | -0.45 | -0.45 | -0.45 | -0.45 |
| 20 | 2 | 2 | -0.3 | -0.3 | -0.3 | -0.3 | -0.3 | -0.3 | -0.3 | -0.3 | 2 | 2 | 2 | 2 | -0.3 | -0.3 | 2 | 2 | -0.3 | -0.3 | -0.3 | -0.3 | -0.3 | -0.3 | 2 |
| 21 | 2 | 2 | -0.3 | -0.3 | -0.3 | -0.3 | -0.3 | -0.3 | -0.3 | -0.3 | 2 | 2 | 2 | 2 | -0.3 | -0.3 | 2 | 2 | -0.3 | -0.3 | -0.3 | -0.3 | -0.3 | -0.3 | 2 |

**B3.4 Table: Transition node → PFC weight matrix**

| | 26 | 27 | 28 | 29 | 30 | 31 | 32 | 33 | 34 | 35 | 36 | 37 | 38 | 39 | 40 | 41 | 42 | 43 | 44 | 45 | 46 | 47 | 48 | 49 |
|---|---|---|---|---|---|---|---|---|---|---|---|---|---|---|---|---|---|---|---|---|---|---|---|---|
| 1 | -0.3 | -0.3 | -0.3 | 2 | 2 | -0.3 | -0.3 | -0.3 | -0.3 | -0.3 | -0.3 | -0.3 | -0.3 | -0.3 | -0.3 | -0.3 | -0.3 | -0.3 | -0.3 | -0.3 | -0.3 | -0.3 | -0.3 | -0.3 |
| 2 | -0.3 | 2 | 2 | -0.3 | -0.3 | 2 | 2 | -0.3 | -0.3 | -0.3 | -0.3 | -0.3 | -0.3 | -0.3 | -0.3 | -0.3 | -0.3 | -0.3 | -0.3 | -0.3 | -0.3 | -0.3 | -0.3 | -0.3 |
| 3 | -0.3 | -0.3 | -0.3 | 2 | 2 | -0.3 | -0.3 | -0.3 | -0.3 | -0.3 | -0.3 | -0.3 | -0.3 | -0.3 | -0.3 | -0.3 | -0.3 | -0.3 | -0.3 | -0.3 | -0.3 | -0.3 | -0.3 | -0.3 |
| 4 | -0.3 | 2 | 2 | -0.3 | -0.3 | 2 | 2 | -0.3 | -0.3 | -0.3 | -0.3 | -0.3 | -0.3 | -0.3 | -0.3 | -0.3 | -0.3 | -0.3 | -0.3 | -0.3 | -0.3 | -0.3 | -0.3 | -0.3 |
| 5 | -0.3 | -0.3 | -0.3 | 2 | 2 | -0.3 | -0.3 | -0.3 | -0.3 | -0.3 | -0.3 | -0.3 | -0.3 | -0.3 | -0.3 | -0.3 | -0.3 | -0.3 | -0.3 | -0.3 | -0.3 | -0.3 | -0.3 | -0.3 |
| 6 | -0.3 | 2 | 2 | -0.3 | -0.3 | 2 | 2 | -0.3 | -0.3 | -0.3 | -0.3 | -0.3 | -0.3 | -0.3 | -0.3 | -0.3 | -0.3 | -0.3 | -0.3 | -0.3 | -0.3 | -0.3 | -0.3 | -0.3 |
| 7 | -0.3 | -0.3 | -0.3 | 2 | 2 | -0.3 | -0.3 | -0.3 | -0.3 | -0.3 | -0.3 | -0.3 | -0.3 | -0.3 | -0.3 | -0.3 | -0.3 | -0.3 | -0.3 | -0.3 | -0.3 | -0.3 | -0.3 | -0.3 |
| 8 | -0.3 | 2 | 2 | -0.3 | -0.3 | -0.3 | -0.3 | 2 | 2 | -0.3 | -0.3 | -0.3 | -0.3 | -0.3 | -0.3 | -0.3 | -0.3 | -0.3 | -0.3 | -0.3 | -0.3 | -0.3 | -0.3 | -0.3 |
| 9 | -0.3 | -0.3 | -0.3 | 2 | 2 | -0.3 | -0.3 | -0.3 | -0.3 | -0.3 | -0.3 | -0.3 | -0.3 | -0.3 | -0.3 | -0.3 | -0.3 | -0.3 | -0.3 | -0.3 | -0.3 | -0.3 | -0.3 | -0.3 |
| 10 | -0.3 | 2 | 2 | -0.3 | -0.3 | -0.3 | -0.3 | 2 | 2 | -0.3 | -0.3 | -0.3 | -0.3 | -0.3 | -0.3 | -0.3 | -0.3 | -0.3 | -0.3 | -0.3 | -0.3 | -0.3 | -0.3 | -0.3 |
| 11 | -0.3 | -0.3 | -0.3 | 2 | 2 | -0.3 | -0.3 | -0.3 | -0.3 | -0.3 | -0.3 | -0.3 | -0.3 | -0.3 | -0.3 | -0.3 | -0.3 | -0.3 | -0.3 | -0.3 | -0.3 | -0.3 | -0.3 | -0.3 |
| 12 | -0.3 | 2 | 2 | -0.3 | -0.3 | -0.3 | -0.3 | 2 | 2 | -0.3 | -0.3 | -0.3 | -0.3 | -0.3 | -0.3 | -0.3 | -0.3 | -0.3 | -0.3 | -0.3 | -0.3 | -0.3 | -0.3 | -0.3 |
| 13 | -0.3 | -0.3 | -0.3 | 2 | 2 | -0.3 | -0.3 | -0.3 | -0.3 | -0.3 | -0.3 | -0.3 | -0.3 | -0.3 | -0.3 | -0.3 | -0.3 | 2 | 2 | -0.3 | -0.3 | -0.3 | -0.3 | -0.3 |
| 14 | 2 | -0.3 | -0.3 | -0.3 | -0.3 | -0.3 | -0.3 | -0.3 | -0.3 | 2 | 2 | -0.3 | -0.3 | -0.3 | -0.3 | -0.3 | -0.3 | 2 | 2 | -0.3 | -0.3 | -0.3 | -0.3 | -0.3 |
| 15 | 2 | 2 | 2 | -0.3 | -0.3 | -0.3 | -0.3 | 2 | 2 | -0.3 | -0.3 | -0.3 | -0.3 | -0.3 | -0.3 | -0.3 | -0.3 | 2 | 2 | -0.3 | -0.3 | -0.3 | -0.3 | -0.3 |
| 16 | -0.3 | -0.3 | -0.3 | 2 | 3 | -0.3 | -0.3 | -0.3 | -0.3 | -0.3 | -0.3 | -0.3 | -0.3 | -0.3 | -0.3 | -0.3 | -0.3 | 2 | 2 | -0.3 | -0.3 | -0.3 | -0.3 | -0.3 |
| 17 | 2 | -0.3 | -0.3 | -0.3 | -0.3 | -0.3 | -0.3 | -0.3 | -0.3 | 2 | 2 | -0.3 | -0.3 | -0.3 | -0.3 | -0.3 | -0.3 | 2 | 2 | -0.3 | -0.3 | -0.3 | -0.3 | -0.3 |
| 18 | 2 | 2 | 2 | -0.3 | -0.3 | -0.3 | -0.3 | 2 | 2 | -0.3 | -0.3 | -0.3 | -0.3 | -0.3 | -0.3 | -0.3 | -0.3 | 2 | 2 | -0.3 | -0.3 | -0.3 | -0.3 | -0.3 |
| 19 | -0.45 | -0.3 | -0.45 | 3 | 3 | -0.45 | -0.45 | -0.45 | -0.45 | -0.45 | -0.45 | -0.45 | -0.45 | 3 | 3 | -0.45 | -0.45 | -0.45 | -0.45 | -0.45 | -0.45 | -0.45 | -0.45 | -0.45 |
| 20 | 2 | -0.3 | -0.3 | -0.3 | -0.3 | -0.3 | -0.3 | -0.3 | -0.3 | 2 | 2 | -0.3 | -0.3 | 2 | 2 | -0.3 | -0.3 | -0.3 | -0.3 | -0.3 | -0.3 | -0.3 | -0.3 | -0.3 |
| 21 | 2 | 2 | 2 | -0.3 | -0.3 | -0.3 | -0.3 | 2 | 2 | -0.3 | -0.3 | -0.3 | -0.3 | 2 | 2 | -0.3 | -0.3 | -0.3 | -0.3 | -0.3 | -0.3 | -0.3 | -0.3 | -0.3 |

**B3.5 Table : Environment node → transition node weight matrix**

Rows = Source environment nodes.; Columns = Target transition nodes.

Continued on next page.

| | 1 | 2 | 3 | 4 | 5 | 6 | 7 | 8 | 9 | 10 | 11 |
|---|---|---|---|---|---|---|---|---|---|---|---|
| 1 | -0.1 | 0.125 | -0.1 | 0.125 | -0.1 | 0.125 | -0.1 | -0.1 | -0.1 | -0.1 | -0.1 |
| 2 | -0.1 | -0.1 | -0.1 | -0.1 | -0.1 | -0.1 | -0.1 | -0.1 | -0.1 | -0.1 | -0.1 |
| 3 | -0.1 | -0.1 | -0.1 | -0.1 | -0.1 | -0.1 | -0.1 | 0.111 | -0.1 | 0.111 | -0.1 |
| 4 | -0.1 | -0.1 | -0.1 | -0.1 | -0.1 | -0.1 | -0.1 | -0.1 | -0.1 | -0.1 | -0.1 |
| 5 | -0.1 | -0.1 | -0.1 | -0.1 | -0.1 | -0.1 | -0.1 | -0.1 | -0.1 | -0.1 | -0.1 |
| 6 | -0.1 | -0.1 | -0.1 | -0.1 | -0.1 | -0.1 | -0.1 | -0.1 | -0.1 | -0.1 | -0.1 |
| 7 | -0.1 | -0.1 | -0.1 | -0.1 | -0.1 | -0.1 | -0.1 | -0.1 | -0.1 | -0.1 | -0.1 |
| 8 | -0.1 | -0.1 | -0.1 | -0.1 | -0.1 | -0.1 | -0.1 | -0.1 | -0.1 | -0.1 | -0.1 |
| 9 | 0.0714 | -0.05 | 0.0714 | -0.05 | 0.0714 | -0.05 | -0.05 | -0.05 | -0.05 | -0.05 | -0.05 |
| 10 | 0.071 | 0.063 | 0.071 | 0.063 | 0.071 | 0.063 | 0.063 | 0.056 | 0.063 | 0.056 | 0.063 |
| 11 | 0.071 | 0.063 | 0.071 | 0.063 | 0.071 | 0.063 | 0.063 | -0.05 | 0.063 | -0.05 | 0.063 |
| 12 | 0.071 | 0.063 | 0.071 | 0.063 | 0.071 | 0.063 | 0.063 | 0.056 | 0.063 | 0.056 | 0.063 |
| 13 | 0.071 | 0.063 | 0.071 | 0.063 | 0.071 | 0.063 | 0.063 | 0.056 | 0.063 | 0.056 | 0.063 |
| 14 | -0.05 | -0.05 | -0.05 | -0.05 | -0.05 | -0.05 | -0.05 | -0.05 | -0.05 | -0.05 | -0.05 |
| 15 | -0.05 | -0.05 | -0.05 | -0.05 | -0.05 | -0.05 | -0.05 | -0.05 | -0.05 | -0.05 | -0.05 |
| 16 | -0.05 | -0.05 | -0.05 | -0.05 | -0.05 | -0.05 | -0.05 | -0.05 | -0.05 | -0.05 | -0.05 |
| 17 | -0.1 | -0.1 | -0.1 | -0.1 | -0.1 | -0.1 | -0.1 | -0.1 | -0.1 | -0.1 | -0.1 |
| 18 | -0.1 | -0.1 | -0.1 | -0.1 | -0.1 | -0.1 | -0.1 | -0.1 | -0.1 | -0.1 | -0.1 |
| 19 | -0.1 | -0.1 | -0.1 | -0.1 | -0.1 | -0.1 | -0.1 | -0.1 | -0.1 | -0.1 | -0.1 |
| 20 | -0.1 | -0.1 | -0.1 | -0.1 | -0.1 | -0.1 | -0.1 | -0.1 | -0.1 | -0.1 | -0.1 |
| 21 | -0.1 | -0.1 | -0.1 | -0.1 | -0.1 | -0.1 | -0.1 | -0.1 | -0.1 | -0.1 | -0.1 |
| 22 | -0.1 | -0.1 | -0.1 | -0.1 | -0.1 | -0.1 | -0.1 | -0.1 | -0.1 | -0.1 | -0.1 |
| 23 | -0.1 | -0.1 | -0.1 | -0.1 | -0.1 | -0.1 | -0.1 | -0.1 | -0.1 | -0.1 | -0.1 |
| 24 | -0.1 | -0.1 | -0.1 | -0.1 | -0.1 | -0.1 | -0.1 | -0.1 | -0.1 | -0.1 | -0.1 |
| 25 | -0.05 | -0.05 | -0.05 | -0.05 | -0.05 | -0.05 | -0.05 | -0.05 | -0.05 | -0.05 | -0.05 |
| 26 | -0.05 | -0.05 | -0.05 | -0.05 | -0.05 | -0.05 | -0.05 | -0.05 | -0.05 | -0.05 | -0.05 |
| 27 | -0.05 | -0.05 | -0.05 | -0.05 | -0.05 | -0.05 | -0.05 | -0.05 | -0.05 | -0.05 | -0.05 |
| 28 | -0.05 | -0.05 | -0.05 | -0.05 | -0.05 | -0.05 | -0.05 | -0.05 | -0.05 | -0.05 | -0.05 |
| 29 | -0.05 | -0.05 | -0.05 | -0.05 | -0.05 | -0.05 | -0.05 | -0.05 | -0.05 | -0.05 | -0.05 |
| 30 | 0.071 | 0.063 | 0.071 | 0.063 | 0.071 | 0.063 | 0.063 | 0.056 | 0.063 | 0.056 | 0.063 |
| 31 | -0.05 | -0.05 | -0.05 | -0.05 | -0.05 | -0.05 | -0.05 | -0.05 | -0.05 | -0.05 | -0.05 |
| 32 | -0.05 | -0.05 | -0.05 | -0.05 | -0.05 | -0.05 | -0.05 | -0.05 | -0.05 | -0.05 | -0.05 |
| 33 | -0.05 | -0.05 | -0.05 | -0.05 | -0.05 | -0.05 | -0.05 | -0.05 | -0.05 | -0.05 | -0.05 |
| 34 | -0.05 | -0.05 | -0.05 | -0.05 | -0.05 | -0.05 | -0.05 | -0.05 | -0.05 | -0.05 | -0.05 |
| 35 | -0.05 | -0.05 | -0.05 | -0.05 | -0.05 | -0.05 | -0.05 | -0.05 | -0.05 | -0.05 | -0.05 |
| 36 | -0.05 | -0.05 | -0.05 | -0.05 | -0.05 | -0.05 | -0.05 | -0.05 | -0.05 | -0.05 | -0.05 |
| 37 | -0.05 | -0.05 | -0.05 | -0.05 | -0.05 | -0.05 | -0.05 | -0.05 | -0.05 | -0.05 | -0.05 |
| 38 | -0.05 | -0.05 | -0.05 | -0.05 | -0.05 | -0.05 | -0.05 | -0.05 | -0.05 | -0.05 | -0.05 |
| 39 | 0.071 | 0.063 | 0.071 | 0.063 | 0.071 | 0.063 | 0.063 | 0.056 | 0.063 | 0.056 | 0.063 |
| 40 | -0.05 | -0.05 | -0.05 | -0.05 | -0.05 | -0.05 | -0.05 | -0.05 | -0.05 | -0.05 | -0.05 |
| 41 | -0.1 | -0.1 | -0.1 | -0.1 | -0.1 | -0.1 | 0.125 | 0.111 | 0.125 | 0.111 | 0.125 |
| 42 | -0.1 | -0.1 | -0.1 | -0.1 | -0.1 | -0.1 | -0.1 | -0.1 | -0.1 | -0.1 | -0.1 |
| 43 | -0.1 | -0.1 | -0.1 | -0.1 | -0.1 | -0.1 | -0.1 | -0.1 | -0.1 | -0.1 | -0.1 |
| 44 | -0.1 | -0.1 | -0.1 | -0.1 | -0.1 | -0.1 | -0.1 | -0.1 | -0.1 | -0.1 | -0.1 |
| 45 | -0.1 | -0.1 | -0.1 | -0.1 | -0.1 | -0.1 | -0.1 | -0.1 | -0.1 | -0.1 | -0.1 |
| 46 | -0.1 | -0.1 | -0.1 | -0.1 | -0.1 | -0.1 | -0.1 | -0.1 | -0.1 | -0.1 | -0.1 |
| 47 | -0.1 | -0.1 | -0.1 | -0.1 | -0.1 | -0.1 | -0.1 | -0.1 | -0.1 | -0.1 | -0.1 |
| 48 | -0.1 | -0.1 | -0.1 | -0.1 | -0.1 | -0.1 | -0.1 | -0.1 | -0.1 | -0.1 | -0.1 |

**B3.5 Table: Environment node → transition node weight matrix**

Continued from previous page

| | 12 | 13 | 14 | 15 | 16 | 17 | 18 | 19 | 20 | 21 |
|---|---|---|---|---|---|---|---|---|---|---|
| 1 | -0.1 | -0.1 | -0.1 | -0.1 | -0.1 | -0.1 | -0.1 | -0.1 | -0.1 | -0.1 |
| 2 | -0.1 | -0.1 | 0.125 | 0.1 | -0.1 | 0.125 | 0.1 | -0.1 | 0.125 | 0.1 |
| 3 | 0.111 | 0.125 | -0.1 | -0.1 | 0.125 | -0.1 | -0.1 | -0.1 | -0.1 | -0.1 |
| 4 | -0.1 | -0.1 | -0.1 | -0.1 | -0.1 | -0.1 | -0.1 | -0.1 | -0.1 | -0.1 |
| 5 | -0.1 | -0.1 | -0.1 | -0.1 | -0.1 | -0.1 | -0.1 | -0.1 | -0.1 | -0.1 |
| 6 | -0.1 | -0.1 | -0.1 | -0.1 | -0.1 | -0.1 | -0.1 | -0.1 | -0.1 | -0.1 |
| 7 | -0.1 | -0.1 | -0.1 | -0.1 | -0.1 | -0.1 | -0.1 | -0.1 | -0.1 | -0.1 |
| 8 | -0.1 | -0.1 | -0.1 | -0.1 | -0.1 | -0.1 | -0.1 | -0.1 | -0.1 | -0.1 |
| 9 | -0.05 | -0.05 | -0.05 | -0.05 | -0.05 | -0.05 | -0.05 | -0.05 | -0.05 | -0.05 |
| 10 | 0.056 | 0.063 | -0.05 | -0.05 | 0.063 | -0.05 | -0.05 | -0.05 | -0.05 | -0.05 |
| 11 | -0.05 | -0.05 | 0.063 | 0.05 | -0.05 | 0.063 | 0.05 | -0.05 | 0.063 | 0.05 |
| 12 | 0.056 | 0.063 | 0.063 | 0.05 | 0.063 | 0.063 | 0.05 | -0.05 | 0.063 | 0.05 |
| 13 | 0.056 | 0.063 | 0.063 | 0.05 | 0.063 | 0.063 | 0.05 | -0.05 | 0.063 | 0.05 |
| 14 | -0.05 | -0.05 | -0.05 | -0.05 | -0.05 | -0.05 | -0.05 | -0.05 | -0.05 | -0.05 |
| 15 | -0.05 | -0.05 | -0.05 | -0.05 | -0.05 | -0.05 | -0.05 | -0.05 | -0.05 | -0.05 |
| 16 | -0.05 | -0.05 | -0.05 | -0.05 | -0.05 | -0.05 | -0.05 | -0.05 | -0.05 | -0.05 |
| 17 | -0.1 | -0.1 | -0.1 | -0.1 | -0.1 | -0.1 | -0.1 | -0.1 | -0.1 | -0.1 |
| 18 | -0.1 | -0.1 | -0.1 | -0.1 | -0.1 | -0.1 | -0.1 | -0.1 | -0.1 | -0.1 |
| 19 | -0.1 | -0.1 | -0.1 | -0.1 | -0.1 | -0.1 | -0.1 | -0.1 | -0.1 | -0.1 |
| 20 | -0.1 | -0.1 | -0.1 | -0.1 | -0.1 | -0.1 | -0.1 | -0.1 | -0.1 | -0.1 |
| 21 | -0.1 | -0.1 | -0.1 | -0.1 | -0.1 | -0.1 | -0.1 | -0.1 | -0.1 | -0.1 |
| 22 | -0.1 | -0.1 | -0.1 | -0.1 | -0.1 | -0.1 | -0.1 | -0.1 | -0.1 | -0.1 |
| 23 | -0.1 | -0.1 | -0.1 | 0.1 | -0.1 | -0.1 | 0.1 | -0.1 | -0.1 | 0.1 |
| 24 | -0.1 | -0.1 | -0.1 | -0.1 | -0.1 | -0.1 | -0.1 | -0.1 | -0.1 | -0.1 |
| 25 | -0.05 | -0.05 | -0.05 | -0.05 | -0.05 | -0.05 | -0.05 | -0.05 | -0.05 | -0.05 |
| 26 | -0.05 | -0.05 | -0.05 | -0.05 | -0.05 | -0.05 | -0.05 | -0.05 | -0.05 | -0.05 |
| 27 | -0.05 | -0.05 | -0.05 | -0.05 | -0.05 | -0.05 | -0.05 | -0.05 | -0.05 | -0.05 |
| 28 | -0.05 | -0.05 | -0.05 | -0.05 | -0.05 | -0.05 | -0.05 | -0.05 | -0.05 | -0.05 |
| 29 | -0.05 | -0.05 | -0.05 | -0.05 | -0.05 | -0.05 | -0.05 | -0.05 | -0.05 | -0.05 |
| 30 | 0.056 | -0.05 | -0.05 | -0.05 | -0.05 | -0.05 | -0.05 | -0.05 | -0.05 | -0.05 |
| 31 | -0.05 | -0.05 | -0.05 | -0.05 | -0.05 | -0.05 | -0.05 | -0.05 | -0.05 | -0.05 |
| 32 | -0.05 | -0.05 | -0.05 | -0.05 | -0.05 | -0.05 | -0.05 | -0.05 | -0.05 | -0.05 |
| 33 | -0.05 | -0.05 | -0.05 | -0.05 | -0.05 | -0.05 | -0.05 | -0.05 | -0.05 | -0.05 |
| 34 | -0.05 | -0.05 | -0.05 | -0.05 | -0.05 | -0.05 | -0.05 | -0.05 | -0.05 | -0.05 |
| 35 | -0.05 | -0.05 | -0.05 | -0.05 | -0.05 | -0.05 | -0.05 | -0.05 | -0.05 | -0.05 |
| 36 | -0.05 | -0.05 | -0.05 | -0.05 | -0.05 | -0.05 | -0.05 | -0.05 | -0.05 | -0.05 |
| 37 | -0.05 | -0.05 | -0.05 | -0.05 | -0.05 | -0.05 | -0.05 | -0.05 | -0.05 | -0.05 |
| 38 | -0.05 | -0.05 | -0.05 | -0.05 | -0.05 | -0.05 | -0.05 | -0.05 | -0.05 | -0.05 |
| 39 | 0.056 | 0.063 | 0.063 | 0.05 | 0.063 | 0.063 | 0.05 | -0.05 | 0.063 | 0.05 |
| 40 | -0.05 | -0.05 | -0.05 | -0.05 | -0.05 | -0.05 | -0.05 | -0.05 | -0.05 | -0.05 |
| 41 | 0.111 | -0.1 | -0.1 | -0.1 | -0.1 | -0.1 | -0.1 | -0.1 | -0.1 | -0.1 |
| 42 | -0.1 | -0.1 | -0.1 | -0.1 | -0.1 | -0.1 | -0.1 | -0.1 | -0.1 | -0.1 |
| 43 | -0.1 | -0.1 | -0.1 | -0.1 | -0.1 | -0.1 | -0.1 | -0.1 | -0.1 | -0.1 |
| 44 | -0.1 | -0.1 | -0.1 | -0.1 | -0.1 | -0.1 | -0.1 | -0.1 | -0.1 | -0.1 |
| 45 | -0.1 | -0.1 | -0.1 | -0.1 | -0.1 | -0.1 | -0.1 | -0.1 | -0.1 | -0.1 |
| 46 | -0.1 | -0.1 | -0.1 | -0.1 | -0.1 | -0.1 | -0.1 | -0.1 | -0.1 | -0.1 |
| 47 | -0.1 | -0.1 | -0.1 | -0.1 | -0.1 | -0.1 | -0.1 | -0.1 | -0.1 | -0.1 |
| 48 | -0.1 | 0.125 | 0.125 | 0.1 | 0.125 | 0.125 | 0.1 | -0.1 | 0.125 | 0.1 |

**B3.6 Table: PFC → Object representations weight matrix**

Rows = Source PFC channels; Columns = Target object nodes.

| | 1 | 2 | 3 | 4 | 5 |
|---|---|---|---|---|---|
| 1 | 0 | 0 | 0 | 0 | 0 |
| 2 | 0 | 0 | 0 | 0 | 0 |
| 3 | 0 | 0 | 0 | 0 | 0 |
| 4 | 0 | 0 | 0 | 0 | 0 |
| 5 | 0 | 0 | 0 | 0 | 0 |
| 6 | 0 | 0 | 0 | 0 | 0 |
| 7 | 0 | 0 | 0 | 0 | 0 |
| 8 | 0 | 0 | 0 | 0 | 0 |
| 9 | 0 | 0 | 0 | 0 | 0 |
| 10 | 0 | 0 | 0 | 0 | 0 |
| 11 | 0 | 0 | 0 | 0 | 0 |
| 12 | 0 | 0 | 0 | 0 | 0 |
| 13 | 0 | 0 | 0 | 0 | 0 |
| 14 | 0 | 0 | 0 | 0 | 0 |
| 15 | 0.3 | 0 | 0 | 0 | 0 |
| 16 | 0.3 | 0 | 0 | 0 | 0 |
| 17 | 0 | 0.3 | 0 | 0 | 0 |
| 18 | 0 | 0.3 | 0 | 0 | 0 |
| 19 | 0 | 0 | 0 | 0 | 0 |
| 20 | 0 | 0 | 0 | 0 | 0 |
| 21 | 0 | 0 | 0.3 | 0 | 0 |
| 22 | 0 | 0 | 0.3 | 0 | 0 |
| 23 | 0 | 0 | 0 | 0 | 0 |
| 24 | 0 | 0 | 0 | 0 | 0 |
| 25 | 0 | 0 | 0 | 0.3 | 0 |

| | 1 | 2 | 3 | 4 | 5 |
|---|---|---|---|---|---|
| 26 | 0 | 0 | 0 | 0.3 | 0 |
| 27 | 0 | 0 | 0 | 0 | 0.3 |
| 28 | 0 | 0 | 0 | 0 | 0.3 |
| 29 | 0.3 | 0.3 | 0.3 | 0 | 0 |
| 30 | 0.3 | 0.3 | 0.3 | 0 | 0 |
| 31 | 0 | 0 | 0 | 0 | 0.3 |
| 32 | 0 | 0 | 0 | 0 | 0.3 |
| 33 | 0 | 0 | 0 | 0 | 0.3 |
| 34 | 0 | 0 | 0 | 0 | 0.3 |
| 35 | 0 | 0 | 0 | 0.3 | 0 |
| 36 | 0 | 0 | 0 | 0.3 | 0 |
| 37 | 0 | 0 | 0 | 0 | 0 |
| 38 | 0 | 0 | 0 | 0 | 0 |
| 39 | 0 | 0 | 0 | 0 | 0 |
| 40 | 0 | 0 | 0 | 0 | 0 |
| 41 | 0 | 0 | 0 | 0 | 0 |
| 42 | 0 | 0 | 0 | 0 | 0 |
| 43 | 0 | 0 | 0 | 0 | 0 |
| 44 | 0 | 0 | 0 | 0 | 0 |
| 45 | 0 | 0 | 0 | 0 | 0 |
| 46 | 0 | 0 | 0 | 0 | 0 |
| 47 | 0 | 0 | 0 | 0 | 0 |
| 48 | 0 | 0 | 0 | 0 | 0 |
| 49 | 0 | 0 | 0 | 0 | 0 |
